# EMULSION: transparent and flexible multiscale stochastic models in human, animal and plant epidemiology

**DOI:** 10.1101/563791

**Authors:** Sébastien Picault, Yu-Lin Huang, Vianney Sicard, Sandie Arnoux, Gaël Beaunée, Pauline Ezanno

**Affiliations:** BIOEPAR, INRA, Oniris, CS40706, 44307 Nantes, France; Univ. Lille, CNRS, Centrale Lille, UMR 9189 - CRIStAL, F-59000 Lille, France

## Abstract

Stochastic mechanistic epidemiological models largely contribute to better understand pathogen emergence and spread, and assess control strategies at various scales (from within-host to transnational scale). However, developing realistic models which involve multi-disciplinary knowledge integration faces three major challenges in predictive epidemiology: lack of readability once translated into simulation code, low reproducibility and reusability, and long development time compared to outbreak time scale. We introduce here EMULSION, an artificial intelligence-based software intended to address those issues and help modellers focus on model design rather than programming. EMULSION defines a domain-specific language to make all components of an epidemiological model (structure, processes, parameters…) explicit as a structured text file. This file is readable by scientists from other fields (epidemiologists, biologists, economists), who can contribute to validate or revise assumptions at any stage of model development. It is then automatically processed by EMULSION generic simulation engine, preventing any discrepancy between model description and implementation. The modelling language and simulation architecture both rely on the combination of advanced artificial intelligence methods (knowledge representation and multi-level agent-based simulation), allowing several modelling paradigms (from compartment- to individual-based models) at several scales (up to metapopulation). The flexibility of EMULSION and its capability to support iterative modelling are illustrated here through examples of progressive complexity, including late revisions of core model assumptions. EMULSION is also currently used to model the spread of several diseases in real pathosystems. EMULSION provides a command-line tool for checking models, producing model diagrams, running simulations, and plotting outputs. Written in Python 3, EMULSION runs on Linux, MacOS, and Windows. It is released under Apache-2.0 license. A comprehensive documentation with installation instructions, a tutorial and many examples are available from: https://sourcesup.renater.fr/www/emulsion-public.

## Introduction

Understanding and predicting the spread of pathogens at several scales (from individuals to territories) under various scenarios (control measures, climate, etc.) relies on realistic mechanistic models [1–5]. Yet, predictive epidemiology is currently facing key methodological challenges which could be faced mobilizing Artificial Intelligence (AI) technologies. First, the lack of model readability often hinders the capability of scientists from life sciences to discuss or revise model assumptions, once translated into simulation code, impairing the users’ confidence as well as critical mind towards model predictions. Second, epidemiological models highly depend on implementation details, which affects reusability in even slightly different contexts. Third, developing mechanistic models for complex systems is an iterative process, rendering it difficult to get reliable forecasts in a duration relevant with regards to on-field needs (e.g. while fighting outbreaks). Fourth, the complexity of realistic models often leads to misinterpretations coming from programming biases [6], thus it is difficult to ensure that model assumptions are correctly implemented in the simulation code. Fifth, model granularity is likely to change throughout the model development process, either to account for increased detail level, or conversely to keep only relevant features at a broader scale. Changes in core model assumptions can also occur late in the modelling process, for instance because a feature first considered negligible must finally be accounted for. This requires a capability to move back and forth between modelling paradigms (stochastic vs. deterministic, compartments vs. individuals) and between scales (individuals, groups, populations, metapopulations) without having to recode large parts of the models.

To address such challenges, the implementation of epidemiological models must first gain transparency, in order to make all assumptions as explicit as possible. Models are a deliberately simplified representation of reality, thus it is crucial to present the underlying assumptions and their implications, to help understand their benefits and limitations and assess their relevance. Once models are translated into simulation code (which is the usual process), scientists without programming skills, involved either in the co-design of the model or in a peer-reviewing process, cannot develop a thorough understanding of the models nor make an informed judgment. Instead, models should be described in a readable form. The ODD protocol (Overview, Design concepts, Details) [7] is a substantial effort to obtain a comprehensive description of model components within a textual template; however, “the lack of real specification” [8] prevents to translate ODD models straight into simulation code. On the other hand, enhancing reusability requires reproducibility (the capability for others to recode a model) and flexibility (the possibility to adapt it). Though necessary, best practices in software engineering [9] cannot settle those issues alone: ad hoc codes can be documented, tested, reproducible, and yet wrong [10].

A first step to reduce development time and errors is to use simulation platforms, which provide high-level built-in features. In general-purpose platforms such as GAMA [11], NetLogo [12] or Repast [13], a substantial programming effort is still required as algorithms involved in epidemiological models are not provided as platform features. Simulation libraries and platforms dedicated to epidemiological issues are rising, e.g. SimInf [14], a R library for data-driven compartment-based models; MicroSim [15], an agent-based platform for several kinds of diseases; or GLEaMviz [16], a metapopulation-oriented platform. To our knowledge, the most advanced approach in terms of diversity of modelling paradigms is Broadwick [17], a Java framework for compartment- and individual-based models with interaction networks, which nevertheless still requires writing large portions of code to derive specific classes and carry out simulations on practical cases.

Artificial intelligence (AI) can help going further, as demonstrated in a promising approach, KENDRICK [18], which defines a domain-specific language (DSL, [19]), which allows to describe models as text files rather than executable code, enforces a clear separation of concerns (infection, demography, etc.) to facilitate model assessment by scientists from several domains, and is used to generate an optimized simulation code, which guarantees that model features are translated without bias into the program. Yet, KENDRICK is designed for compartment-based models only.

To the best of our knowledge, none of existing solutions address those methodological challenges simultaneously, the most advanced approaches providing either a flexibility in modelling paradigms at the expense of software development efforts, or an enhanced readability through a DSL limited to a specific modelling paradigm.

This article introduces EMULSION, an AI-based framework which tackles those challenges in an integrated approach. EMULSION intends to help modellers develop mechanistic stochastic models of complex systems in epidemiology at several scales using multiple paradigms, and to facilitate the coconstruction and assessment of model components (biological assumptions, model structure, parameters, scenarios, etc.) with epidemiologists or scientists from other relevant fields, throughout model development. This makes our software an outstanding contribution towards reliable, reactive and transparent predictive epidemiology.

## Design and implementation

To address the challenges presented above, EMULSION combines two AI methods [20]. The first one is a DSL designed for the description of all components of an epidemiological model, to make them explicit in a human-readable form as a structured text file, so that scientists from different fields can better interact with modellers throughout the modelling process, and discuss, assess or revise model structure, assumptions, parameters at any moment without having to read or write any line of simulation code. The second one is the use of a generic simulation engine, whose core architecture relies upon a multi-level agent-based system [21]. This allows several scales (individuals, groups, populations, metapopulations) and modelling paradigms (compartment- or individual-based models) to be encompassed within a homogeneous software interface, as agents act as wrappers which can be dynamically combined regardless of what they have to compute and the scale at which they operate. To run an experiment, the simulation engine reads the DSL file containing the model description, assembles the agents required for a particular type of model and a specific scale, initializes parameters, functions, processes specified in the DSL file, and make them interact to produce simulation outputs (Fig. 1).

**Fig 1.**
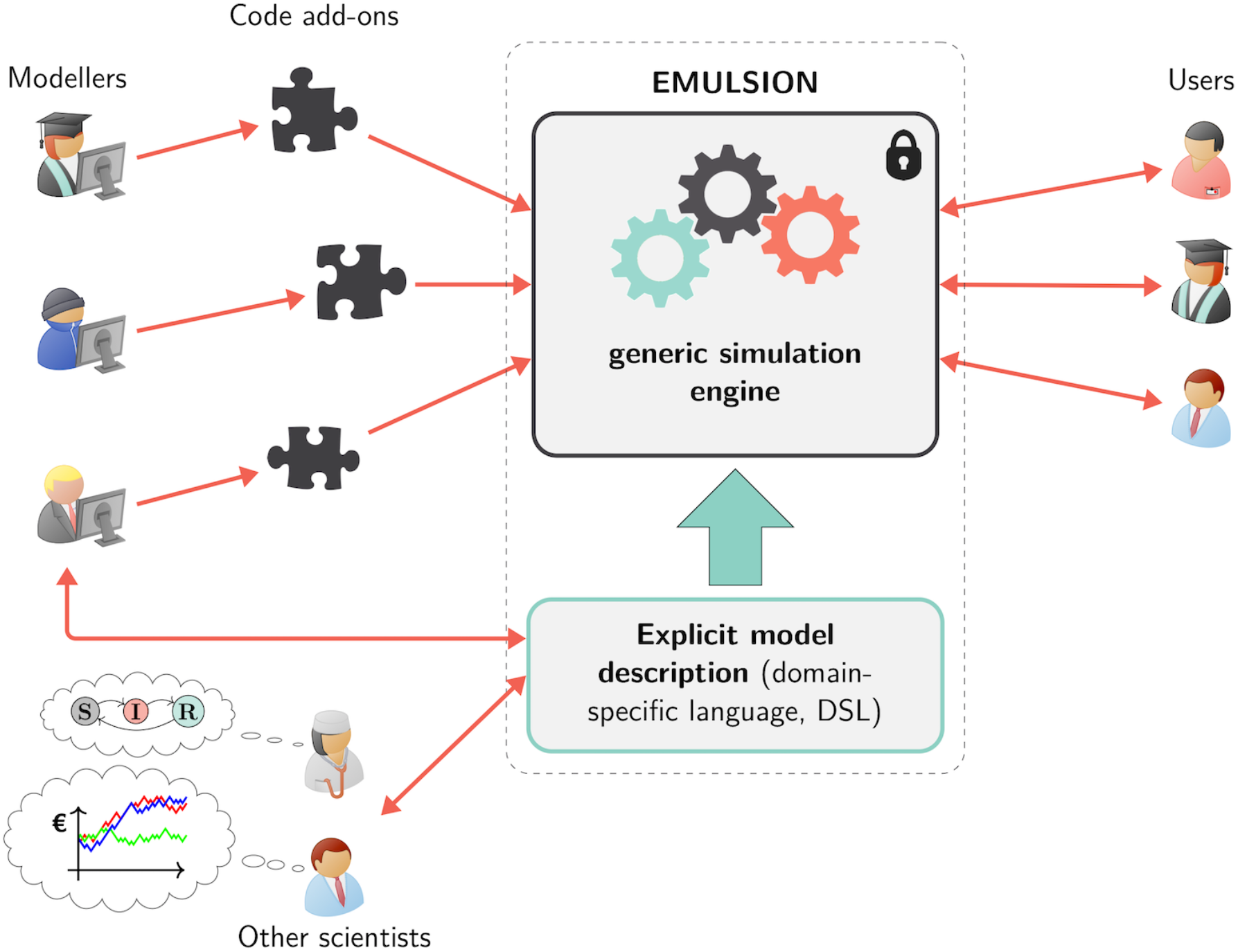
Approach enforced in the EMULSION framework. A generic simulation engine is coupled to a domain-specific modelling language (DSL), reinforcing interactions between modellers and scientists from other fields. Knowledge involved in epidemiological models is kept explicit, understandable and revisable as a structured text file. A few specific software add-ons can be written to complement the simulation engine if needed.

### Representing processes: from flow diagrams to state machines

Processes occurring in the pathosystem (infection, demography, migrations…) are a core component of epidemiological models, often described by flow diagrams [22,23] with nodes denoting state variables (amount of individuals in each state), and transitions labelled with rates. Though flow diagrams can be easily derived into ordinary differential equations or into stochastic difference equations, they often mix several concerns in a unique, monolithic representation, involve implicit computational assumptions (e.g. an exponential distribution of durations in states), and cannot capture additional information (e.g. conditions, actions) which have to be introduced later at implementation. EMULSION relies upon a formalism close to flow diagrams but more accurate: finite state machines [24], classically used in computer science.

Compared to flow diagrams, state machines describe the evolution of one individual instead of a population, and one state machine represents one single process, so that a complex flow diagram may be split into several simpler state machines. Besides, states can be endowed with additional properties, such as a duration distribution specifying how long an individual is expected to stay in the current state, and actions performed by individuals when entering, being in, or leaving the state. Transitions are labelled with either a rate, a probability or an absolute amount (rates are automatically converted into probabilities). They can also specify: 1) calendar conditions to indicate time periods when transitions are available; 2) escape conditions allowing to free from state duration constraints; 3) individual conditions to filter which agents are allowed to cross the transition; 4) actions performed by individuals crossing the transition, i.e. after leaving their current state and before entering their new one (Fig 2).

**Fig 2.**
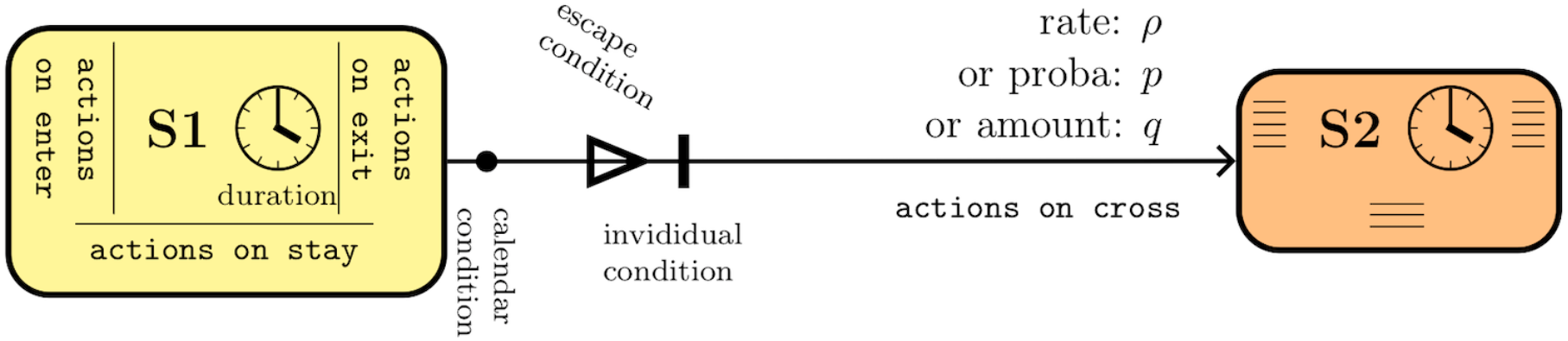
Structure of a transition between two states in state machines. Individuals can be given a duration and actions when entering, staying in, or leaving the state. Transitions feature a rate, probability, or amount, and can be associated with actions performed on crossing, time-dependent (“calendar”) conditions, or individual conditions restricting the capability to cross the transition, and escape conditions allowing individuals to leave their state before the nominal duration.

### A multi-level agent-based simulation architecture to encompass paradigms and scales

Multi-agent systems are composed of autonomous entities (agents) endowed with a behaviour and interacting in a shared environment. In the last decade, multi-level agent-based systems emerged using agents to explicitly represent intermediary abstraction levels (groups, sub-populations, organizations.) with behaviours of their own, between individuals and the whole system [25–28]. Recent advances in this field [21] led to design patterns, i.e. systematic solutions for recurrent modelling issues. Those patterns were used in EMULSION to build the architecture of the generic simulation engine, in which nested agents are in charge of implementing a specific modelling paradigm at a given scale. Agents currently defined in EMULSION allow to implement the main paradigms used in epidemiological models: 1) compartment-based models [29], where state variables represent aggregate amounts of individuals which only differ by few key variables, such as health state or age group; and 2) individual-based models [30] necessary for finer grained representations. Besides, EMULSION provides a hybrid approach which combines the capability of representing detailed information through individuals, with an adaptive grouping of individuals based on their state, to optimize computation (Fig 3, and S1 Appendix, Table 1). The same approach allows to wrap different scales within agents to build either groups, populations, or metapopulations [31,32] which can handle region-wide models at a moderate computational cost (focusing only on relevant populations endowed with a contact structure). Scales and paradigms can be chosen independently: hence, a metapopulation at regional scale can rely at the local scale upon compartment-based, individual-based or hybrid models, depending on the required detail level. In this modular architecture, simulations are run in discrete time to better cope with the potential complexity of interactions between agents from all scales.

**Fig 3.**
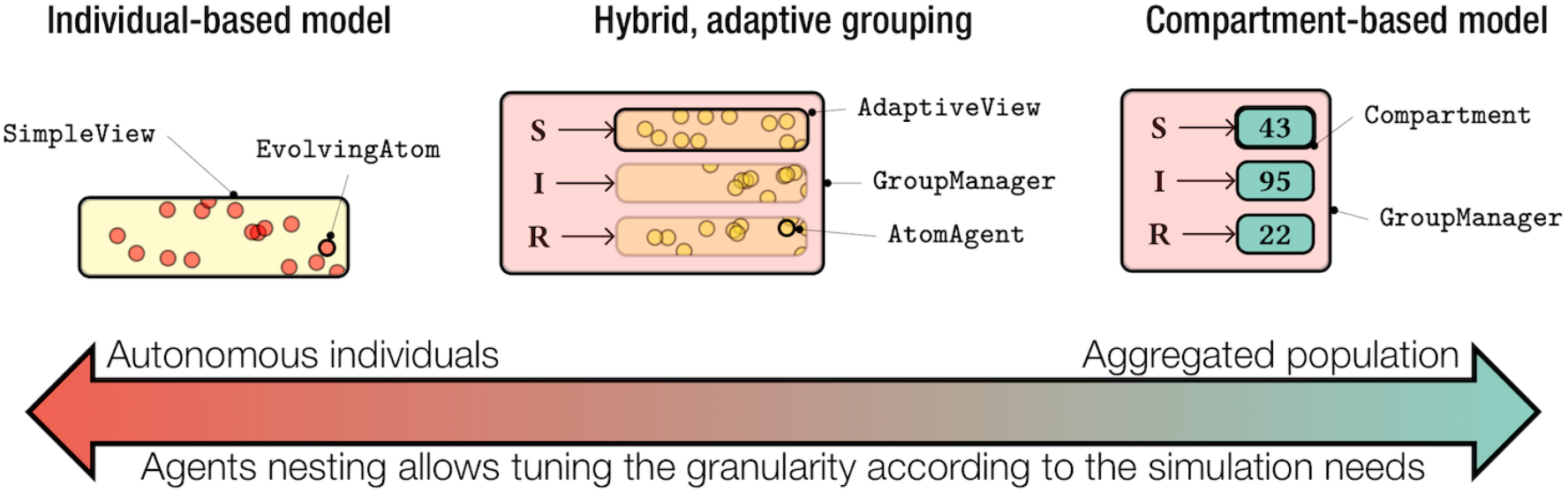
Diversity of modelling paradigms in EMULSION models. EMULSION allows to represent within a same formalism (nested agents) several modelling paradigms, from the finest grained (individuals) to aggregations (compartments), including intermediary representations as a trade-off between computation time and preservation of individual details. The chosen modelling paradigm is associated with the appropriate combinations of agents.

### A domain-specific language for epidemiological models

The main benefits of the DSL defined in EMULSION are a modular decomposition of the model, which reduces dependencies between model components and facilitates further extensions, and a high readability by non-modeller scientists. When EMULSION simulation engine reads the DSL, it selects the agent classes corresponding to the chosen paradigm and scales, instantiate agents depending on initial conditions, transforms expressions into Python functions (using SymPy, a library for symbolic computation), builds the state machines, and runs the simulations. Hence, the execution of simulations based on an EMULSION model is univocal, all required information being directly usable (yet human-readable) by the simulation engine.

EMULSION comes with several built-in actions (changing the values of variables, creating new individuals, etc.) which can be used in states and transitions. When the user defines levels, groups, or states, EMULSION automatically provides associated variables (e.g. population counts, individual tests for each state) to make model description as self-sufficient as possible in model files. In very specific cases, processes or actions may require the definition of small code add-ons, as demonstrated below for data-driven events in a metapopulation. But in the majority of situations, no additional code is required.

An EMULSION model is a text file based on the YAML format [33], which is mainly based on nested lists or key-value pairs, ending with strings, numbers or booleans (yes/no). Each model is composed of several first-level keys (called “sections” in what follows) to provide information by topic: model name, modelling paradigms and scales, processes occurring at each level, structure of state machines, list of parameters and expressions, individual actions, initial conditions, outputs. Section and subsection names are chosen to be as self-evident as possible, and a substantial part of the documentation is dedicated to explaining the syntax of the DSL in detail, with comprehensive examples and tutorials.

EMULSION models can be built by adapting existing examples (changing only a few sections) rather than starting from scratch. Especially, the main part of a model consists in section “state_machines”, which defines the state machines underlying the processes occurring in the pathosystem, and section “parameters”, which provides a comprehensive, commented list of parameters, distributions and functions used in the calculations. Almost all items have to be accompanied by a textual description of their meaning and role. A full example of SIR model implemented with hybrid modelling paradigm is provided in S1 Text. Also, in S1 Appendix, the three modelling paradigms provided by EMULSION (compartment, hybrid and individual-based) are compared on two SIR models (one with constant population, one with birth and death processes): differences in model files are presented side-by-side to highlight how to transform one paradigm into the other; simulation outcomes are shown for the three paradigms in each model implemented with EMULSION and for the equivalent compartment-based model implemented with the R library SimInf [14].

## Results

### Incremental design of complex epidemiological models

EMULSION enables to build non-trivial models incrementally. We illustrate this below with an example from within-population to between-population scales, and demonstrate how easy it is to operate late changes in core model assumptions.

First, a classical SIR model is developed (S: susceptible, I: infectious, R: resistant), assuming a frequency-dependent force of infection. In this example, we use the hybrid modelling paradigm, where individuals are grouped according to similar health states. Transitions (infection and recovery) are driven by a state machine (“health_state”) which is associated with a way of grouping individuals (all individuals of similar health states are grouped). The corresponding YAML file with full description is provided in S1 Text.

Second, we introduce demography and two age groups: juveniles (J) and adults (A). We assume that juveniles become adults at rate “maturation”, that adults produce juveniles at rate “birth” and that both die at rate “mortality”. An additional process is required in the model, hence a new state machine (called “age_group”), also associated to a grouping. The differences introduced compared to the first case are highlighted in S2 Text.

Third, the population model is upscaled at the metapopulation scale, assuming that pathogens spread among populations due to a contact network, which is described by observed data. This part involves two code add-ons: one for reading a CSV file which gives the dates, age groups, quantity, source and destination for each move, another to perform the relevant migrations at each time step. The differences introduced in the previous model are highlighted in S3 Text, and Python code add-ons required to handle data are provided in S4 Text.

After being used to better understand pathogens spread in a population, epidemiological models are often extended to assess control strategies. Let now assume that the user wants to assess the effect of targeted vaccination, e.g. of juveniles in spring. This obviously involves substantial changes in the infection process, to incorporate a vaccinated state and associated transitions. The modularity of EMULSION allows modellers to change the “health_state” state machine without any impact on the metapopulation structure. First, a new health state “vaccinated” (V) must be added. A calendar is introduced in the “time_info” section to define when “spring” occurs. Then, the transition from “S” to “V” has to specify that it can only be used during spring (“when: spring”) and that it concerns only juveniles (“cond: is_J”). The differences introduced in the previous model are highlighted in S5 Text.

### Application to real diseases

EMULSION is currently used to understand pathogen spread within structured populations and metapopulations, and to assess control strategies. A first study was conducted on Q fever, a zoonosis caused by the bacteria *Coxiella burnetii* and affecting mainly domestic ruminants. To better understand the interplay between within- and between-herd levels in bovine dairy herds, and identify the contributions of transmission pathways at regional scale, two existing models (respectively a within- [34] and a between-herd [35] model) were re-implemented using EMULSION. Assumptions of the within-herd model were simplified, keeping relevant assumptions and removing those less crucial in the perspective of the between-herd dynamics [36], which was easily tackled due to EMULSION modularity. EMULSION was also used to assess strategies for the control of Bovine Respiratory Diseases in young bulls at fattening. In a perspective to reduce antibiotic use, metaphylaxis (collective treatment triggered after first cases are detected) was compared to early detection methods, based on sensors measuring hyperthermia durations. Three tightly coupled processes were modelled: infection, hyperthermia, and treatment, and scenarios combining metaphylaxis policies with detection methods were assessed with regards to reduction of antibiotics doses, infection duration, and false positives [37].

Being developed at INRA (French National Institute for Agricultural Research), real disease applications mainly come from veterinary epidemiology, but human and plant diseases can also be addressed. As an example, three published non trivial models were reimplemented with EMULSION (S2 Appendix), addressing real diseases in each field and tackling different issues: the multi-year seasonality of measles outbreaks [22], the spread of a vector-borne zoonosis, the Rift Valley fever [38], and the spatial spread and control of Bahia Bark Scaling of Citrus [39]. Beside reproducing published results in human, animal and plant epidemiology, this shows how EMULSION DSL can help represent complex modelling features in a flexible and explicit way, and how to extend EMULSION through a Python code add-on to incorporate functionalities which are not provided by EMULSION generic simulation engine (here, a dispersal kernel for the spatially explicit model).

### Addressing challenges in disease transmission modelling

The DSL and architecture of EMULSION, both modular and flexible, provide powerful methods to address several challenges in epidemiological modelling [40–44]. First, model transparency, through the DSL, helps engage dialogue across disciplines and communicate with stakeholders and health managers, also contributing to a collaborative design and a long-term maintenance of models. The DSL, making all model components explicit, also fosters comparison and simplification, to address complex systems with relevant models as simple as possible. Second, the separation of concerns allows modellers to design processes (infection, demography, detection, control, etc.) and scales (individuals, populations, metapopulations) independently and at the relevant detail level, which helps handle multiple hosts, pathogen diversity, and realistic detection methods and control measures. Third, multi-scale and multi paradigm modelling capabilities natively provided by EMULSION help model within-host heterogeneity and unify multiple scales of transmission.

### Availability and future directions

The EMULSION package documentation, with installation instructions (either from PyPI or by cloning the GIT source repository), tutorial and examples, can be found on SourceSup (a forge dedicated to French public research institutes) at: https://sourcesup.renater.fr/www/emulsion-public. The package comes with 32 examples to illustrate features of EMULSION modelling language, from very simple SIR models declined in compartmental, individual-based, or hybrid approaches, to epidemiological models in structured populations mixing infectious process, demography, seasonality, forcing external variables, and non-exponential distributions of state durations. Examples also demonstrate how to move from within- to between-population models using data-driven movements. Updates will be released regularly based on the integration on new modelling features or code optimizations.

As demonstrated, EMULSION is flexible enough to cover a broad diversity of stochastic and mechanistic epidemiological models at several scales. Current version of the software is fully usable by modellers and currently deployed to develop models for real diseases assuming various propagation pathways (direct contacts, environment contamination, airborne or vector-borne transmission, vertical transmission), demographic and herd management processes, detection methods (observation of clinical signs, sensors, tests), and control measures (vaccination, antibiotics, tests at purchase, selective culling). Specific features in those epidemiological models, requiring for now dedicated add-ons, will be incorporated to the DSL and to the simulation engine as generic modules if recurrent enough among models, to help encompass a broad range of mechanisms within EMULSION.

We identified three priorities for future work to be carried out in the short term. First, we aim at better automating data integration. Models rely on observed commercial, meteorological, and epidemiological data, either to define parameter values or to confront model predictions with observations. The DSL should thus allow to specify how to inform parameter values from data and how to define data-driven processes. Second, we will address spatially explicit models. For now, discrete metapopulations are possible, relevant for network contact representation but less for proximity contacts (airborne transmission, neighbourhood contacts, vector-borne transmission, etc.). Third, we plan to allow a model to combine several modelling paradigms, e.g. using compartments for vectors and individuals for hosts in a vector-borne disease model. In the longer term, we plan to use the DSL for optimized code generation (e.g. in C/C++). The choice of Python for implementing EMULSION simulation engine was well adapted to facilitate the modular and incremental development of the software, and its portability on various operating systems and environments. Genericity was also achieved through introspection capabilities of the language, but which came with an additional computational cost. This constitutes a limit of the software when tackling multi-scale models with numerous components (e.g. large number of interacting populations), models which analysis is based mainly on massive simulations. Such models written using EMULSION’s DSL then would benefit from being automatically translated into simulation codes generated from templates, allowing for higher computational performances.

The generic modelling approach we promote in EMULSION both has advantages and drawbacks. The gathering of various model features within a homogeneous software tool, which can be picked up and combined according to the modeller’s needs with little or even no additional code, reduces development time, ensures that model assumptions and processes are well implemented, and allows optimized code generation as a mid-term perspective. As a common software limitation, potential bugs in the generic engine could impact all models developed with EMULSION: but its diffusion as an open-source software among a broad community of users is the best way to get feedback on potential malfunctions, identify their origin and correct them rapidly.

To our knowledge, EMULSION is currently the only software which addresses all major computing issues in the design of mechanistic models in epidemiology. It helps modellers focus on model assumptions and structure rather than coding, ensures more reliable models, provides classical modelling paradigms at multiple scales, and makes models explicit and readable, so that they can be assessed by other scientists and shared in reproducible publications. As a result, the whole model design process is accelerated, which substantially enhances the reactiveness of epidemiologists against emerging outbreaks.

## Supporting information

**S1 Text. EMULSION model file for a simple SIR model with hybrid modelling paradigm.** This file (“hybrid_SIR.yaml”) contains all model sections required to describe a classical stochastic SIR model. The hybrid modelling paradigm mixes an individual-based approach (individuals are created in the model) and the compartment-based approach (individuals are grouped automatically depending on their similarities), which renders the hybrid model performance almost as high as for a compartment-based model while enabling individual features.

**S1 Appendix. Comparison of simple SIR models (with constant population and with birth/death processes) implemented with compartments in R using library SimInf, and in EMULSION with compartment-based, hybrid and individual-based modelling paradigms.** For each model, simulation outcomes for 500 stochastic repetitions are shown and compared with the deterministic version. We also provide an indication on calculation time for each model and paradigm. A state machine diagram describes the states and transitions involved in the model. Finally, differences between respective EMULSION files are presented side-by-side and highlighted.

**S2 Text. Changes to introduce age groups and demography in the hybrid SIR model.** Differences between original and modified models (resp. “hybrid_SIR.yaml” and “hybrid_SIR_ages.yaml”) are presented side-by-side and highlighted (in bold; red: parts removed, green: parts added; yellow: parts modified; dashes: similar parts not shown).

**S3 Text. Changes to upscale the previous SIR model with demography to the metapopulation level, with data-driven migrations.** Differences between original and modified models (resp. “hybrid_SIR_ages.yaml” and “hybrid_SIR_ages _metapop.yaml”) are presented side-by-side and highlighted (in bold; red: parts removed, green: parts added; yellow: parts modified; dashes: similar parts not shown).

**S4 Text. Python code for add-ons associated with the metapopulation.** This file (“metapop_movements.py”) is required to connect the model described in the YAML file to data used on trade movements. It is composed of two functions: one for reading the CSV file, the other for moving individuals from one population to another according to data.

**S5 Text. Changes to previous model (metapopulation) to introduce vaccination for a targeted group and a specific period of the year.** Differences between original and modified models (resp. “hybrid_SIR_ages _metapop.yaml” and “hybrid_VSIR_ages_metapop.yaml”) are presented side-by-side and highlighted (in bold; red: parts removed, green: parts added; yellow: parts modified; dashes: similar parts not shown).

**S2 Appendix. Case studies in human, animal and plant epidemiology.** This document demonstrates how to reimplement with EMULSION three published non-trivial models, making their assumptions and structure explicit: 1) a term-timed forced model of measles exhibiting multi-annual outbreaks, and the impact of a vaccination measure; 2) a model of a vector-borne zoonosis, Rift Valley fever; 3) a spatially explicit individual-based model of the spread and control of a plant disease, the Bahia Bark Scaling of Citrus.

**S6 Text. Installation procedure, tests and help page for EMULSION.** Please refer to EMULSION documentation for the latest version: https://sourcesup.renater.fr/www/emulsion-public

**S1 File. Zip file containing all example files mentioned in the software paper.** File “README.md” provides description and usage for each of them.

**S2 File. Zip file containing EMULSION source files.** This zip file is a clone of EMULSION public Git repository. To install from this file rather than from PyPI, go to EMULSION documentation page: https://sourcesup.renater.fr/www/emulsion-public/pages/Install.html and follow Git-based installation instructions.

## Acknowledgments

This work was carried out with the financial support of the French Research Agency (ANR), Program Investments for the Future, project ANR-10-BINF-07 (MIHMES), and the European Union through the European fund for the regional development (FEDER) of Pays-de-la-Loire. It was also funded by the French Research Agency (ANR) through project CADENCE (ANR-16-CE32-0007-01), the Animal Health Division of the French National Institute for Agricultural Research (INRA), and the INRA meta-programme “Sustainable management of animal health” through project FORESEE. We are also very grateful to Thibaut Morel-Journel for his careful reading and relevant comments.

## Notes

#### Summary of Updates

Emphasized the capability of EMULSION to handle models of real diseases in human, animal and plant epidemiology, with new supporting information (S2 Appendix). New subsection "Addressing challenges in disease transmission modelling".

