## Supplementary material for "EMULSION: transparent and flexible multiscale stochastic models in human, animal and plant epidemiology": S1 Text

### S1 Text – EMULSION model file for a SIR model with hybrid modelling paradigm

#### Contents


**A Content of EMULSION model file** **1**

**B Corresponding state machine diagram** **3**


This document provides the full EMULSION file (`hybrid_SIR.yaml`) for describing a SIR model using the hybrid modelling approach (section A) and the corresponding state machine diagram (section B).

Note: EMULSION provides variables which are built automatically from levels and states defined in the model. For instance, defining a level named “population” generates a variable named `total_population` representing the total number of individuals in the population. Similarly, defining a state “S” provides a variable `total_S` containing the number of individuals in S state, and a boolean variable `is_S` per individual, which is 1 if the individual is in state S, and 0 otherwise.

#### A Content of EMULSION model file

```
# general information
model_name: hybrid_SIR

model_info:
  abstract: 'This model is a simple discrete-time, stochastic,
  hybrid SIR model (with individuals grouped automatically).'
  author: 'Sebastien Picault'

time_info:
  # time unit used for all time-related parameters (e.g. rates/durations)
  time_unit: 'days'
  # duration of one time step (in time units)
  delta_t: 1

# definition of modelling paradigm, processes and scales
levels:
  population:
    desc: 'level of the population'
    aggregation_type: 'hybrid'
    contains:
      - individuals
  individuals:
    desc: 'level of the individuals'
```

```yaml
processes:
# only one process, at population level
  population:
    - infection

grouping:
  population:
    # To execute process "infection" in the population, individuals
    # are grouped according to the value of their variable
    # "health_state". Then, for each subgroup, e.g. all "S"
    # (susceptible) individuals, the simulation engine determines
    # which transitions of the "health_state" state machine are
    # available and which individuals will evolve through each transition.
    infection:
      # process "infection" is driven by state machine "health_state"
      machine_name: health_state
      # and individuals in the same "health_state" are grouped together
      key_variables: [health_state]

# description of state machines
state_machines:
  health_state:
    desc: 'The state machine which defines the evolution of health states'
    states:
      # list of states with their properties
      - S:
          name: 'Susceptible'
          desc: 'suceptible of becoming infected'
          fillcolor: 'lightskyblue'
          default: yes
      - I:
          name: 'Infectious'
          desc: 'infected and able to transmit the disease'
          fillcolor: 'red'
      - R:
          name: 'Resistant'
          desc: 'healthy again and resistant to infection'
          fillcolor: 'midnightblue'
    # list of transitions between states
    transitions:
      - {from: S, to: I, rate: 'transmission_I * total_I / total_population'}
      - {from: I, to: R, rate: 'recovery'}

# description and values of parameters/expressions
parameters:
  initial_population_size:
    desc: 'initial number of individuals in the population'
    value: 100
  initial_prevalence:
    desc: 'initial proportion of infectious individuals in the population'
    value: 0.1
  transmission_I:
```

```yaml
    desc: 'transmission rate from infectious individuals (/day)'
    value: 0.5
  recovery:
    desc: 'recovery rate (/day)'
    value: 0.1
  percentage_prevalence:
    desc: 'proportion of infectious individuals (%)'
    value: '100 * total_I / total_population'

# prototypes = examples of typical agents for each level,
# characterized by specific variable values
prototypes:
  individuals:
    - healthy:
        desc: 'healthy individuals'
        health_state: S
    - infected:
        desc: 'infected individuals'
        health_state: I

# initial conditions
initial_conditions:
  population:
    - prototype: healthy
      amount: 'initial_population_size * (1 - initial_prevalence)'
    - prototype: infected
      amount: 'initial_population_size * initial_prevalence'

# outputs frequency and variables to track in addition to amounts of
# individiuals in each state
outputs:
  type: csv
  population:
    period: 1
    extra_vars:
      - percentage_prevalence
...
```

#### B Corresponding state machine diagram

The state machine diagram below can be produced by EMULSION with the following command:

```
emulsion diagrams hybrid_SIR.yaml --format pdf
```

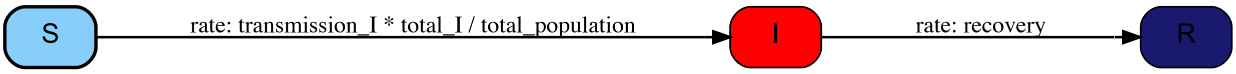
