## Supplementary material for "EMULSION: transparent and flexible multiscale stochastic models in human, animal and plant epidemiology": S1 Appendix

### S1 Appendix – Comparison of modelling paradigms

This document illustrates how two classical models (SIR, section A and SIR with birth and death processes, section B) are implemented using EMULSION using the three modelling paradigms provided by the software: compartment-based, hybrid and invididual-based. For each modelling paradigm and each model, EMULSION outputs are also compared with outputs produced for the same model using the R library SimInf (Widgren *et al.* 2016), which is compartment-based.

A comparison of execution times for each modelling paradigm and each model, based on 500 stochastic repetitions for each EMULSION model, is given in table 1. Simulations were run on a MacBook Pro (3.3 GHz Intel Core i5, 16 Gb RAM, MacOS 10.14). Memory usage was about 140 Mb in each case.

| | | **compartment** | **hybrid** | **IBM** |
|---|---|---|---|---|
| SIR model | duration (s) | 84 | 130 | 167 |
| | *relative* | *1* | *1.5* | *2.0* |
| SIR model + birth/death | duration (s) | 120 | 191 | 387 |
| | *relative* | *1* | *1.6* | *3.2* |
| | *relative to SIR* | *1.4* | *2.3* | *4.6* |

**Table 1:** Execution time (s) for the 500 stochastic repetitions with each modelling paradigm and each model

#### Contents


**A SIR model** **2**

**B SIR model with births and deaths** **6**

### A SIR model

This section compares how a classical SIR model is implemented in EMULSION as compartment-based, hybrid and invididual-based models (respectively, `compartment_SIR.yaml`, `hybrid_SIR.yaml` and `IBM_SIR.yaml`).

All files correspond to the same epidemiological model, represented by the state machine diagram on Figure 1. This diagram can be produced with the following command:

```
emulsion diagrams compartment_SIR.yaml
```

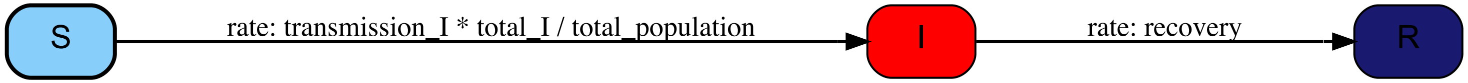


**Figure 1:** State machine diagram associated with a classical SIR model.

The deterministic version of this model can be described through the following Ordinary Differential Equation (ODE) system, where $\beta$ denotes the transmission rate (parameter `transmission_I`), $\gamma$ the recovery rate (`recovery`), and $N = S + I + R$ the total population (`total_population`), and $S, I, R$ the amount of individuals in each health state (resp. susceptible, infectious, resistant).

$$\begin{cases} \frac{\mathrm{d}S}{\mathrm{d}t} &= -S\frac{\beta I}{N} \\ \frac{\mathrm{d}I}{\mathrm{d}t} &= S\frac{\beta I}{N} - \gamma I \\ \frac{\mathrm{d}R}{\mathrm{d}t} &= \gamma I \end{cases}$$

Figure 2 presents simulation outputs produced by EMULSION for each paradigm and by the compartment-based R library SimInf, with 500 stochastic repetitions. The dynamics of the deterministic model was added to each of them.

Figure 3 (resp. 4) shows the differences between the files used to represent the SIR in EMULSION as a compartment-based model and as a hybrid model (resp. as a hybrid model and as an individual-based model).

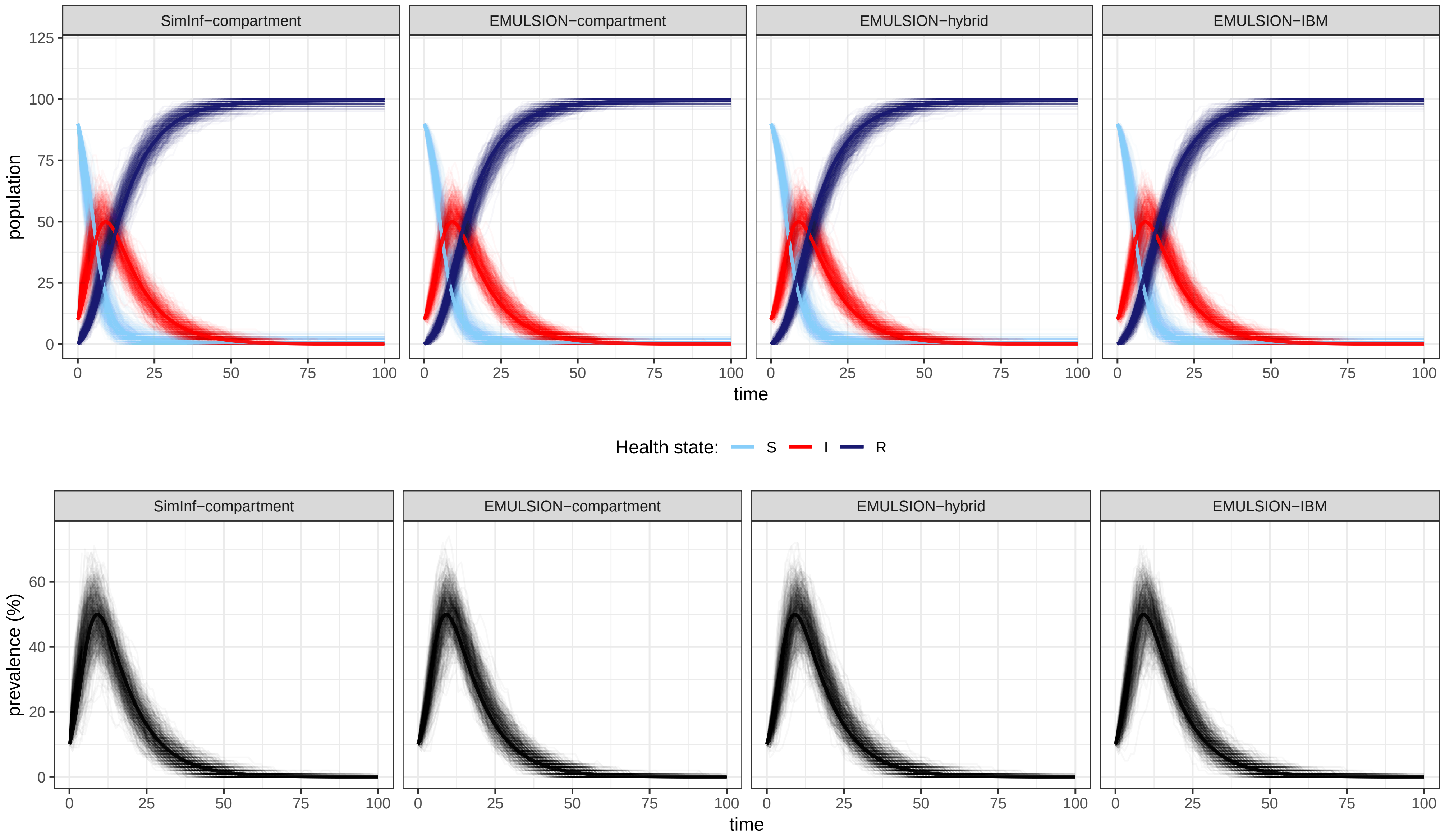


**Figure 2:** Simulation outputs produced for the SIR model (500 stochastic repetitions) with the compartment-based R library SimInf (column 1) and in each modelling paradigm with EMULSION (columns 2-4). Strong lines represent the deterministic version of the model, which corresponds to the average behaviour when infection persists. Parameters: `transmission_I` = 0.5; `recovery` = 0.1; `initial_population_size` = 100 ; `initial_prevalence` = 0.1

compartment_SIR.yaml

```
# general information
model_name: compart_SIR

model_info:
  abstract: 'This model is a simple discrete-time, stochastic,
  compartment-based SIR model.'
  author: 'Sebastien Picault'

---
# definition of modelling paradigm, processes and scales
levels:
  population:
    desc: 'level of the population'
    aggregation_type: 'compartment'

---
  percentage_prevalence:
    desc: 'proportion of infectious individuals (%)'
    value: '100 * total_I / total_population'

# initial conditions
initial_conditions:
  population:
    - population:
        - total: 'initial_population_size'
        - vars: [I]
          amount: 'initial_population_size * initial_prevalence'

```

hybrid_SIR.yaml

```
# general information
model_name: hybrid_SIR

model_info:
  abstract: 'This model is a simple discrete-time, stochastic,
  hybrid SIR model (with individuals grouped automatically).'
  author: 'Sebastien Picault'

---
# definition of modelling paradigm, processes and scales
levels:
  population:
    desc: 'level of the population'
    aggregation_type: 'hybrid'
    contains:
      - individuals
  individuals:
    desc: 'level of the individuals'

---
  percentage_prevalence:
    desc: 'proportion of infectious individuals (%)'
    value: '100 * total_I / total_population'

# prototypes = examples of typical agents for each level,
# characterized by specific variable values
prototypes:
  individuals:
    - healthy:
        desc: 'healthy individuals'
        health_state: S
    - infected:
        desc: 'infected individuals'
        health_state: I

# initial conditions
initial_conditions:
  population:
    - prototype: healthy
      amount: 'initial_population_size * (1 - initial_prevalence)'
    - prototype: infected
      amount: 'initial_population_size * initial_prevalence'

```

**Figure 3:** Differences between the compartment-based (`compartment_SIR.yaml`) and the hybrid (`hybrid_SIR.yaml`) versions of the SIR model in EMULSION, presented side-by-side and highlighted (in bold; red: parts removed, green: parts added; yellow: parts modified; dashes: similar parts not shown).

hybrid_SIR.yaml

```
# general information
model_name: hybrid_SIR
model_info:
  abstract: 'This model is a simple discrete-time, stochastic,
  hybrid SIR model (with individuals grouped automatically).'
  author: 'Sebastien Picault'
---
# definition of modelling paradigm, processes and scales
levels:
  population:
    desc: 'level of the population'
    aggregation_type: 'hybrid'
    contains:
      - individuals
  individuals:
    desc: 'level of the individuals'
processes:
# only one process, at population level
  population:
    - infection
grouping:
  population:
    infection:
      # process "infection" is driven by state machine "health_state"
      machine_name: health_state
      # and individuals in the same "health_state" are grouped together
      key_variables: [health_state]
```

IBM_SIR.yaml

```
# general information
model_name: IBM_SIR
model_info:
  abstract: 'This model is a simple discrete-time, stochastic,
  individual-based SIR model (with fully autonomous individuals).'
  author: 'Sebastien Picault'
---
# definition of modelling paradigm, processes and scales
levels:
  population:
    desc: 'level of the population'
    aggregation_type: 'IBM'
    contains:
      - individuals
  individuals:
    desc: 'level of the individuals'
processes:
# only one process, at individuals level, driven directly by state machines
  individuals:
    - health_state
```

**Figure 4:** Differences between the hybrid (`hybrid_SIR.yaml`) and the individual-based (`IBM_SIR.yaml`) versions of the SIR model in EMULSION, presented side-by-side and highlighted (in bold; red: parts removed, green: parts added; yellow: parts modified; dashes: similar parts not shown).

#### B SIR model with births and deaths

This section compares how a classical SIR model with birth and death processes is implemented in EMULSION as compartment-based, hybrid and invididual-based models (respectively, `compartment_SIR_demo.yaml`, `hybrid_SIR_demo.yaml` and `IBM_SIR_demo.yaml`).

All files correspond to the same epidemiological model, represented by the state machine diagram on Figure 5. This diagram can be produced with the following command:

```
emulsion diagrams compartment_SIR_demo.yaml
```

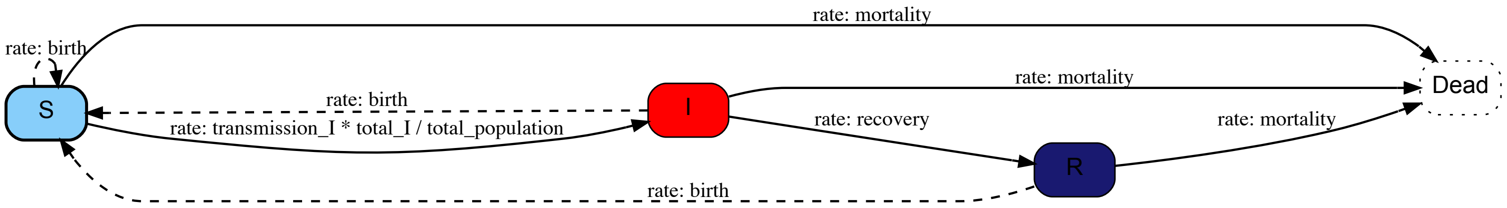


**Figure 5:** State machine diagram associated with a SIR model with birth and death processes. The "Dead" state surrounded by dots is intended to represent individuals removed from the system; dashed arrows represent production links, i.e. how individuals produce new individuals.

The deterministic version of this model can be described through the following Ordinary Differential Equation (ODE) system, where $\beta$ denotes the transmission rate (parameter `transmission_I`), $\gamma$ the recovery rate (`recovery`), $b$ the birth rate (`birth`), $\mu$ the mortality rate (`mortality`), and $N = S+I+R$ the total population (`total_population`), and $S, I, R$ the amount of individuals in each health state (resp. susceptible, infectious, resistant).

$$\begin{cases} \frac{\mathrm{d}S}{\mathrm{dt}} &= -S\frac{\beta I}{N} + bN - \mu S \\ \frac{\mathrm{d}I}{\mathrm{dt}} &= S\frac{\beta I}{N} - \gamma I - \mu I \\ \frac{\mathrm{d}R}{\mathrm{dt}} &= \gamma I - \mu R \end{cases}$$

Figure 6 presents simulation outputs produced by EMULSION for each paradigm and by the compartment-based R library SimInf, with 500 stochastic repetitions. The dynamics of the deterministic model was added to each of them.

Figure 7 (resp. 8) shows the differences between the files used to represent the SIR in EMULSION as a compartment-based model and as a hybrid model (resp. as a hybrid model and as an individual-based model).

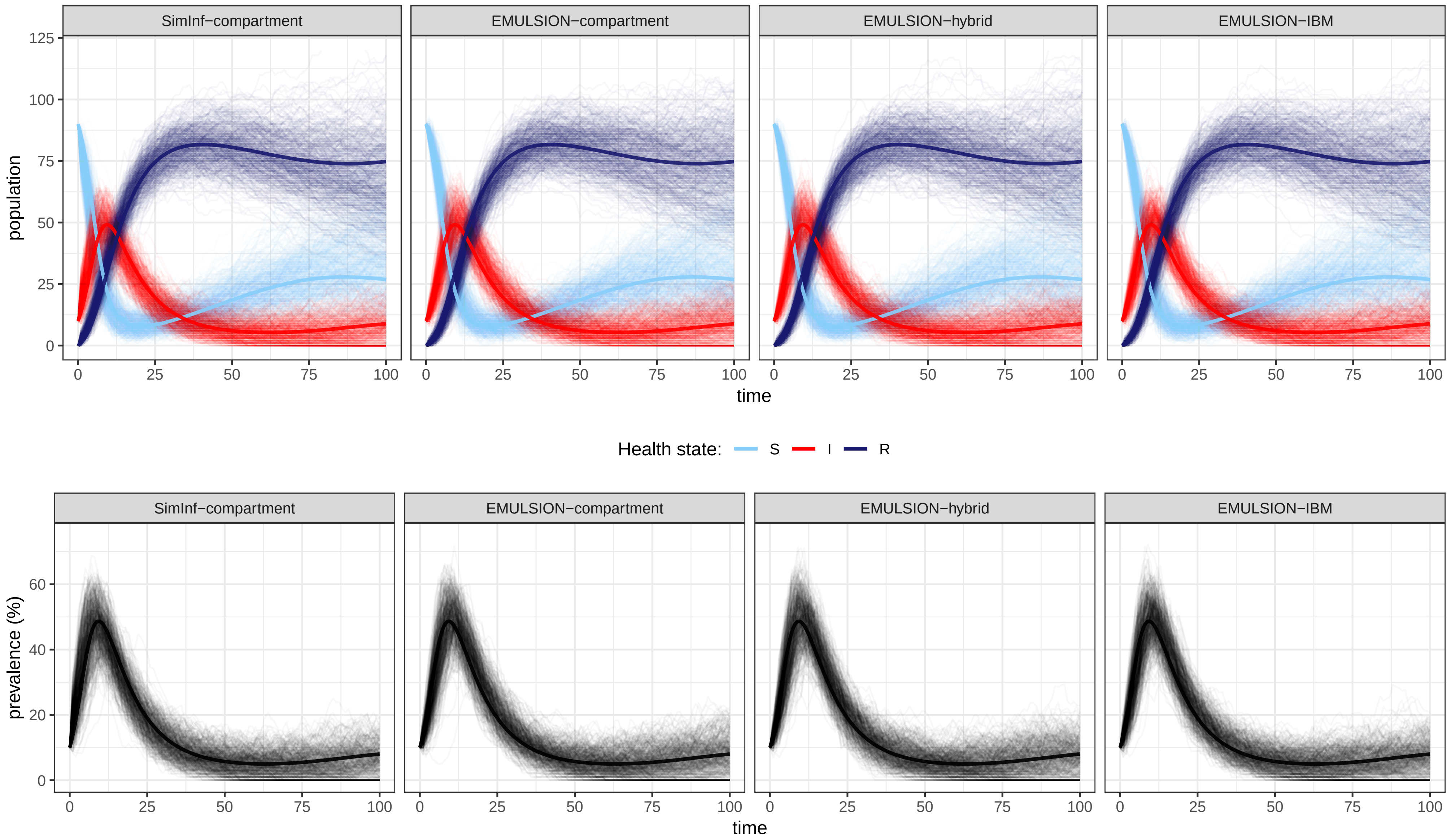


**Figure 6:** Simulation outputs produced for the SIR model with birth/death (500 stochastic repetitions) with the compartment-based R library SimInf (column 1) and in each modelling paradigm with EMULSION (columns 2–4). Strong lines represent the deterministic version of the model, which corresponds to the average behaviour when infection persists. Parameters: `transmission_I` = 0.5; `recovery` = 0.1; `birth` = 0.011; `mortality` = 0.01; `initial_population_size` = 100 ; `initial_prevalence` = 0.1

compartment_SIR_demo.yaml

```
# general information
model_name: compart_SIR_demo

model_info:
  abstract: 'This model is a simple discrete-time, stochastic,
  compartment-based SIR model with births and deaths.'

  author: 'Sebastien Picault'

---
# definition of modelling paradigm, processes and scales
levels:
  population:
    desc: 'level of the population'
    aggregation_type: 'compartment'

---
    productions:
      - {from: S, to: S, rate: 'birth'}
      - {from: I, to: S, rate: 'birth'}
      - {from: R, to: S, rate: 'birth'}

---
  mortality:
    desc: 'mortality rate (/day)'
    value: 0.01

# initial conditions
initial_conditions:
  population:
    - population:
      - total: 'initial_population_size'
      - vars: [I]
        amount: 'initial_population_size * initial_prevalence'

```

hybrid_SIR_demo.yaml

```
# general information
model_name: hybrid_SIR_demo

model_info:
  abstract: 'This model is a simple discrete-time, stochastic,
  hybrid SIR model (with individuals grouped automatically),
  with births and deaths.'
  author: 'Sebastien Picault'

---
# definition of modelling paradigm, processes and scales
levels:
  population:
    desc: 'level of the population'
    aggregation_type: 'hybrid'
    contains:
      - individuals
  individuals:
    desc: 'level of the individuals'

---
    productions:
      - {from: S, to: S, rate: 'birth', prototype: 'healthy'}
      - {from: I, to: S, rate: 'birth', prototype: 'healthy'}
      - {from: R, to: S, rate: 'birth', prototype: 'healthy'}

---
  mortality:
    desc: 'mortality rate (/day)'
    value: 0.01

# prototypes = examples of typical agents for each level,
# characterized by specific variable values
prototypes:
  individuals:
    - healthy:
        desc: 'healthy individuals'
        health_state: S
    - infected:
        desc: 'infected individuals'
        health_state: I

# initial conditions
initial_conditions:
  population:
    - prototype: healthy
      amount: 'initial_population_size * (1 - initial_prevalence)'
    - prototype: infected
      amount: 'initial_population_size * initial_prevalence'

```

**Figure 7:** Differences between the compartment-based (`compartment_SIR_demo.yaml`) and the hybrid (`hybrid_SIR_demo.yaml`) versions of the SIR model with birth/death in EMULSION, presented side-by-side and highlighted (in bold; red: parts removed, green: parts added; yellow: parts modified; dashes: similar parts not shown).

hybrid_SIR_demo.yaml

```
# general information
model_name: hybrid_SIR_demo

model_info:
  abstract: 'This model is a simple discrete-time, stochastic,
  hybrid SIR model (with individuals grouped automatically),
  with births and deaths.'
  author: 'Sebastien Picault'

---
# definition of modelling paradigm, processes and scales
levels:
  population:
    desc: 'level of the population'
    aggregation_type: 'hybrid'
    contains:
      - individuals
  individuals:
    desc: 'level of the individuals'

processes:
# only one process, at population level
  population:
    - infection

grouping:
  population:
    infection:
      # process "infection" relies on the health_state state machine
      machine_name: health_state
      # and on the grouping of all "individuals" in the same health state
      key_variables: [health_state]

```

IBM_SIR_demo.yaml

```
# general information
model_name: IBM_SIR_demo

model_info:
  abstract: 'This model is a simple discrete-time, stochastic,
  individual-based SIR model (with fully autonomous individuals),
  with births and deaths.'
  author: 'Sebastien Picault'

---
# definition of modelling paradigm, processes and scales
levels:
  population:
    desc: 'level of the population'
    aggregation_type: 'IBM'
    contains:
      - individuals
  individuals:
    desc: 'level of the individuals'

processes:
# only one process, at individuals level, driven directly by state machines
  individuals:
    - health_state

```

**Figure 8:** Differences between the hybrid (`hybrid_SIR_demo.yaml`) and the individual-based (`IBM_SIR_demo.yaml`) versions of the SIR model with birth/death in EMULSION, presented side-by-side and highlighted (in bold; red: parts removed, green: parts added; yellow: parts modified; dashes: similar parts not shown).
