## Supplementary material for "EMULSION: transparent and flexible multiscale stochastic models in human, animal and plant epidemiology": S2 Text

hybrid_SIR.yaml

```yaml
# general information
model_name: hybrid_SIR

model_info:
  abstract: 'This model is a simple discrete-time, stochastic,
  hybrid SIR model (with individuals grouped automatically).'
  author: 'Sebastien Picault'
---

processes:
# only one process, at population level
  population:
    - infection

grouping:
  population:
    infection:
      # process "infection" is driven by state machine "health_state"
      machine_name: health_state
      # and individuals in the same "health_state" are grouped together
      key_variables: [health_state]

---
    # list of transitions between states
    transitions:
      - {from: S, to: I, rate: 'transmission_I * total_I / total_population'}
      - {from: I, to: R, rate: 'recovery'}

```

hybrid_SIR_ages.yaml

```yaml
# general information
model_name: hybrid_SIR_ages

model_info:
  abstract: 'This model is a simple discrete-time, stochastic,
  hybrid SIR model (with individuals grouped automatically), with
  demography based on age groups.'
  author: 'Sebastien Picault'
---

processes:
# now two processes at population level
  population:
    - infection
    - aging

grouping:
  population:
    infection:
      # process "infection" is driven by state machine "health_state"
      machine_name: health_state
      # and individuals in the same "health_state" are grouped together
      key_variables: [health_state]
    aging:
      # process "aging" is driven by state machine "age_group"
      machine_name: age_group
      # and individuals in the same "age_group" are grouped together
      key_variables: [age_group]

---
    # list of transitions between states
    transitions:
      - {from: S, to: I, rate: 'transmission_I * total_I / total_population'}
      - {from: I, to: R, rate: 'recovery'}
  age_group:
    desc: 'The state machine which defines the evolution of age groups.'
    states:
      - J:
          name: 'Juvenile'
          desc: 'juvenile individuals'
          fillcolor: 'pink'
      - A:
          name: 'Adult'
          desc: 'adult individuals (producing new juveniles)'
          fillcolor: 'brown'
      - Dead:
          name: 'Dead'
          desc: 'compartment to put dead animals'
          fillcolor: white
          autoremove: yes
    transitions:
      - {from: J, to: A, rate: 'maturation'}
      # death (assuming no disease-induced mortality)
      - {from: J, to: Dead, rate: 'mortality'}
      - {from: A, to: Dead, rate: 'mortality'}
    productions:
      # births (assuming no vertical transmission)
      - {from: A, to: J, rate: 'birth', prototype: 'newborn'}

```

```
# description and values of parameters/expressions
parameters:
  initial_population_size:
    desc: 'initial number of individuals in the population'
---
  percentage_prevalence:
    desc: 'proportion of infectious individuals (%)'
    value: '100 * total_I / total_population'

# prototypes = examples of typical agents for each level,
# characterized by specific variable values
prototypes:
  individuals:
    - healthy:
        desc: 'healthy individuals'
        health_state: S

    - infected:
        desc: 'infected individuals'
        health_state: I

---
  type: csv
  population:
    period: 1
    extra_vars:
      - percentage_prevalence

...
```

```
# description and values of parameters/expressions
parameters:
  initial_population_size:
    desc: 'initial number of individuals in the population'
---
  percentage_prevalence:
    desc: 'proportion of infectious individuals (%)'
    value: '100 * total_I / total_population'
  maturation:
    desc: 'rate at which juveniles become adults (/day)'
    value: '1/20'
  mortality:
    desc: 'the mortality rate (/day)'
    value: 0.01
  birth:
    desc: 'the birth rate (/day), here calculated to approximatively
    balance mortality and maturation in this model'
    value: 0.012

# prototypes = examples of typical agents for each level,
# characterized by specific variable values
prototypes:
  individuals:
    - healthy:
        desc: 'healthy individuals'
        health_state: S
        age_group: random
    - infected:
        desc: 'infected individuals'
        health_state: I
        # here we assume that initially infected individuals are adults
        age_group: A
    - newborn:
        desc: 'newly created individuals'
        health_state: default
        age_group: J

---
  type: csv
  population:
    period: 1
    extra_vars:
      - percentage_prevalence
      - total_population
...
```
