## Supplementary material for "EMULSION: transparent and flexible multiscale stochastic models in human, animal and plant epidemiology": S3 Text

hybrid_SIR_ages.yaml

```yaml
# general information
model_name: hybrid_SIR_ages

model_info:
  abstract: 'This model is a simple discrete-time, stochastic,
  hybrid SIR model (with individuals grouped automatically), with
  demography based on age groups.'

  author: 'Sebastien Picault'

time_info:
  # time unit used for all time-related parameters (e.g. rates/durations)
  time_unit: 'days'
  # duration of one time step (in time units)
  delta_t: 1

# definition of modelling paradigm, processes and scales
levels:
  population:
    desc: 'level of the population'
    aggregation_type: 'hybrid'
    contains:
      - individuals
  individuals:
    desc: 'level of the individuals'

processes:
# now two processes at population level
  population:
    - infection
    - aging

---
parameters:
  initial_population_size:
    desc: 'initial number of individuals in the population'
    value: 100
  initial_prevalence:
    desc: 'initial proportion of infectious individuals in the population'
    value: 0.1
  transmission_I:
    desc: 'transmission rate from infectious individuals (/day)'
```

hybrid_SIR_ages_metapop.yaml

```yaml
# general information
model_name: hybrid_SIR_ages_metapop

model_info:
  abstract: 'This model is a simple discrete-time, stochastic,
  hybrid SIR model (with individuals grouped automatically), with
  demography based on age groups, at the metapopulation scale, with
  data-based trade movements.'
  author: 'Sebastien Picault'

time_info:
  # time unit used for all time-related parameters (e.g. rates/durations)
  time_unit: 'days'
  # duration of one time step (in time units)
  delta_t: 1
  # specify a longer simulation duration than default
  total_duration: 200
  # specify the date where simulation starts (to connect with recorded data)
  origin: 'January 1, 2018'

# definition of modelling paradigm, processes and scales
levels:
  population:
    desc: 'level of the population'
    aggregation_type: 'hybrid'
    contains:
      - individuals
  individuals:
    desc: 'level of the individuals'
  metapop:
    desc: 'level of the metapopulation'
    contains:
      - population
    aggregation_type: 'metapopulation'
    # The metapopulation is explicitly linked to a specific class in a
    # Python code add-on
    file: metapop_movements.py
    class_name: Metapopulation

processes:

  population:
    - infection
    - aging
  metapop:
    # here the process is the name of a procedure defined in the
    # Python code
    - exchange_individuals

---
parameters:
  initial_population_size:
    desc: 'initial number of individuals in the population'
    value: 100

  transmission_I:
    desc: 'transmission rate from infectious individuals (/day)'
```

```
    value: 0.5
---
  birth:
    desc: 'the birth rate (/day), here calculated to approximatively
    balance mortality and maturation in this model'
    value: 0.012

# prototypes = examples of typical agents for each level,
# characterized by specific variable values
---
    - newborn:
        desc: 'newly created individuals'
        health_state: default
        age_group: J

# initial conditions
initial_conditions:
  population:
    - prototype: healthy
      amount: 'initial_population_size * (1 - initial_prevalence)'
    - prototype: infected
      amount: 'initial_population_size * initial_prevalence'

```

```
    value: 0.5
---
  birth:
    desc: 'the birth rate (/day), here calculated to approximatively
    balance mortality and maturation in this model'
    value: 0.012
  nb_pops:
    desc: 'number of populations in metapopulation'
    value: 10
  init_prevalent_pop:
    desc: 'initial proportion of infected populations'
    value: 0.1

# variables
statevars:
  initial_prevalence:
    desc: 'initial proportion of infectious individuals in the population
    (now a variable of each population instead of a simulation parameter)'

# prototypes = examples of typical agents for each level,
# characterized by specific variable values
---
    - newborn:
        desc: 'newly created individuals'
        health_state: default
        age_group: J
    - imported_movement:
        desc: 'profile of individuals from outside the metapopulation,
        assuming no external risk'
        health_state: S
        age_group: default
  population:
    - healthy_pop:
        desc: 'populaiton initially infection-free'
        initial_prevalence: 0
    - infected_pop:
        desc: 'population initially infected with prevalence 0.1'
        initial_prevalence: 0.1

# initial conditions
initial_conditions:
  population:
    - prototype: healthy
      amount: 'initial_population_size * (1 - initial_prevalence)'
    - prototype: infected
      amount: 'initial_population_size * initial_prevalence'
  metapop:
    - prototype: infected_pop
      amount: 'nb_pops * init_prevalent_pop'
    - prototype: healthy_pop
      amount: 'nb_pops * (1 - init_prevalent_pop)'

```
