## Supplementary material for "EMULSION: transparent and flexible multiscale stochastic models in human, animal and plant epidemiology": S4 Text

```
"""Definition of specific code add-on for the EMULSION model to handle
commercial movements in the metapopulation.

"""

from   pathlib                import Path
import csv
import dateutil.parser       as     dup
import numpy                 as     np
from   emulsion.agent.managers import MetapopProcessManager

DATA_FILE = 'moves.csv'

#===============================================================
# CLASS Metapopulation (LEVEL 'metapop')
#===============================================================
class Metapopulation(MetapopProcessManager):
    """
    level of the metapop.

    """

    #----------------------------------------------------------------
    # Level initialization. This method is called automatically by
    # EMULSION simulation engine after the application of initial
    # conditions.
    # ----------------------------------------------------------------
    def initialize_level(self, **others):
        """Initialize an instance of Metapopulation.
        Additional initialization parameters can be introduced here if needed.
        """
        # read a CSV data file for moves:
        # date of movement, source pop, destination pop, age group, quantity

        # and restructure it according to origin_date and delta_t:
        # {step: {source_id: [(dest_id, age_group, qty), ...],
        #         ...},
        #  ...}
        origin = self.model.origin_date
        step_duration = self.model.step_duration
        moves = {}
        with open(Path(self.model.input_dir, DATA_FILE)) as csvfile:
            # read the CSV file
            csvreader = csv.DictReader(csvfile, delimiter=',')
            for row in csvreader:
                day = dup.parse(row['date'])
                if day < origin:
                    # ignore dates before origin_date
                    continue
                # convert dates into simulation steps
                step = (day - origin) // step_duration
                # group information by step and source pop
                if step not in moves:
```

```
                moves[step] = {}
            src, dest, qty = int(row['source']), int(row['dest']), int(row['qty'])
            if src not in moves[step]:
                moves[step][src] = []
            moves[step][src].append([dest, row['age'], qty])
    self.moves = moves

#-----------------------------------------------------------------
# Processes
#-----------------------------------------------------------------
def exchange_individuals(self):
    """Check if movements have to be performed at current time step. If
    so, identify source and target populations, select individuals
    corresponding to the data, and proceed to the movement.

    """
    if self.statevars.step in self.moves:
        pops = self.get_populations()
        for source in self.moves[self.statevars.step]:
            for dest, age, qty in self.moves[self.statevars.step][source]:
                # neither source/dest in simulated pops
                if (source not in pops) and (dest not in pops):
                    # ignoring movement from source to dest (outside the metapopulation)
                    continue
                # source not in simulated pops: create individual from prototype
                if source not in pops:
                    # movement to dest coming from outside the metapopulation
                    # retrieve prototype definition from the model
                    prototype = self.model.get_prototype(name='imported_movement',
                                                         level='individuals')
                    # change age group to comply with movement
                    prototype['age_group'] = self.get_model_value(age)
                    individuals = [pops[dest].new_atom(custom_prototype=prototype)
                                   for _ in range(qty)]
                else:
                    # find convenient individuals
                    candidates = pops[source].select_atoms('age_group', age,
                                                           process='aging')
                    # try to move the appropriate quantity
                    nb = min(len(candidates), qty)
                    if nb > 0:
                        individuals = np.random.choice(candidates, nb, replace=False)
                        pops[source].remove_atoms(individuals)
                    else:
                        individuals = []
                if dest not in pops:
                    pass
                    # movement from dest going outside the metapopulation
                else:
                    pops[dest].add_atoms(individuals)
```
