## Supplementary material for "EMULSION: transparent and flexible multiscale stochastic models in human, animal and plant epidemiology": S5 Text

hybrid_SIR_ages_metapop.yaml

```yaml
# general information
model_name: hybrid_SIR_ages_metapop

model_info:
  abstract: 'This model is a simple discrete-time, stochastic,
  hybrid SIR model (with individuals grouped automatically), with
  demography based on age groups, at the metapopulation scale, with
  data-based trade movements.'
  author: 'Sebastien Picault'
---
  # specify the date where simulation starts (to connect with recorded data)
  origin: 'January 1, 2018'

---
      - R:
          name: 'Resistant'
          desc: 'healthy again and resistant to infection'
          fillcolor: 'midnightblue'
    # list of transitions between states
    transitions:
      - {from: S, to: I, rate: 'transmission_I * total_I / total_population'}
      - {from: I, to: R, rate: 'recovery'}
  age_group:
    desc: 'The state machine which defines the evolution of age groups.'
---
  init_prevalent_pop:
    desc: 'initial proportion of infected populations'
    value: 0.1

```

hybrid_VSIR_ages_metapop.yaml

```yaml
# general information
model_name: hybrid_VSIR_ages_metapop

model_info:
  abstract: 'This model is a simple discrete-time, stochastic,
  hybrid SIR model (with individuals grouped automatically), with
  demography based on age groups, at the metapopulation scale, with
  data-based trade movements. Vaccination is introduced for a targeted
  group and a specific period of the year'
  author: 'Sebastien Picault'
---
  # specify the date where simulation starts (to connect with recorded data)
  origin: 'January 1, 2018'
  calendars:
    # this calendar defines the vaccination period
    vaccinal:
      period: {days: 365}
      events:
        spring: {begin: 'March 21', end: 'June, 21'}

---
      - R:
          name: 'Resistant'
          desc: 'healthy again and resistant to infection'
          fillcolor: 'midnightblue'
      - V:
          name: 'Vaccinated'
          desc: 'temporarily immune to infection'
          fillcolor: 'orange'
    # list of transitions between states
    transitions:
      - {from: S, to: I, rate: 'transmission_I * total_I / total_population'}
      - {from: I, to: R, rate: 'recovery'}
      # vaccinate Juveniles in spring
      - {from: S, to: V, when: 'spring', cond: is_J, rate:
        'vaccination'}
      # vaccinal protection gradually fades out
      - {from: V, to: S, rate: 'waning'}
  age_group:
    desc: 'The state machine which defines the evolution of age groups.'
---
  init_prevalent_pop:
    desc: 'initial proportion of infected populations'
    value: 0.1
  vaccination:
    desc: 'rate at which individuals are vaccintated (/day)'
    value: 0.1
  waning:
    desc: 'rate at which vaccinal protection is lost (/day)'
    value: '1/100'

```
