## Supplementary material for "EMULSION: transparent and flexible multiscale stochastic models in human, animal and plant epidemiology": S2 Appendix

### S2 Appendix – Case studies in human, animal or plant epidemiology

This document illustrates how EMULSION features can help model real diseases in human, animal and plant epidemiology. Three non-trivial models used in real-disease studies from the literature are re-implemented here using EMULSION, demonstrating how the DSL can cover a broad diversity of model structures and applied issues.

All files required to reproduce theses figures (EMULSION models, Python code add-on, R and bash scripts) are provided in `S1_file.zip`.

#### Contents

**A Human health: term-time forced model for multi-annual measles seasonality and introduction of vaccination**
**(Keeling & Rohani 2008, ch. 5)** **2**

**B Animal health: a model of a vector-borne disease, Rift Valley fever**
**(Cavalerie *et al.* 2015)** **8**

**C Plant health: a spatially explicit individual-based model of Bahia Bark Scaling of Citrus spread and control**
**(Cunniffe *et al.* 2014)** **15**

### A Human health: term-time forced model for multi-annual measles seasonality and introduction of vaccination

#### Original model

This section reproduces a temporally forced model for measles, described in Keeling and Rohani, *Modeling Infectious Diseases in Humans and Animals* (2008, Princeton University Press), § 5.2.5. Measles dynamics is represented by a SEIR model with births and deaths (fig. 1), accounting for possible vaccination of newborns. Besides, the authors assume a temporal forcing of the transmission rate:

$$\beta(t) = \frac{\beta_0}{\frac{1}{365}((1+b_1)D_+ + (1-b_1)D_-)}(1 + b_1 Term(t))$$

where $D_+$ and $D_-$ denote the numbers of school and holiday days respectively, $b_1$ the amplitude of seasonality, and $\beta_0$ the average transmission rate (so that $\overline{\beta(t)} = \beta_0$). The temporal forcing is based on real holiday periods (Christmas, Dec. 21–Jan. 6; Easter, Apr. 10–25; Summer, Jul. 19–Sep. 9; Autumn, Oct. 27–Nov. 3). This forcing, combined to the amplitude of seasonality, is responsible for multi-annual measles outbreaks (for instance, a 3-year cycle with low values of $b_1$).

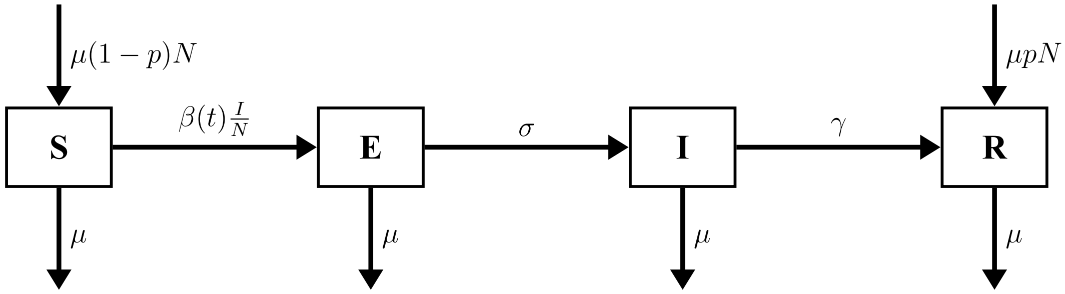


**Figure 1:** Flow diagram corresponding to the temporally forced measles model (after Keeling and Rohani 2008). Parameters: $\mu$: birth/mortality rate; $N$: total population; $p$: proportion of vaccinated newborns.

#### Implementation in EMULSION

This model was re-implemented with EMULSION as a stochatic compartment-based model (`measles.yaml`, p. 5). The corresponding state machine diagram is provided on fig. 2.

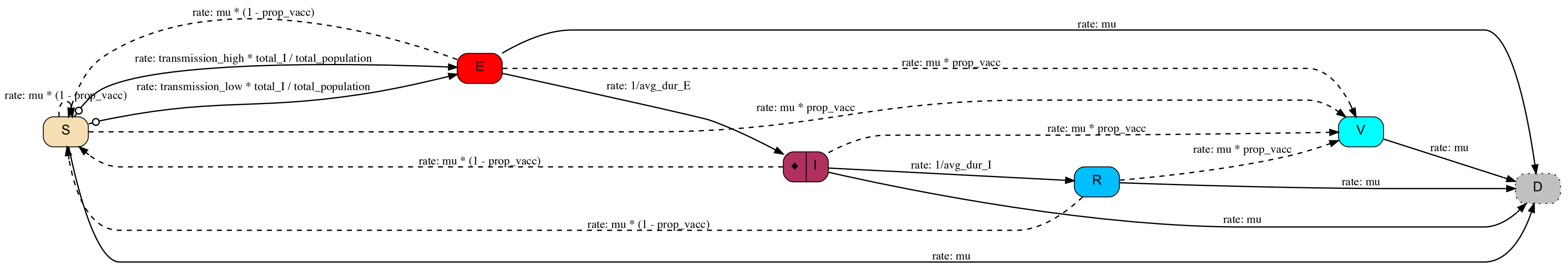


**Figure 2:** State machine diagram for the measles model in EMULSION. Plain arrows denote transitions between states, while dashed arrows denote production links (creation of new individuals). Removal of individuals is made explicit through the "D" (deceased) state, which can be recognized as a sink due to the dotted box. The circle on the transitions from S to E indicates a calendar condition (theses transitions are available on given periods only). The lozenge in I state signals an action when entering state I (actually, recording incidence). This figure was produced as follows: `emulsion diagrams measles.yaml --format pdf`

##### Reproduction of published results

We used this model to exhibit the multi-annual dynamics of measles outbreaks, expected as explained in Keeling and Rohani 2008, and show the impact of vaccination (fig. 3).

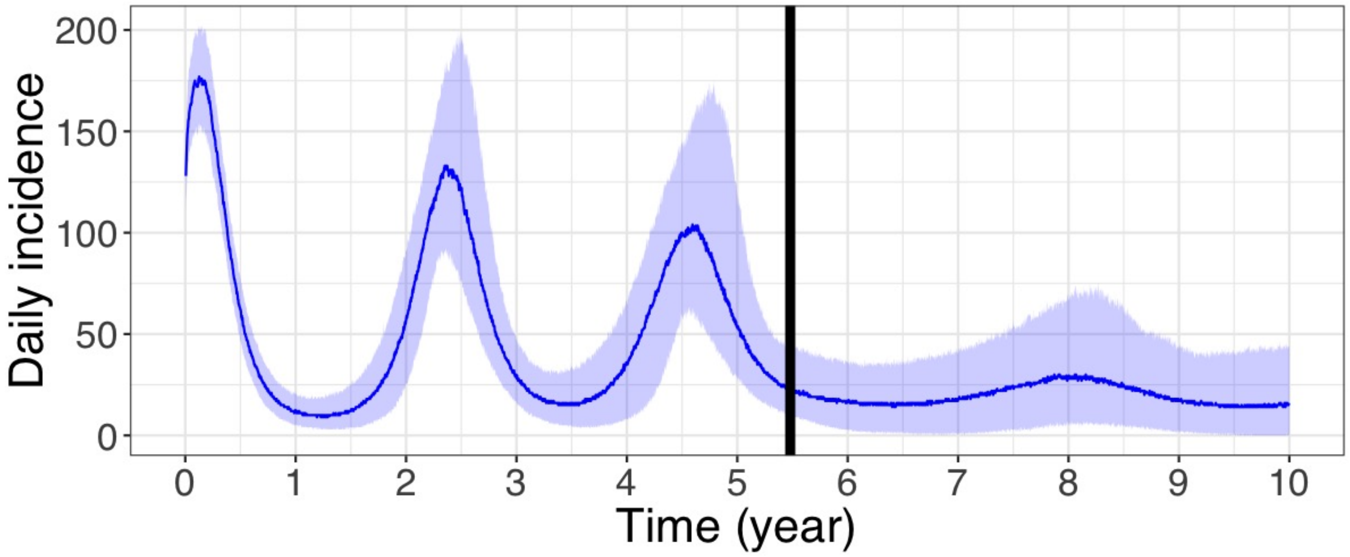


**Figure 3:** Daily incidence in the measles model (500 stochastic repetitions: median and 90% credibility interval), clearly exhibiting multi-annual outbreaks. After t=2000 days (black vertical line), vaccination is applied to 50% of newborns, leading to either to eradication or a reduced and delayed incidence peak. Parameters: see `measles.yaml`.

##### Notes on EMULSION features used for reimplementing the model

- Term-time forcing is done using an explicit **calendar** of holiday periods, defined as events. EMULSION automatically associates boolean tests with each event, so that they can be used to specify alternative transitions from S to E with the `when` keyword (lines 71–78), depending on the period of the year (school vs. holiday)
- The incidence is calculated as a cumulate variable, using an action when entering state I (line 56)
- The proportion of vaccinated newborns is determined dynamically according to when the vaccination campaign starts (lines 109–114)

##### Model in EMULSION DSL (`measles.yaml`)

```yaml
---
model_name: compart_seasonal_measles

model_info:
  abstract: 'This model is a simple discrete-time, stochastic
  implementation of the measles model described in Keeling & Rohani
  2008, Modelling Infectious Diseases in Humans and Animals.'
  author: 'Sebastien Picault'

time_info:
  time_unit: 'weeks'
  delta_t: 1
  origin: 'January 1, 2000'
  total_duration: '365*10'
  calendars:
    holidays:
      period: {days: 365}
      events:
        summer: {begin: 'July 19', end: 'September 8'}
        autumn: {begin: 'October 27', end: 'November 3'}
        christmas: {begin: 'December 21', end: 'January 6'}
        easter: {begin: 'April 10', end: 'April 25'}

levels:
  population:
    desc: 'level of the population'
    aggregation_type: 'compartment'

processes:
  population:
    - infection

grouping:
  population:
    infection:
      machine_name: health_state
      key_variables: [health_state]

state_machines:
  health_state:
    desc: 'The state machine which defines the evolution of health states in the disease'
    states:
      - S:
          name: 'Susceptible'
          desc: 'suceptible of becoming infected'
          fillcolor: 'wheat'
      - E:
          name: 'Exposed'
          desc: 'infected but not yet able to transmit the disease'
          fillcolor: 'red'
      - I:
          name: 'Infectious'
          desc: 'infected and able to transmit the disease'
          fillcolor: 'maroon'
          on_enter:
            - record_change: 'cum_incidence'
      - R:
          name: 'Resistant'
          desc: 'healthy again and resistant to infection'
          fillcolor: 'deepskyblue'
      - V:
          name: 'Vaccinated'
          desc: 'resistant to infection due to vaccination'
          fillcolor: 'cyan'
```

```
      - D:
          name: 'Dead'
          desc: 'compartment to represent deceased individuals'
          fillcolor: 'gray'
          autoremove: yes
    transitions:
      - from: S
        to: E
        when: 'Not(OR(summer, autumn, christmas, easter))'
        rate: 'transmission_high * total_I / total_population'
      - from: S
        to: E
        when: 'OR(summer, autumn, christmas, easter)'
        rate: 'transmission_low * total_I / total_population'
      - {from: E, to: I, rate: '1/avg_dur_E'}
      - {from: I, to: R, rate: '1/avg_dur_I'}
      - {from: S, to: D, rate: 'mu'}
      - {from: E, to: D, rate: 'mu'}
      - {from: I, to: D, rate: 'mu'}
      - {from: R, to: D, rate: 'mu'}
      - {from: V, to: D, rate: 'mu'}
    productions:
      - {from: S, to: S, rate: 'mu * (1 - prop_vacc)'}
      - {from: E, to: S, rate: 'mu * (1 - prop_vacc)'}
      - {from: I, to: S, rate: 'mu * (1 - prop_vacc)'}
      - {from: R, to: S, rate: 'mu * (1 - prop_vacc)'}
      - {from: S, to: V, rate: 'mu * prop_vacc'}
      - {from: E, to: V, rate: 'mu * prop_vacc'}
      - {from: I, to: V, rate: 'mu * prop_vacc'}
      - {from: R, to: V, rate: 'mu * prop_vacc'}

parameters:
  initial_pop_size:
    desc: 'initial size of the population'
    value: 1000000
  prop_S:
    desc: 'initial proportion of susceptible individuals in the population'
    value: 0.06
  prop_EI:
    desc: 'initial proportion of exposed and of infectious individuals in the population'
    value: 0.001
  mu:
    desc: 'mortality/birth rate (/day)'
    value: '0.02 / 365'
  prop_vacc:
    desc: 'proportion of vaccinated newborn (depends on when vaccination campaign starts)'
    value: 'IfThenElse(time < vaccination_start, 0, 0.5)'
  vaccination_start:
    desc: 'Delay between the beginning of the simulation and the beginning of the vaccination campaign (days)'
    value: 2000
  transmission_high:
    desc: 'transmission rate during school terms'
    value: 'corrected_beta0 * (1 + b1)'
  transmission_low:
    desc: 'transmission rate during holidays'
    value: 'corrected_beta0 * (1 - b1)'
  dur_term:
    desc: 'duration of school terms'
    value: '365 - dur_holidays'
  dur_holidays:
    desc: 'duration of holidays'
    # value: 92
    value: 'duration_of_summer + duration_of_christmas + duration_of_easter + duration_of_autumn'
  beta0:
    desc: 'transmission rate (/day)'
    value: '1250 / 365'
```

```yaml
  corrected_beta0:
    desc: 'transmission rate (/day) corrected according to the proportion of holidays to ensure that the average
    transmission rate is beta0'
    value: 'beta0 * 365 / ((1 + b1) * dur_term + (1 - b1) * dur_holidays)'
  b1:
    desc: 'amplitude of the seasonality (0-1)'
    value: 0.025
  avg_dur_E:
    desc: 'average duration of the exposed state (days)'
    value: 8
  avg_dur_I:
    desc: 'average duration of the infectious state (days)'
    value: 5

initial_conditions:
  population:
    - population:
        - vars: [S]
          amount: 'initial_pop_size * prop_S'
        - vars: [E]
          amount: 'initial_pop_size * prop_EI'
        - vars: [I]
          amount: 'initial_pop_size * prop_EI'
        - vars: [R]
          amount: 'initial_pop_size * (1 - prop_EI - prop_S)'
        - vars: [V]
          amount: 0

outputs:
  type: csv
  population:
    period: 1
    extra_vars:
      - cum_incidence
...
```

### B Animal health: a model of a vector-borne disease, Rift Valley fever

#### Original model

This section reproduces a model of a vector-borne disease, the Rift Valley fever, initially published in Cavalerie *et al.*, "A Stochastic Model to Study Rift Valley Fever Persistence with Different Seasonal Patterns of Vector Abundance: New Insights on the Endemicity in the Tropical Island of Mayotte", *PLOS ONE* (2015), `DOI:10.1371/journal.pone.0130838`.

It is compartment-based, hosts infection following a SEIR process (states being denoted respectively by SH, EH, IH and RH in fig. 4), while vectors infection is described by a SEI dynamics (SV, EV and IV states) with two states for juvenile vectors in aquatic stage (susceptible and infected, respectively SA and IA). The emergence rate from juvenile to adult vectors, $\varphi(t)$, is environment-driven (modelled by a periodic function). Infection is introduced through a single infected host, one year after simulation begins.

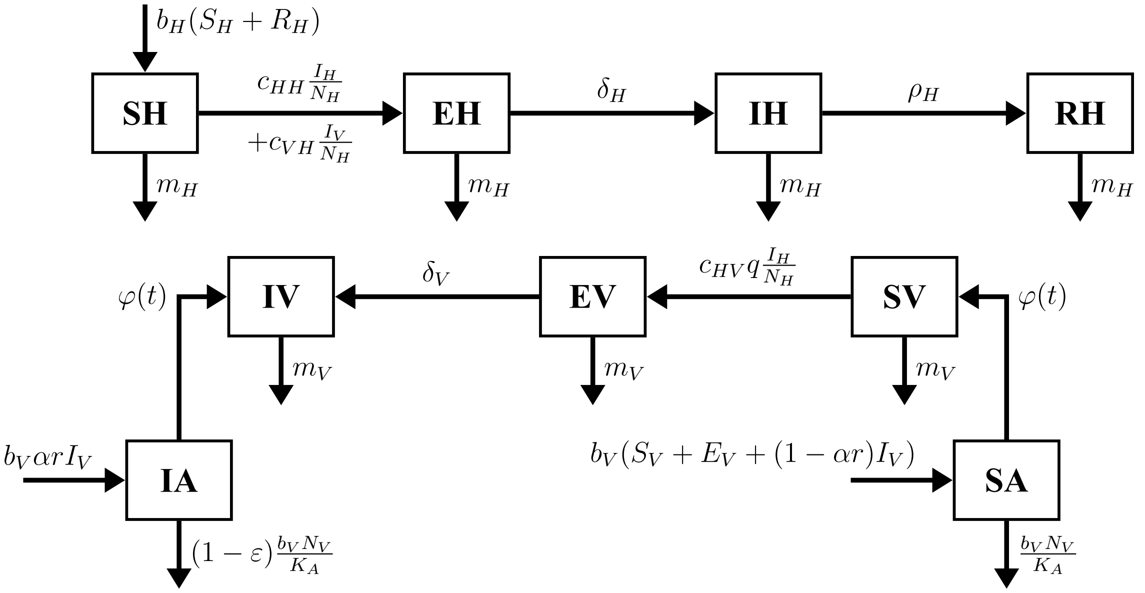


**Figure 4:** Flow diagram corresponding to the Rift Valley fever model (after Cavalerie *et al.* 2015) with compartments for hosts (top) and vectors (bottom). Parameters: $b_H, b_V, m_H, m_V$: birth/mortality rates for hosts and vectors, respectively; $N_H, N_V$: total population of hosts/vectors; $c_{HH}$: direct transmission rate; $c_{VH}, c_{HV}$: transmission probability from vector to host/host to vector; $1/\delta_H, 1/\delta_V$: duration of incubation in hosts/vectors; $1/\rho_H, 1/\rho_V$: duration of viraemia in hosts/vectors; $q$: biting rate; $K_A$: carrying capacity for aquatic stage; $\varepsilon$: proportion of infected eggs not affected by death; $\alpha$: transovarian transmission probability; $\varphi(t)$: emergence rate, environment-dependend periodic forcing function.

#### Implementation in EMULSION

This model was re-implemented with EMULSION as a stochastic compartment-based model (`rvf.yaml`, p. 11). The corresponding state machine diagram is provided on fig. 5.

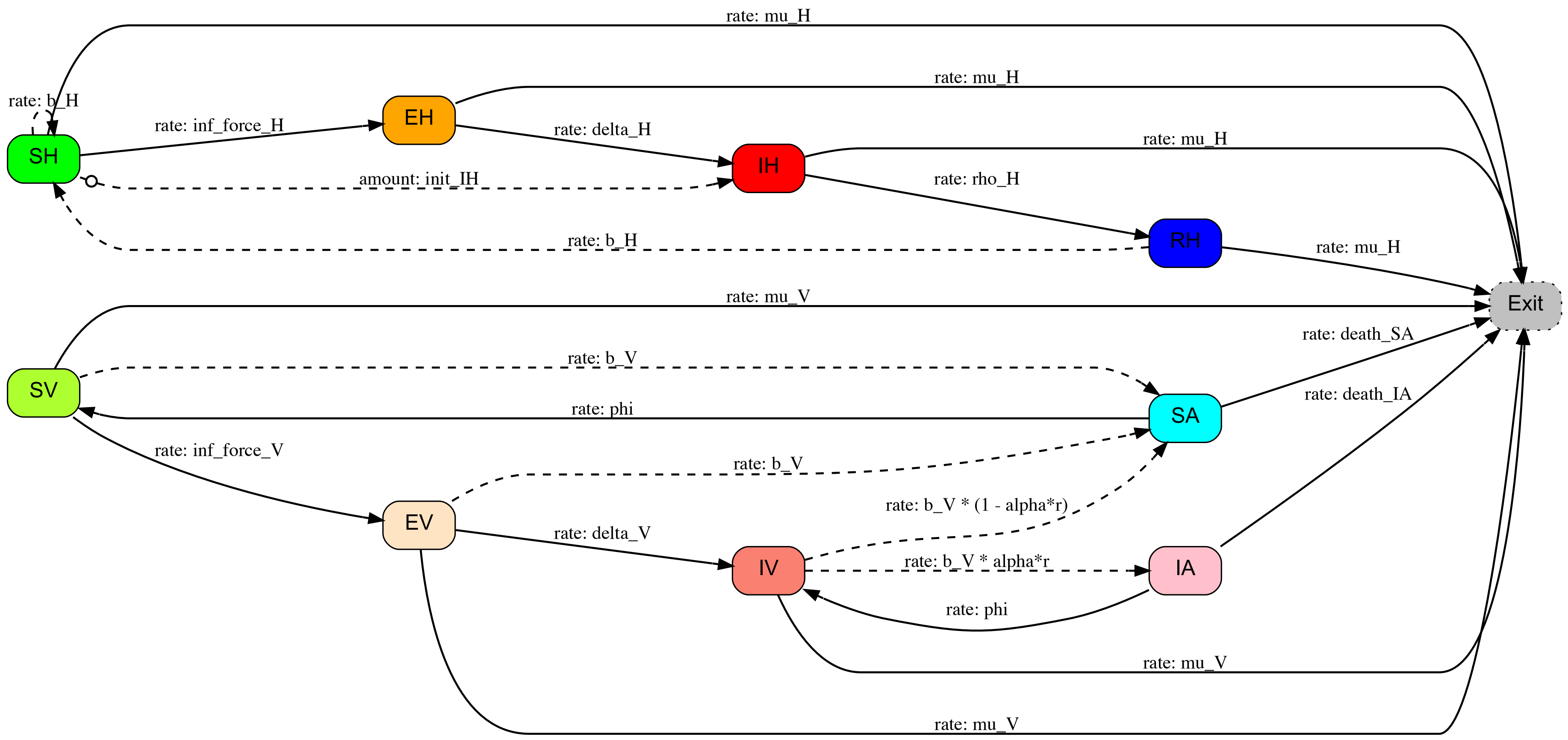


**Figure 5:** State machine diagram for the RVF model in EMULSION. Plain arrows denote transitions between states, while dashed arrows denote production links. Removal of individuals is made explicit through the "Exit" (deceased) state. The circle on the production link between HS and HI indicates a calendar condition (introduction of an infected host). This figure was produced as follows: `emulsion diagrams rvf.yaml --format pdf`

#### Reproduction of published results

The RVF model was run for 1000 stochastic repetitions using the forcing function defined as “scenario C” in Cavalerie *et al.* 2015. RVF dynamics as shown in the original article was fully reproduced with EMULSION (fig. 6).

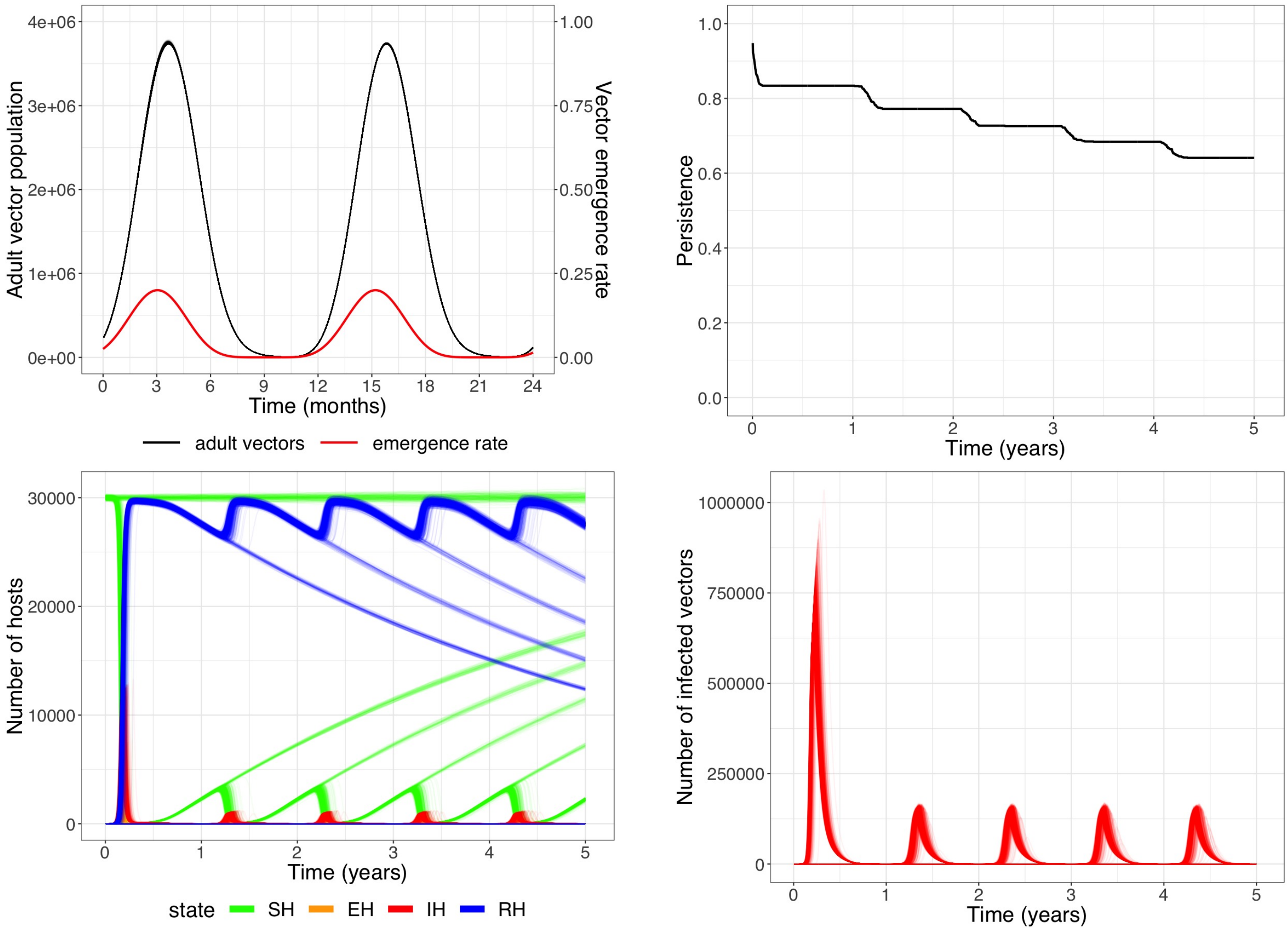


**Figure 6:** RVF dynamics (1000 stochastic repetitions). Top left: Population of adult vectors in the two first years after infection, compared to the forcing function. This figure is similar to Cavalerie *et al.* 2015, fig. 2 (scenario *c*). Top right: Persistence of infection over 5 years. Bottom left: Number of hosts in each state. Bottom right: Number of infected vectors. These figures are similar to Cavalerie *et al.* 2015, fig. 4 (scenario *c*). Parameters: see `rvf.yaml` below.

#### Notes on EMULSION features used for reimplementing the model

- Introduction of an infected host after 1 year is done using an explicit **calendar** with an event `disease_introduction` used to enable a specific production link which creates animals in compartment IH (lines 17–20 and 100–104)
- The seasonality of vector emergence (from aquatic to adult stage) is forced by a periodic function of time (lines 88–89 and 175–179)

#### Model in EMULSION DSL (`rvf.yaml`)

```yaml
---
model_name: RVF_compart

model_info:
  abstract: 'This model is a simple discrete-time, stochastic, compartment-based model for the Rift Valley fever
  (vector-borne disease) reproducing model in Cavalerie et al. 2015'
  author:
    - 'Sebastien Picault'
    - 'Pauline Ezanno'
  DOI: '10.1371/journal.pone.0130838'

time_info:
  time_unit: days
  delta_t: 1
  origin: 'November 1, 2006'
  total_duration: '6*365'
  calendars:
    disease:
      events:
        disease_introduction: {date: 'November 1, 2007'}

levels:
  herd:
    desc: level of the herd
    aggregation_type: compartment

processes:
  herd:
    - infection

grouping:
  herd:
    infection:
      machine_name: health_state
      key_variables: [health_state]

state_machines:
  health_state:
    desc: 'state machine describing health states of hosts and vectors'
    states:
      - SH:
          name: SusceptibleHost
          desc: 'Susceptible Hosts'
          fillcolor: 'green'
      - EH:
          name: ExposedHost
          desc: 'Exposed Hosts, infected but not yet able to infect vectors'
          fillcolor: 'orange'
      - IH:
          name: InfectiousHost
          desc: 'Infectious Hosts, infected and able to infect vectors'
          fillcolor: 'red'
      - RH:
          name: ImmuneHost
          desc: 'Immune Hosts, resistant to infection'
          fillcolor: 'blue'
      - SV:
          name: SusceptibleVector
          desc: 'Susceptible Vectors'
          fillcolor: 'greenyellow'
      - SA:
          name: SusceptibleVectorJuv
          desc: 'Susceptible Juvenile Vectors (aquatic stage)'
          fillcolor: 'cyan'
```

```
   - EV:
     name: ExposedVector
     desc: 'Exposed Vectors, infected but not yet able to infect hosts'
     fillcolor: 'bisque'
   - IV:
     name: InfectiousVector
     desc: 'Infectious Vectors, infected and able to infect hosts'
     fillcolor: 'salmon'
   - IA:
     name: InfectiousVectorJuv
     desc: 'Infectious Juvenile Vectors (aquatic stage)'
     fillcolor: 'pink'
   - Exit:
     name: DeadIndividuals
     desc: 'Artificial compartment for deceased hosts and vectors'
     autoremove: yes
     fillcolor: 'gray'
  transitions:
   - {from: SH, to: EH, rate: 'inf_force_H'}
   - {from: EH, to: IH, rate: 'delta_H'}
   - {from: IH, to: RH, rate: 'rho_H'}
   - {from: SV, to: EV, rate: 'inf_force_V'}
   - {from: EV, to: IV, rate: 'delta_V'}
   - {from: SA, to: SV, rate: 'phi'}
   - {from: IA, to: IV, rate: 'phi'}
   - {from: SH, to: Exit, rate: 'm_H'}
   - {from: EH, to: Exit, rate: 'm_H'}
   - {from: IH, to: Exit, rate: 'm_H'}
   - {from: RH, to: Exit, rate: 'm_H'}
   - {from: SV, to: Exit, rate: 'm_V'}
   - {from: EV, to: Exit, rate: 'm_V'}
   - {from: IV, to: Exit, rate: 'm_V'}
   - {from: SA, to: Exit, rate: 'death_SA'}
   - {from: IA, to: Exit, rate: 'death_IA'}
  productions:
   - from: SH
    to: IH
    amount: 'init_IH'
    when: 'disease_introduction'
    desc: 'introduction of the disease after 1 year'
   - {from: SH, to: SH, rate: 'b_H'}
   - {from: RH, to: SH, rate: 'b_H'}
   - {from: SV, to: SA, rate: 'b_V'}
   - {from: EV, to: SA, rate: 'b_V'}
   - {from: IV, to: SA, rate: 'b_V * (1 - alpha*r)'}
   - {from: IV, to: IA, rate: 'b_V * alpha*r'}

parameters:
 death_SA:
  desc: 'death rate among SA (density-dependance in aquatic stage)'
  value: 'b_V * (total_SV + total_EV + total_IV) / K_A'
 death_IA:
  desc:  'death rate among IA (density-dependance in aquatic stage)'
  value: 'death_SA * (1 - epsilon)'
 inf_force_H:
  desc: 'force of infection experienced by susceptible hosts'
  value: ' (c_vh * q * total_IV  + c_hh * total_IH)/ total_H'
 inf_force_V:
  desc: 'force of infection experienced by susceptible adult vectors'
  value: 'c_hv * q * total_IH / total_H'
 epsilon:
  desc: 'proportion of infected eggs not affected by death among aquatic stages'
  value: 0.44
 K_A:
  desc: 'carrying capacity in juvenile vectors'
  value: 1000000
```

```yaml
  alpha:
    desc: 'transovarian transmission probability'
    value: 1/279
  r:
    desc: 'proportion of Aedes in the vector population (to our best knowledge) - All vectors give birth to S individu
    excepted part of infected Aedes (vertical transmission)'
    value: 0.5
  c_vh:
    desc: 'transmission probability from infected vector to susceptible host'
    value: 0.4
  c_hv:
    desc: 'transmission probability from infected host to susceptible vector'
    value: 0.6
  q:
    desc: 'biting rate'
    value: 0.25
  c_hh:
    desc: 'rate of direct transmission among hosts'
    value: '1/1000'
  delta_H:
    desc: '1/incubation period'
    value: '1/2'
  rho_H:
    desc: 'rate of immunity acquisition'
    value: '1/6'
  delta_V:
    desc: '1/EIP (extrinsinc incubation period)'
    value: '1/6'
  b_H:
    desc: 'daily host birth rate'
    value: '1/(5*365)'
  b_V:
    desc: 'daily vector renewal rate'
    value: 4
  m_H:
    desc: 'daily host mortality rate'
    value: '1/(5*365)'
  m_V:
    desc: 'daily vector mortality rate'
    value: '1/20'
  theta:
    desc: 'minimum development time before emergence in optimal
    conditions'
    value: 5
  phi:
    desc: 'emergence rate, forcing function based on the time elapsed since beginning of simulation (using variable t
    provided by EMULSION DSL)'
    value: ' (((sin(2 * pi * time / 365) + 1) / 2)**3) / theta'
    source: 'Cavalerie et al 2015, scenario C'
  init_H:
    desc: 'initial host population size'
    value: 30000
  init_IH:
    desc: 'initial number of infected hosts'
    value: 1
  init_V:
    desc: 'initial vector population size'
    value: 1000
  total_H:
    desc: 'total number of hosts'
    value: 'total_SH + total_EH + total_IH + total_RH'

initial_conditions:
  herd:
    - population:
        - total: 'init_H + init_V'
```

```
        - vars: [SH]
          amount: 'init_H'
        - vars: [SV]
          amount: 'init_V'

outputs:
  type: csv
  herd:
    period: 1
    extra_vars:
      - phi
...
```

### C Plant health: a spatially explicit individual-based model of Bahia Bark Scaling of Citrus spread and control

#### Original model

This section reproduces a spatially explicit model of plant disease, the Bahia Bark Scaling of Citrus (BBSC), originally published in Cunniffe *et al.*, "Cost-Effective Control of Plant Disease When Epidemiological Knowledge is Incomplete: Modelling Bahia Bark Scaling of Citrus", *PLOS Computational Biology* (2014), `DOI:10.1371/journal.pcbi.1003753`. This model is individual-based, each tree experiencing a SEIR infection process described by the flow diagram below (fig 7):

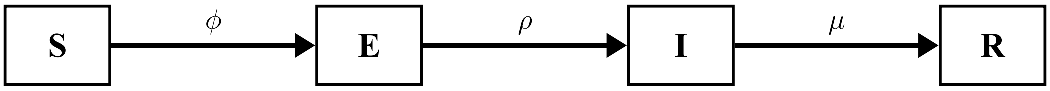


**Figure 7:** Flow diagram corresponding to SEIR model of the BBSC (after Cunniffe *et al.* 2014). R state represent removed hosts. Parameters: $\phi$: infection rate experienced by susceptible hosts; $1/\rho$: average duration of latent period; $\mu$: mortality rate (assumed 0 in the model).

Trees are planted in groves composed of several rows. They are all introduced at the same age, assuming this initial plantation as the only source of disease introduction (through E plants). Additional assumptions are not represented in the flow diagram:

- trees planted in the grove are juvenile hosts, which cannot infect other hosts nor be infected by infected hosts; they become "epidemiologically competent" (adult) after two years
- natural mortality is not modelled, infected trees being removed (R state) only as a consequence of a periodic scouting and roguing, with a given probability of dectection during the scouting campaign (thus $\mu = 0$)

Besides, disease spread is controlled by a dispersal kernel:

$$K(d;\alpha) = \frac{\exp(\frac{-d}{\alpha})}{2\pi\alpha^2}$$

so that infection rate experienced by host $i$ due to all infected hosts $j \in \Omega_I$ is:

$$\phi_i = \beta \sum_{j\in\Omega_I} K(d_{ij};\alpha)$$

#### Implementation in EMULSION

This model was re-implemented with EMULSION as an individual-based model (`bbsc.yaml`, p. 19). The corresponding state machine diagram is provided on fig. 8). Since EMULSION does not provide built-in functions or processes for spatially explicit models yet, a small Python code add-on was written to initialize tree positions and distances and compute the contribution of infected hosts using the dispersal kernel (`bbsc.py`, p. 22). For the sake of simplicity we focused on one grove, neglecting neighbouring trees from adjacent groves.

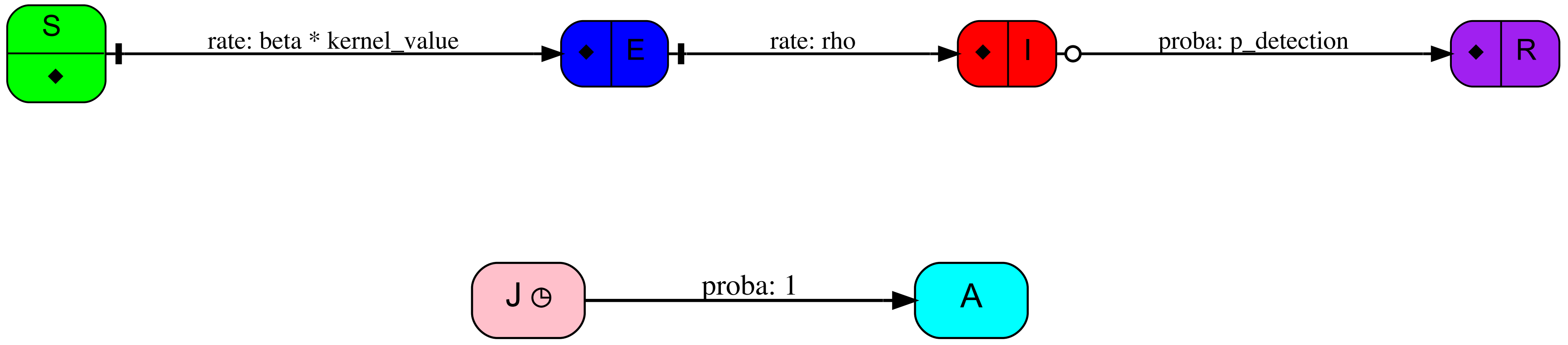


**Figure 8:** State machine diagrams for the BBSC model in EMULSION (top: health states, bottom: maturity). Plain arrows denote transitions between states, while dashed arrows denote production links. The circle on the transitions from I to R indicates a calendar condition (scouting events) and allow I individuals to become R with a probability `p_detection`. Vertical bars on the transitions from S to E and from E to I signals a condition (being adult). The clock in J state indicates that a specific duration distribution is associated with this state, before which individuals cannot leave the J state. These figures were produced as follows: `emulsion diagrams bbsc.yaml --format pdf`

#### Reproduction of published results

Simulations were run to compare uncontrolled disease spread at grove scale and the impact of control measures, consisting in a periodic scouting with a given probability of detecting infected trees, followed by immediate roguing. Removed trees are not replanted.

Figure 9 shows the evolution of asymptomatic trees (susceptible or exposed) over 20 years, as in Cunniffe `et al.` 2014, fig. 1b. The spatial spread is displayed for one stochastic repetition of each scenario (fig. 10, similar to fig. 1c in original article).

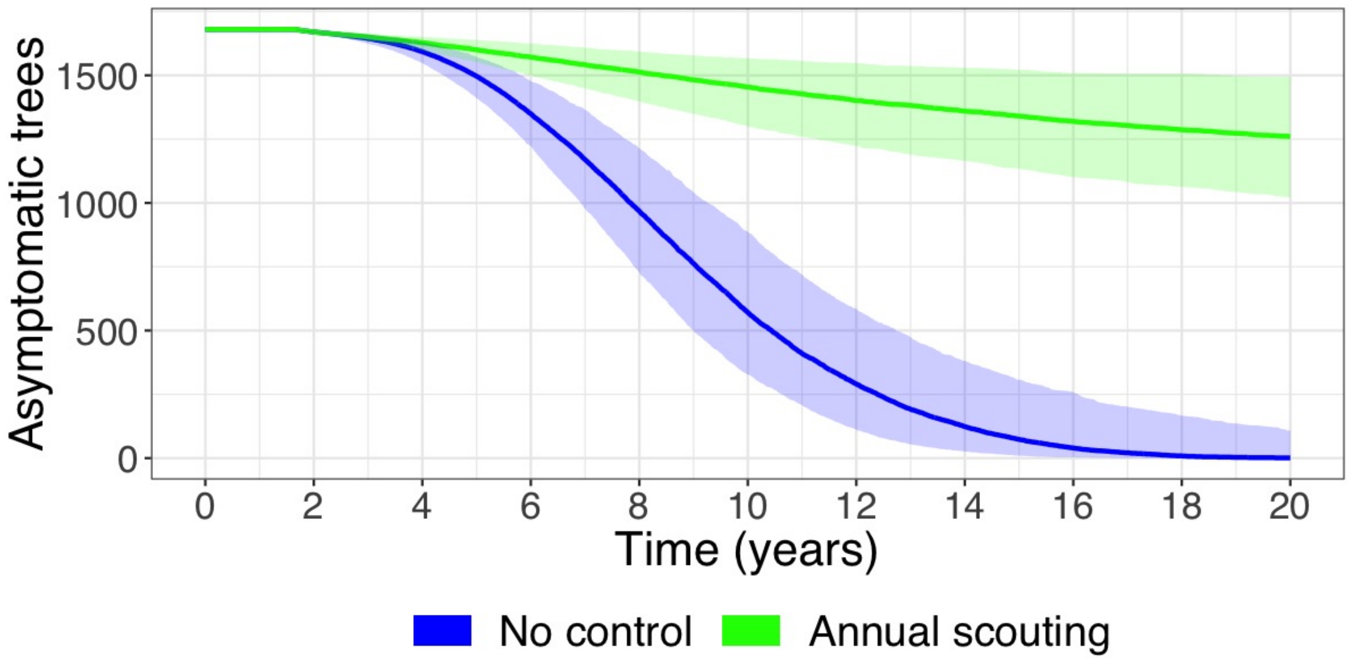


**Figure 9:** Evolution of the amount of asymptomatic (S + E) trees (median and 90% credibility interval, 500 stochastic repetitions) without control (blue) or with annual scouting and roguing (green). Parameters: see `bbsc.yaml`

#### Notes on EMULSION features used for reimplementing the model

- Periodic scouting relies upon an explicit **calendar** (lines 16–20) and a `when` keyword in I to R transition (lines 67–74)
- The link between the model and a Python code add-on is made through the definition of levels, which are explicitly associated with a Python class and a file (lines 22-33)

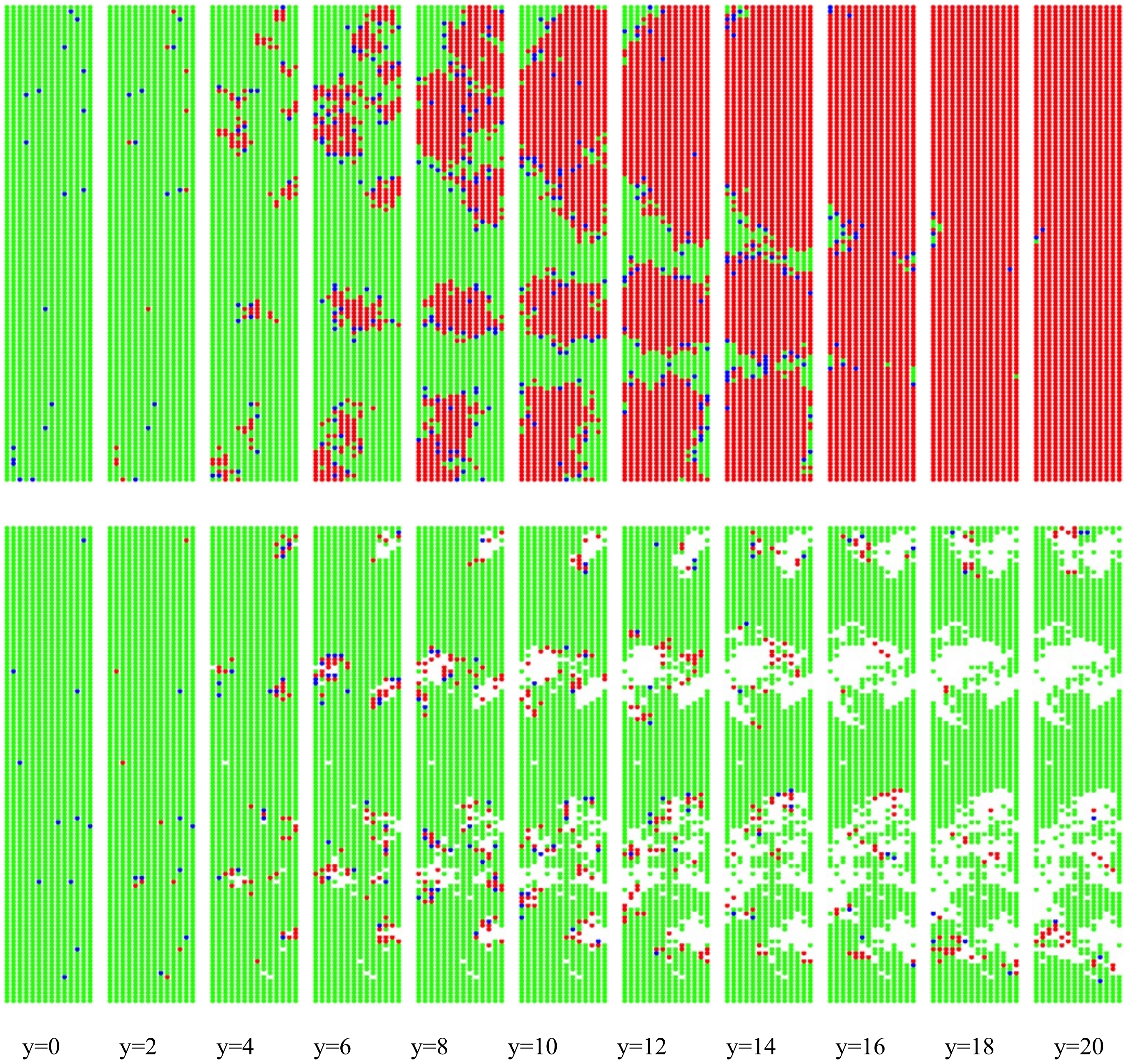


**Figure 10:** BBSC spread in a grove over 20 years (one image every 2 years) in one stochastic repetition for each scenario (top: no control; bottom: annual scouting and roguing). Each dot represents a tree, the colour indicates its health state as defined in state machine diagram on fig. 8, except for trees removed by roguing which are also removed from the image.

.

#### Model in EMULSION DSL (bbsc.yaml)

```yaml
---
model_name: IBM_BBSC

model_info:
  abstract: 'This model is a simple discrete-time, stochastic,
  individual-based SEIR model for Bahia Bark Scaling of Citrus (BBSC)
  reproducing model in Cunniffe et al 2014'
  author: 'Sebastien Picault'
  DOI: '10.1371/journal.pcbi.1003753'

time_info:
  time_unit: 'weeks'
  delta_t: 1
  origin: 'January 1'
  total_duration: '20*52'
  calendars:
    control:
      period: {weeks: 52}
      events:
        scouting: {date: 'January 31'}

levels:
  grove:
    desc: 'Level of the spatialized population'
    aggregation_type: 'IBM'
    contains:
      - trees
    file: bbsc.py
    class_name: Grove
  trees:
    desc: 'Level of the individual plants'
    file: bbsc.py
    class_name: Tree

processes:
  grove:
    # Python process to update vector tree_health_status (vector V
    # such that V[i] = 1 if tree i is infectious, 0 otherwise). Done
    # once at the beginning of each time step, used then by each tree
    # in action 'compute_kernel'
    - compute_tree_health_status
  trees:
    - maturity
    - health_state

state_machines:
  health_state:
    desc: 'The state machine which defines the evolution of health
    states
    '
    states:
      - S:
          name: 'Susceptible'
          desc: 'uninfected'
          fillcolor: 'green'
          on_stay:
            - action: compute_kernel
      - E:
          name: 'Exposed'
          desc: 'latently infected, neither symptomatic nor infectious'
          fillcolor: 'blue'
          on_enter:
            - log_vars: [is_E]
      - I:
```

```yaml
          name: 'Infectious'
          desc: 'both infectious and symptomatic'
          fillcolor: 'red'
          on_enter:
            - log_vars: [is_I]
      - R:
          name: 'Removed'
          desc: 'removed by control'
          fillcolor: 'purple'
          on_enter:
            - log_vars: [is_R]
    transitions:
      - from: S
        to: E
        rate: 'beta * kernel_value'
        cond: is_A
      - from: E
        to: I
        rate: 'rho'
        cond: is_A
      - when: scouting
        from: I
        to: R
        proba: 'p_detection'
  maturity:
    desc: 'The state machine which defines the evolution of hosts from
    immaturity to epidemiological maturity'
    states:
      - J:
          name: 'Juveniles'
          desc: 'juveniles that cannot become infected or transmit infection'
          fillcolor: 'pink'
          duration: 'delta'
      - A:
          name: 'Adult trees'
          desc: 'adult trees that are epidemiologically competent'
          fillcolor: 'cyan'
    transitions:
      - {from: J, to: A, proba: 1}

parameters:
  rho:
    desc: 'rate of onset of infectiousness/symptoms'
    value: '0.185 / 4'
    # value: '0.135 / 4'
    source: 'sampled in [0.135, 0.235] per month in original paper'
  alpha:
    desc: 'dispersal scale (m)'
    value: 2.585
    # value: 1.96
    source: 'sampled in [1.96, 3.21] m in original paper'
  beta:
    desc: 'rate of infection (m2/week)'
    value: '5.05 / 4'
    # value: '2.79 / 4'
    source: 'sampled in [2.79, 7.31] m2/month in original paper'
  delta:
    desc: 'delay before epidemiological maturity (weeks)'
    value: '21.65 * 4'
    # value: '25.4 * 4'
    source: 'sampled in [17.9, 25.4] month in original paper'
  p_detection:
    desc: 'probability of detecting an infected tree during scouting'
    value: 0.6
  grove_width:
    desc: 'number of rows in the grove'
```

```
    value: 14
    # value: 2
  grove_length:
    desc: 'number of trees per row'
    value: 120
    # value: 4
  between_tree_space:
    desc: 'distance between trees within the same row (m)'
    value: 4
  between_row_space:
    desc: 'distance between adjacent rows (m)'
    value: 'between_tree_space * 3 / 2'
  initial_grove_size:
    desc: 'initial number of trees in the grove'
    value: 'grove_width * grove_length'
  E0:
    desc: 'initial proportion of exposed immature trees in the grove'
    value: 0.01
  asymptomatic:
    desc: 'number of asymptomatic trees'
    value: 'total_S + total_E'

statevars:
  kernel_value:
    desc: 'variable which contains the result of the summation over
    infectious hosts based on the exponential kernel, calculated by
    action compute_kernel, and used to decide whether the host becomes
    infected or not'
  tree_health_status:
    desc: 'vector V such that V[i] = 1 if tree i is infectious, 0 otherwise'

actions:
  compute_kernel:
    desc: 'Python-defined function to make a susceptible host compute the summation
    over infectious hosts based on the exponential kernel. The result is stored in
    a variable named kernel_value'


prototypes:
  trees:
    - juvenile_tree:
        desc: 'juvenile plant, with a health state determined by the
        initial prevalence E0'
        health_state: 'random(1-E0, E0, 0, 0)'
        maturity: J

initial_conditions:
  grove:
    - prototype: juvenile_tree
      amount: 'initial_grove_size'

outputs:
  type: csv
  grove:
    period: 1
    extra_vars:
      - scouting
      - asymptomatic
...
```

##### Python code add-on for spatial computations (`bbsc.py`)

```
"""This file is aimed at providing specific code add-on for the BBSC
model (bbsc.yaml). It is based on the code skeletton generated
automatically by EMULSION (command: emulsion generate bbsc.yaml).

"""

import numpy as np
from emulsion.agent.managers import IBMProcessManager
from emulsion.agent.views import SimpleView
from emulsion.agent.atoms import EvolvingAtom


#===============================================================
# CLASS Grove (LEVEL 'grove')
#===============================================================
class Grove(IBMProcessManager):
    """Level of the spatial population."""

    #-----------------------------------------------------------
    # Level initialization: operations performed when creating a Grove
    #-----------------------------------------------------------

    def initialize_level(self, **others):
        """Initialize a grove."""
        # retrieve environment size and store it in grove's variables
        self.statevars.width = int(self.get_model_value('grove_width'))
        self.statevars.length = int(self.get_model_value('grove_length'))

        # init variable tree_health_status: a vector V such that V[i] = 1
        # if tree i is infectious, 0 otherwise
        # this vector is stored in the statevars of the grove agent
        self.statevars.tree_health_status = np.zeros(self.statevars.width * self.statevars.length)

    #-----------------------------------------------------------
    # Processes
    #-----------------------------------------------------------

    def compute_tree_health_status(self):
        """Update vector tree_health_status V[i] = 1 if tree i is infectious, 0 otherwise

        """
        # iterate other trees contained in this grove
        for tree in self.select_atoms():
            # compute tree ID (agent ID modulo the size of the grove)
            tree_ID = (tree.agid - 1) % (self.statevars.width * self.statevars.length)
            # associate the tree ID with its status (1 if I, 0 otherwise)
            self.statevars.tree_health_status[tree_ID] = tree.is_I

#===============================================================
# CLASS Tree (LEVEL 'trees')
#===============================================================
class Tree(EvolvingAtom):
    """Level of the individual plants."""

    #-----------------------------------------------------------
    # Actions (called from bbsc.yaml)
    #-----------------------------------------------------------

    def compute_kernel(self, *args, **kwargs):
        """Python-defined function to make a susceptible host compute the
        summation over infectious hosts based on the exponential
        kernel. The result is stored in a variable named kernel_value.

        """
```

```
        # if first usage, kernel term with for all other neighbours
        if 'K_values' not in self.statevars:
            # retrieve parameter values
            width = int(self.get_model_value('grove_width'))
            length = int(self.get_model_value('grove_length'))
            alpha = self.get_model_value('alpha')
            dist_x = self.get_model_value('between_row_space')
            dist_y = self.get_model_value('between_tree_space')
            # compute coords of current tree
            my_tree_id = (self.agid - 1) % (width * length)
            my_row, my_col = my_tree_id // length, my_tree_id % length
            self.statevars.K_values = np.zeros(width * length)
            # compute kernel values according to distances
            for tree_id in range(width * length):
                if tree_id == my_tree_id:
                    self.statevars.K_values[tree_id] = 0
                else:
                    row, col = tree_id // length, tree_id % length
                    dist = np.sqrt(((row - my_row) * dist_x)**2 + ((col - my_col) * dist_y)**2)
                    self.statevars.K_values[tree_id] = np.exp(-dist / alpha) / (2 * np.pi * alpha**2)
            # initialize kernel value to zero
            self.statevars.kernel_value = 0
        # otherwise, compute the dot product between K_values and the health states of other agents
        else:
            self.statevars.kernel_value = np.dot(self.statevars.K_values,
                                                 self.upper_level().statevars.tree_health_status)
```
