## Supplementary material for "EMULSION: transparent and flexible multiscale stochastic models in human, animal and plant epidemiology": S6 Text

### S6 Text – Installation instructions for EMULSION

**D Test EMULSION installation** **3**

**E Main command-line options for EMULSION** **6**

This document provides instructions for installing EMULSION, testing the installation, and describes the main commands and options available for the `emulsion` command. The latest version of EMULSION and of the documentation is available on: `https://sourcesup.renater.fr/www/emulsion-public`

#### A Requirements

**System** EMULSION has been designed under **MacOS** and **Linux**, and also works with **Windows 10**, with minor limitations such as the absence of shell completion. Installation procedures are specific to each system.
Also, in what follows, we assume that MacOS and Linux users are working with a **bash** shell, which often is the default configuration. To check this point, typing: `echo $SHELL` should print: `/bin/bash`. Otherwise, the commands below may require some adjustments (please refer to the documentation of your shell).

**Language** EMULSION is written in **Python 3** (version 3.6 or higher). In what follows, we assume that `python3` and `pip3` refer to the Python3 installation. If not, they should be replaced in the commands below by the proper names (e.g. with Windows: respectively `python.exe` and `pip.exe`).
The installation of Python 3 is fully described on Python website (`https://www.python.org/downloads/`). When installing on Windows, the box ”Add Python 3.x to PATH” must be checked to make python commands available from the terminal.

#### B Install EMULSION in the Python environment

##### B.1 Linux and MacOS

Open a terminal and type the following commands:

```
sudo pip3 install emulsion
init_emulsion
source $HOME/.bashrc
```

Notes:

1. Depending on how Python was installed on your system, `sudo` may be optional.
2. The second command (`init_emulsion`) initializes command-line completion (available with a bash shell, under Linux or MacOS), which allows to use `TAB` key to get suggestions on what is expected (options, files, parameters...) in the command. It also creates (or modifies) the environment variable named `PYTHONPATH` to make EMULSION able to find code add-ons located in the same directory as your models.

##### B.2 Windows

Open a terminal (either ”Command Prompt” also known as `cmd.exe`, or ”Windows Power Shell”), then type:

```
pip.exe install emulsion
```

To make EMULSION able to find code add-ons in the directory where your model files are located, you must declare (or modify) an environment variable named `PYTHONPATH` to add the current directory (“`.`”).

To do so, you can use the graphical interface (from System Preferences), or the terminal. The syntax depends on the kind of terminal you are using:

- with the ”Command Prompt” (`cmd.exe`), type:

  ```
  setx PYTHONPATH ".;%PYTHONPATH%"
  ```

- with ”Windows Power Shell”, type:

  ```
  setx PYTHONPATH ".;$Env:PYTHONPATH"
  ```

#### C Install Graphviz (for state machine diagrams)

Graphviz (`http://www.graphviz.org`) is highly recommended (otherwise, simulations will run but the diagrams representing model structure will not be produced).

##### C.1 Linux

```
sudo apt install graphviz
```

##### C.2 MacOS

Graphviz can be installed on MacOS using for instance **homebrew** (`https://brew.sh/`):

```
brew install graphviz
```

##### C.3 Windows

To install on Windows, download the latest stable version from Graphviz website (`https://graphviz.gitlab.io/_pages/Download/Download_windows.html`) and update the `PATH` environment variable according to the location of executable files are (especially `dot.exe`).
For instance, if `dot.exe` is located in `C:\Program Files (x86)\Graphviz\bin`:

- using the “Command Prompt” (`cmd.exe`):

  ```
  setx PATH "C:\Program Files (x86)\Graphviz\bin;%PATH%"
  ```

- using “Windows Power Shell”:

  ```
  setx PATH "C:\Program Files (x86)\Graphviz\bin;$Env:PATH"
  ```

### D Test EMULSION installation

1. Download model examples: `https://sourcesup.renater.fr/www/emulsion-public/models.zip`

2. Extract the archive, open a terminal in the `models` directory, then type:

```
cd quickstart
emulsion run --plot quickstart.yaml --view-model --silent
```

3. This should produce the following output in the terminal:

```
Simulation level:herd
Generated state machine diagram img/Quickstart_age_group_machine.svg
Generated state machine diagram img/Quickstart_life_cycle_machine.svg
Generated state machine diagram img/Quickstart_health_state_machine.svg
100%|**************************************************************| 10/10
Simulation finished in 12.37 s
Outputs stored in outputs/counts.csv
Outputs plot in file: img/Quickstart.html
```

4. The default web navigator should open and display the following figures. They corresponds, for each state machine defined in the model, to the state machine diagram and the amounts of individuals in each state over time for each stochastic repetition, plus additional variables specified as `extra_vars` in the model file. All these outputs are stored in a file named `counts.csv` in the output directory (by default, `outputs/`).

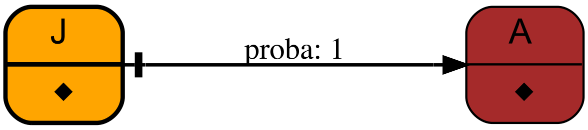

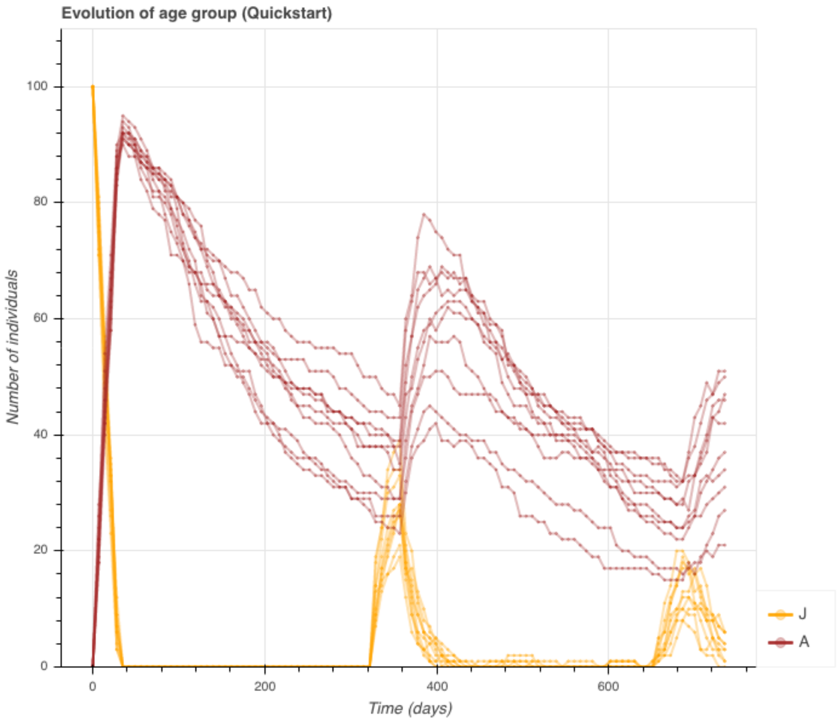

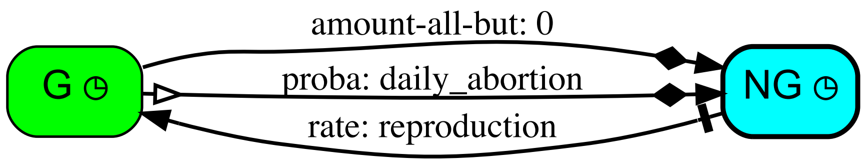
amount-all-but: 0
proba: daily_abortion
rate: reproduction
G
NG

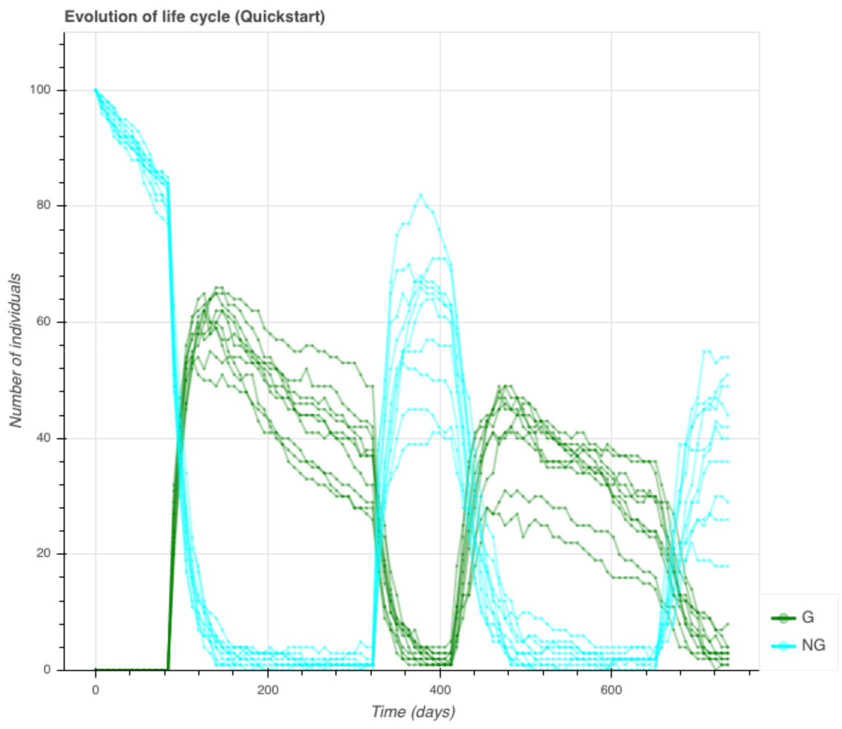
Evolution of life cycle (Quickstart)
Number of individuals
Time (days)
0
20
40
60
80
100
0
200
400
600
G
NG

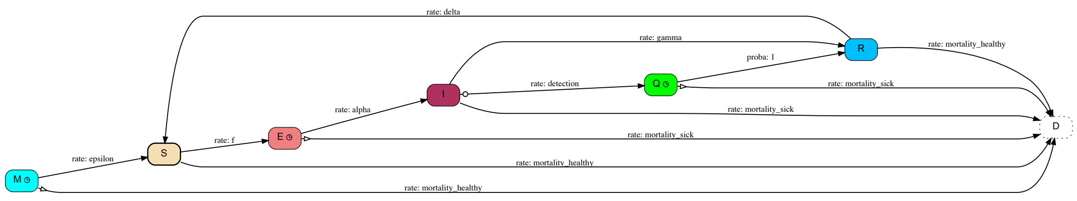
rate: delta
rate: gamma
proba: 1
rate: mortality_healthy
rate: detection
rate: mortality_sick
rate: alpha
rate: mortality_sick
rate: f
rate: mortality_sick
rate: epsilon
rate: mortality_healthy
rate: mortality_healthy
M
S
E
I
Q
R
D

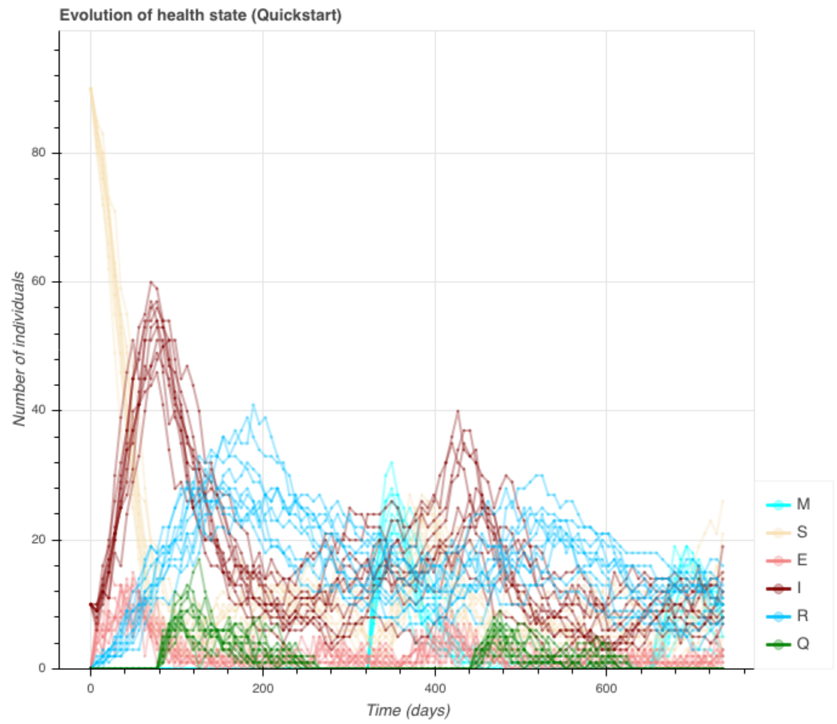
Evolution of health state (Quickstart)
Number of individuals
Time (days)
0
20
40
60
80
0
200
400
600
M
S
E
I
R
Q

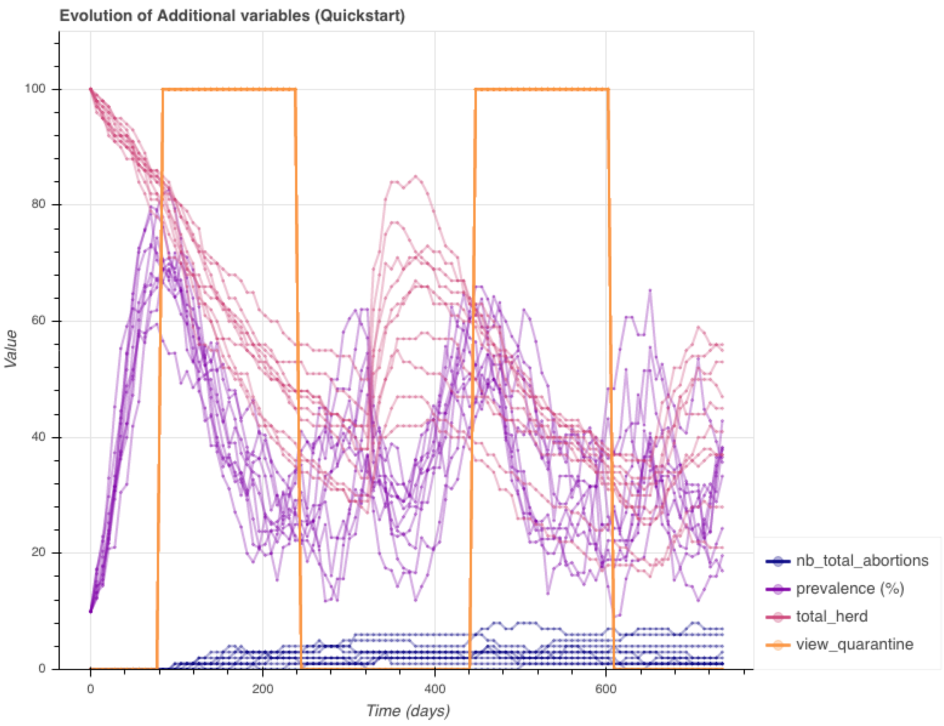
Evolution of Additional variables (Quickstart)
Value
Time (days)
0
20
40
60
80
100
0
200
400
600
nb_total_abortions
prevalence (%)
total_herd
view_quarantine

### E Main command-line options for EMULSION

This is a summary of the most frequently used commands and options for EMULSION. To get all available features, type: `emulsion -h`

```
Usage:
    emulsion run [--plot] MODEL [options] [(-p KEY=VALUE)...]
    emulsion diagrams MODEL
    emulsion show MODEL [options] [(-p KEY=VALUE)...]
    emulsion describe MODEL PARAM...
    emulsion plot MODEL [options]
    emulsion (-h | --help | -V | --version)

Commands:
  run MODEL                 Run simulations based on the specified MODEL (MODEL is
                            the path to the YAML file describing the model to run).
  diagrams MODEL            Generate diagrams for state machines defined in MODEL
                            (without running simulations), open them and exit.
  show MODEL                Print all MODEL parameter values and exit.
  describe MODEL PARAM...   Describe the role of specified PARAMeters in the
                            MODEL and exit.
  plot MODEL                Plot outputs for MODEL (assumed already run) and exit.

Options:
  -h --help                 Display this help page and exit.
  -V --version              Display version number and exit.
  --plot                    Plot outputs just after running the model.
  -r RUNS --runs RUNS       Specify the number of repetitions of the same model
                            [default: 10].
  -t STEPS --time STEPS     Specify the number of time steps to run in each
                            repetition. If the model defines a total_duration,
                            it is used as time limit, unless the '-t' option
                            is explicitly specified. Otherwise, the default
                            value is 100 steps.
  -p KEY=VAL                Change parameter named KEY to the specified VALue.
  --view-model              Produce diagrams to represent the state machines of
                            the model (requires Graphviz). Figures are stored in
                            figure-dir.
  --silent                  Show only the progression of repetitions instead
                            of the progression of each simulation.
  --output-dir OUTPUT       Specify a directory for simulation outputs
                            [default: outputs].
  --input-dir INPUT         Specify a directory for simulation inputs
                            [default: data].
  --figure-dir FIG          Specify a directory for graphic outputs (figures)
                            [default: img].
  --log-params              When producing CSV outputs, insert the name and value
                            of each parameter explicitly changed by option -p.
```
