## Supplementary material for "EMULSION: transparent and flexible multiscale stochastic models in human, animal and plant epidemiology": S1 File: bbsc.R

```
library(ggplot2)
library(tidyr)
library(dplyr)

args = commandArgs(trailingOnly=TRUE)

width = as.integer(args[1])
length = as.integer(args[2])
between_trees = as.integer(args[3])
between_rows = as.integer(args[4])
simu = as.integer(args[5])

theme_EMULSION <- theme_bw() +
    theme(text=element_text(family="Helvetica", size=20),
          axis.text=element_text(family="Helvetica", size=16),
          legend.text=element_text(family="Helvetica", size=18),
          )


## overview of states

t <- read.table("bbsc-counts-nocontrol.csv", sep=",", header=TRUE)
ts <- read.table("bbsc-counts-scout.csv", sep=",", header=TRUE)

REPET = max(t$simu_id) - min(t$simu_id) + 1

states <- rbind(t %>%
                select(-A, -J) %>%
                gather("state", "nb", 2:5) %>%
                mutate(scouting=FALSE),
                ts %>%
                select(-A, -J) %>%
                gather("state", "nb", 2:5) %>%
                mutate(scouting=TRUE))
states$state <- factor(states$state, levels=c("S", "E", "I", "R"))

p <- ggplot(states, aes(x=step/52, y=nb, group=simu_id, colour=state)) +
    geom_line(aes(group=interaction(state,simu_id,scouting), linetype=scouting),
              alpha=min(1,5/REPET)) +
    xlab("Time (years)") + ylab("Number of trees") +
    scale_x_continuous(breaks=seq(0,20,2), limits=c(0,NA)) +
    geom_vline(data=ts %>% filter(scouting=="True"), aes(xintercept=step/52),
               size=2, linetype="dashed", colour="gray", alpha=0.5) +
    guides(colour=guide_legend(override.aes = list(size=2, alpha=1),
                               keywidth=2)) +
    scale_colour_manual(values=c("S"="green", "E"="blue", "I"="red", "R"="purple")) +
    theme_EMULSION +
    theme(legend.position="bottom", legend.direction="horizontal")


asym <- rbind(t %>%
              group_by(step) %>%
              summarise(med=median(asymptomatic),
                        q5=quantile(asymptomatic, probs=0.05),
                        q95=quantile(asymptomatic, probs=0.95)) %>%
              mutate(scout="No control"),
              ts %>%
              group_by(step) %>%
              summarise(med=median(asymptomatic),
                        q5=quantile(asymptomatic, probs=0.05),
                        q95=quantile(asymptomatic, probs=0.95)) %>%
              mutate(scout="Annual scouting")
              )
asym$scout = factor(asym$scout, levels=c("No control", "Annual scouting"))

p <- ggplot(asym, aes(x=step/52, colour=scout)) +
    geom_line(aes(y=med), size=1) +
    geom_ribbon(aes(ymin=q5,ymax=q95, fill=scout), colour=FALSE, alpha=0.2) +
    xlab("Time (years)") + ylab("Asymptomatic trees") +
    scale_x_continuous(breaks=seq(0,20,2), limits=c(0,NA)) +
    guides(colour=FALSE,
           fill=guide_legend(title="",
                               override.aes = list(size=2, alpha=1),
                               keywidth=2)) +
    scale_colour_manual(values=c("Annual scouting"="green", "No control"="blue")) +
    scale_fill_manual(values=c("Annual scouting"="green", "No control"="blue")) +
    theme_EMULSION +
    theme(legend.position="bottom", legend.direction="horizontal")

jpeg("bbsc-asymptomatic.jpg", width=1200, height=600, quality=100, res=150)
print(p)
dev.off()


## visualize one grove for default
log <- read.table("bbsc-log-nocontrol.csv", sep=",",
                  col.names=c("simu_id", "step", "class", "tree_id", "state"))
log$state <- factor(log$state, levels=c("S", "E", "I", "R"))
logs <- read.table("bbsc-log-scout.csv", sep=",",
                  col.names=c("simu_id", "step", "class", "tree_id", "state"))
logs$state <- factor(logs$state, levels=c("S", "E", "I", "R"))

nb_trees = width*length
tree_ids = data.frame(tree_id = 1:nb_trees)

tree_row <- function (id) {
    ((id - 1) %% nb_trees) %% length
}

tree_col <- function (id) {
    ((id - 1) %% nb_trees) %/% length
}

for (t in 2*(0:10)) {
    history <- log %>%
        filter(simu_id == simu, step <= t*52) %>%
        select(-simu_id, -class)
    max_step <- history %>%
        group_by(tree_id) %>%
        summarise(step=max(step))
    latest <- history %>%
        inner_join(max_step, by=c("tree_id", "step")) %>%
        right_join(tree_ids, by=c("tree_id")) %>%
        mutate(x=tree_col(tree_id), y=tree_row(tree_id))
    latest$state[is.na(latest$state)] = "S"

p <- ggplot(latest, aes(x*between_rows,
                  y*between_trees,
                  colour=state)) +
    geom_point(size=0.1) +
    theme(aspect.ratio=1) +
    guides(colour=FALSE) +
    scale_colour_manual(values=c("S"="green", "E"="blue", "I"="red", "R"="white")) +
    theme_void()

    jpeg(paste("bbsc-nocontrol-", t, ".jpg", sep=""),
         width=width*between_rows, height=length*between_trees,
         quality=100, res=200)
    print(p)
    dev.off()

    history <- logs %>%
        filter(simu_id == simu, step <= t*52) %>%
        select(-simu_id, -class)
    max_step <- history %>%
        group_by(tree_id) %>%
        summarise(step=max(step))
    latest <- history %>%
        inner_join(max_step, by=c("tree_id", "step")) %>%
        right_join(tree_ids, by=c("tree_id")) %>%
        mutate(x=tree_col(tree_id), y=tree_row(tree_id))
    latest$state[is.na(latest$state)] = "S"

p <- ggplot(latest, aes(x*between_rows,
                  y*between_trees,
                  colour=state)) +
    geom_point(size=0.1) +
    theme(aspect.ratio=1) +
    guides(colour=FALSE) +
    scale_colour_manual(values=c("S"="green", "E"="blue", "I"="red", "R"="white")) +
    theme_void()

    jpeg(paste("bbsc-scout-", t, ".jpg", sep=""),
         width=width*between_rows, height=length*between_trees,
         quality=100, res=200)
    print(p)
    dev.off()
}
```
