## Supplementary material for "EMULSION: transparent and flexible multiscale stochastic models in human, animal and plant epidemiology": S1 File: bbsc.sh

```
#!/bin/bash
# Usage:
#   ./bbsc.sh nb_repetitions grove_width grove_length between_tree_space between_row_space [repetition_to_show]
# Example:
#   ./bbsc.sh 500 14 120 4 6 0
#
# - Run two scenarios for the simulation of BBSC spread and control:
#   1) no control
#   2) annual scouting and roguing
# - Transform log files to make it easier to parse in R scripts
#   log file contains lines such as:
#   simu_id,step,agent_class,agent_id,is_X=True
#   where X is E, I or R.
#   => replace 'is_X=True' with 'X'
# - Run R script

NBRUNS=$1
WIDTH=$2
LENGTH=$3
DY=$4
DX=$5
SIMUID=$6

if [ -z "$SIMUID" ]
then
    SIMUID=0
fi

python3 -m emulsion run bbsc.yaml -r $NBRUNS --silent --output-dir outputs/bbsc-nocontrol -p grove_width=$WIDTH -p grove_length=$LENGTH -p between_tree_space=$DY -p between_row_space=$DX -p p_detection=0
python3 -m emulsion run bbsc.yaml -r $NBRUNS --silent --output-dir outputs/bbsc-scout -p grove_width=$WIDTH -p grove_length=$LENGTH -p between_tree_space=$DY -p between_row_space=$DX

sed 's/is_\([A-Za-z]*\)=True/\1/g' outputs/bbsc-nocontrol/log.txt > bbsc-log-nocontrol.csv
sed 's/is_\([A-Za-z]*\)=True/\1/g' outputs/bbsc-scout/log.txt >bbsc-log-scout.csv

cp outputs/bbsc-nocontrol/counts.csv bbsc-counts-nocontrol.csv
cp outputs/bbsc-scout/counts.csv bbsc-counts-scout.csv

Rscript bbsc.R $WIDTH $LENGTH $DY $DX $SIMUID
```
