## Supplementary material for "EMULSION: transparent and flexible multiscale stochastic models in human, animal and plant epidemiology": S1 File: compare_paradigms.R

```
## setwd("/Users/picault/recherche/articles/2019/Software Paper/ploscb_software/figures/S1_appendix")

library(ggplot2)
library(tidyr)
library(dplyr)

args = commandArgs(trailingOnly=TRUE)
## Rscript compare_paradigms MODEL

model = args[1]

paradigms <- c("SimInf-compartment", "EMULSION-compartment", "EMULSION-hybrid", "EMULSION-IBM")

deter <- read.table(paste("determ_", model, ".csv", sep=""), sep=",", header=TRUE)
states <- factor(c("S", "I", "R"), levels=c("S", "I", "R"))

all <- data.frame()
for (para in paradigms) {
    tmp <- read.table(paste(para, "_", model, ".csv", sep=""), sep=",", header=TRUE) %>%
        select(simu_id, step, S, I, R, percentage_prevalence)
    tmp <- tmp %>% mutate(method=para)
    all <- rbind(all, tmp)
}
all$method = factor(all$method, levels=c("SimInf-compartment", "EMULSION-compartment", "EMULSION-hybrid", "EMULSION-IBM"))
NB_RUNS <- max(all$simu_id) + 1

mutall <- all %>%
    gather("state", "qty", c("S", "I", "R")) %>%
    mutate(state=factor(state, levels=c("S", "I", "R")))

p <- ggplot(mutall, aes(x=step, colour=state, y=qty)) +
    facet_wrap(~ method, ncol=4) +
    geom_line(aes(group=interaction(simu_id, state)), alpha=0.03) +
    geom_line(data=deter, aes(x=time, y=S, colour=states[1]), size=1, alpha=0.9) +
    geom_line(data=deter, aes(x=time, y=I, colour=states[2]), size=1, alpha=0.9) +
    geom_line(data=deter, aes(x=time, y=R, colour=states[3]), size=1, alpha=0.9) +
    ylab("population") +
    xlab("time") +
    ylim(0, 120) +
    guides(colour=guide_legend(title="Health state:")) +
    scale_colour_manual(values=c("S"="lightskyblue", "I"="red", "R"="midnightblue")) +
    theme_bw() +
    theme(legend.position = "bottom", legend.direction="horizontal")

pdf(paste("comparison_", model, ".pdf", sep=""),
    ## paper="a4r",
    width=12, height=4)
print(p)
dev.off()


p2 <- ggplot(all, aes(x=step, y=percentage_prevalence)) +
    facet_wrap(~method, ncol=4) +
    geom_line(aes(group=simu_id), alpha=0.03) +
    geom_line(data=deter, aes(x=time, y=100*I/(S+I+R)), alpha=0.9, size=1) +
    ylab("prevalence (%)") +
    xlab("time") +
    ylim(0, 75) +
    ## guides(colour=guide_legend(title="state")) +
    ## scale_colour_manual(values=c("S"="lightskyblue", "I"="red", "R"="midnightblue")) +
    theme_bw()

pdf(paste("prevalence_", model, ".pdf", sep=""),
    ## paper="a4r",
    width=12, height=3)
print(p2)
dev.off()


## agg <- all %>%
##     group_by(step, method) %>%
##     summarise(med=median(percentage_prevalence),
##               q5=quantile(percentage_prevalence, probs=0.05),
##               q95=quantile(percentage_prevalence, probs=0.95))

## p <- ggplot(agg, aes(x=step)) +
##     facet_wrap(~ method, ncol=4) +
##     geom_line(aes(y=med), size=1, alpha=0.6) +
##     geom_ribbon(aes(ymin=q5, ymax=q95), colour=FALSE, alpha=0.1) +
##     geom_line(data=deter, aes(x=time, y=100*I/(S+I+R)),
##               linetype="dashed", size=1, alpha=0.6) +
##     theme_bw() +
##     ylab("prevalence (%)") +
##     xlab("time")
## p
```
