## Supplementary material for "EMULSION: transparent and flexible multiscale stochastic models in human, animal and plant epidemiology": S1 File: compare_paradigms.sh

```
#!/bin/bash
# Usage:
#     ./compare_paradigms.sh MODEL NBRUNS
# Example:
#     ./compare_paradigms.sh SIR_demo 500

# - Run EMULSION for models of the form compartment_MODEL.yaml,
#   hybrid_MODEL.yaml and IBM_MODEL.yaml for NBRUNS stochastic
#   repetitions
# - Generate diagrams using EMULSION
# - Run the deterministic version of MODEL (determ_MODEL.py)
# - Plot a comparison of model outputs

MODEL="$1"
NBRUNS="$2"

if [ -z "$NBRUNS" ]
then
    NBRUNS=500
fi

echo "Running SimInf code for SIR and SIR with demo"
Rscript SimInf_code.R $NBRUNS

for paradigm in compartment hybrid IBM
do
    echo "Producing state machine diagrams for model $MODEL with $paradigm modelling paradigm"
    python -m emulsion diagrams "$paradigm""_$MODEL.yaml" --format pdf
    echo "Running model $MODEL with $paradigm modelling paradigm for $NBRUNS repetitions"
    python -m emulsion run "$paradigm""_$MODEL.yaml" -r "$NBRUNS" --silent
    mv outputs/counts.csv "EMULSION-$paradigm""_$MODEL.csv"
done

echo "Running deterministic version of model $MODEL"
python "determ_$MODEL.py" > "determ_$MODEL.csv"

echo "Producing a comparison of outputs for model $MODEL"
Rscript compare_paradigms.R "$MODEL"
echo "See files comparison_$MODEL.pdf and prevalence_$MODEL.pdf"
```
