## Supplementary material for "EMULSION: transparent and flexible multiscale stochastic models in human, animal and plant epidemiology": S1 File: determ_SIR_demo.py

```
from scipy.integrate import odeint
import numpy as np

b = 0.011
mu = 0.01
beta = 0.5
gamma = 0.1

def F(X, t):
    s, i, r = X
    return (
        -(mu + beta * i / (s + i + r)) * s + b * (s + i + r),
        beta * i * s / (s + i + r) - (gamma + mu) * i,
        gamma * i - mu * r
    )

temps = np.linspace(0, 100, 1000)
sol = odeint(F, [90, 10, 0], temps)

print("time,S,I,R")
for t, line in zip(temps, sol):
    s, i, r = line
    print("{},{},{},{}".format(t, s, i, r))
```
