## Supplementary material for "EMULSION: transparent and flexible multiscale stochastic models in human, animal and plant epidemiology": S1 File: measles.R

```
library(ggplot2)
library(tidyr)
library(dplyr)

theme_EMULSION <- theme_bw() +
    theme(text=element_text(family="Helvetica", size=20),
          axis.text=element_text(family="Helvetica", size=16),
          legend.text=element_text(family="Helvetica", size=18),
          )

t <- read.table('outputs/measles/counts.csv', header=TRUE, sep=",")

t <- t %>% group_by(simu_id) %>% mutate(incidence=c(0, diff(cum_incidence)))

## p <- ggplot(t, aes(x=step/365)) +
##     xlab("Time (year)") +
##     ylab("Daily incidence") +
##     ## geom_line(aes(y=I, group=simu_id, colour="infectious"), alpha=0.2) +
##     geom_line(aes(y=incidence, group=simu_id, colour="daily incidence"), alpha=0.05, size=0.1, show.legend=FALSE) +
##     geom_vline(xintercept=2000/365, colour="black", size=2) +
##     scale_colour_manual(values=c("daily incidence" = "blue", "infectious" = "red")) +
##     scale_x_continuous(breaks=seq(0,10,1), limits=c(0,NA)) +
##     theme_EMULSION #+
##     ## guides(colour=NULL) #colour=guide_legend(title="", override.aes=list(size=2, alpha=1))) +
##     ## theme(legend.position="bottom",
##     ##       legend.direction="horizontal")
## p

inc <- t %>%
    group_by(step) %>%
    summarise(med=median(incidence),
              q5=quantile(incidence, probs=0.05, na.rm=TRUE),
              q95=quantile(incidence, probs=0.95, na.rm=TRUE))

p <- ggplot(inc, aes(x=step/365,))+
    xlab("Time (year)") +
    ylab("Daily incidence") +
    geom_line(aes(y=med), colour="blue") +
    geom_ribbon(aes(ymin=q5,ymax=q95), fill="blue", colour=FALSE, alpha=0.2) +
    geom_vline(xintercept=2000/365, colour="black", size=2) +
    scale_x_continuous(breaks=seq(0,10,1), limits=c(0,NA)) +
    theme_EMULSION

jpeg("measles-outputs.jpg", width=1200, height=500, quality=100, res=150)
print(p)
dev.off()
```
