## Supplementary material for "EMULSION: transparent and flexible multiscale stochastic models in human, animal and plant epidemiology": S1 File: README.md

EMULSION example files
======================

This directory contains all files required to reproduce the examples
given in the Supporting Information for Picault *et al*, "EMULSION:
transparent and flexible multiscale stochastic models in
human, animal or plant epidemiology".

Part A - Files used in S1 Appendix
-------------------------------------------

- Files `compartment_SIR.yaml`, `hybrid_SIR.yaml` and `IBM_SIR.yaml`
  describe a classical SIR model, implemented using respectively a
  compartmental, hybrid and individual-based modelling paradigm.
- Files `compartment_SIR_demo.yaml`, `hybrid_SIR_demo.yaml` and
  `IBM_SIR_demo.yaml` describe a classical SIR model with births and
  deaths, implemented using respectively a compartmental, hybrid and
  individual-based modelling paradigm.
- The R script `SimInf_code.R` produces outputs for both models using
  the compartment-based paradigm implemented with the R library SimInf
  (Widgren *et al.* 2016).

After installing EMULSION, each model can be run and plotted
separately, for instance to plot the first one, with the state machine
diagram, after 50 stochastic repetitions:

```
emulsion run --plot compartment_SIR.yaml -r 50 --silent --view-model
```

State machine diagrams can be produced without running the simulation:

```
emulsion diagrams compartment_SIR.yaml
```


The comparison of the three modelling paradigms (+ SimInf outputs) can be done using the
shell script provided here (assuming R is installed, with packages `SimInf`,
`ggplot2`, `tidyr` and `dplyr`):

```
./compare_paradigms.sh SIR
./compare_paradigms.sh SIR_demo
```

This will produce the files `comparison_SIR.pdf`,
`prevalence_SIR.pdf`, `comparison_SIR_demo.pdf` and
`prevalence_SIR_demo.pdf`: the resulting figures (fig. 2 and 6 in S1
Appendix).


Part B - Files used in S1-S5 Text
---------------------------------

The following files demonstrate the step-by-step design of a complex
epidemiological model in EMULSION:

- File `hybrid_SIR.yaml` is a SIR model implemented using the hybrid
  modelling paradigm. It is the starting point of this example.
- In model `hybrid_SIR_ages.yaml`, demography has been introduced
  through two age groups, juveniles (J) and adults (A), together with
  births and deaths.
- Model `hybrid_SIR_ages_metapop.yaml`, is scaled up to the
  metapopulation level, with a data-driven contact network. A small
  code add-on is provided in `metapop_movements.py` to read the
  CSV data file (`data/moves.csv`)
- Model `hybrid_VSIR_ages_metapop.yaml` introduces a target control
  measure (the vaccination of juveniles in spring), based on an
  additional health state and a calendar. It is also based on the
  previous code add-on and data file.


Part C - Files used in S2 Appendix
-------------------------------------------

### Measles: multi-annual outbreaks and vaccination

- File `measles.yaml` is the re-implementation within EMULSION of a
  model of a human disease (measles), based on the original model
  presented by Keeling and Rohani (2008) in their book.
- File `measles.R` is a R script intended to display daily measles
  incidence with the median and 90% confidence interval from raw
  EMULSION outputs

State machine diagrams (fig. 2 in S2 Appendix) can be produced without running the simulation:

```
emulsion diagrams measles.yaml --format pdf
```

To reproduce the results presented in S2 Appendix, run the following
instructions (for test purposes, reduce the number of repetitions!):

```
emulsion run measles.yaml -r 500 --silent --output-dir outputs/measles
```

The presentation of the results can be done using the `measles.R`
script (assuming R is installed, with packages `ggplot2`, `tidyr` and
`dplyr`), which produces files `measles-outputs.pdf` (figure 3 in S2
Appendix):

```
Rscript measles.R
```

### Rift Valley fever: a vector-borne disease

- File `rvf.yaml` is the re-implementation within EMULSION of a model
  of Rift Valley fever, a vector-borne disease, based on the original
  model presented by Cavalerie *et al.* (2015).
- File `rvf.R` is a R script intended to reproduce four figures from
  original article based on raw EMULSION outputs

State machine diagrams (fig. 5 in S2 Appendix) can be produced without running the simulation:

```
emulsion diagrams rvf.yaml --format pdf
```

To reproduce the results presented in S2 Appendix, run the following instructions:

```
emulsion run rvf.yaml -r 1000 --silent --output-dir outputs/rvf
```

The presentation of the results can be done using the `rvf.R`
script (assuming R is installed, with packages `ggplot2`, `tidyr` and
`dplyr`), which produces files `rvf2.jpg`, `rvf4-1.jpg`, `rvf4-2.jpg`,
`rvf4-3.jpg` (figure 6 in S2 Appendix):

```
Rscript rvf.R
```


### Bahia Bark Scaling of Citrus: spread and control

- File `bbsc.yaml` is the re-implementation within EMULSION of a
  spatially explicit model of a plant disease (Bahia Bark Scaling of
  Citrus), based on the original model presented by Cunniffe *et al.*
  (2014).
- File `bbsc.R` is a R script intended to compare two scenarios (no
  control vs. annual scouting and roguing) and display the spatial
  spread on one repetition for each case.


State machine diagrams (fig. 8 in S2 Appendix) can be produced without running the simulation:

```
emulsion diagrams bbsc.yaml --format pdf
```

To reproduce the results presented in S2 Appendix, run the following script:

```
# nb_repetitions grove_width grove_length between_tree_space between_row_space repetition_to_show
./bbsc.sh 500 14 120 4 6 0
```

The presentation of the results is done by this script using the
`bbsc.R` script (assuming R is installed, with packages `ggplot2`,
`tidyr` and `dplyr`), which produces files `bbsc-asymptomatic.jpg`
(fig. 9 in S2 Appendix) and a series of images of the grove evolution
for each scenario: `bbsc-nocontrol-N.jpg` and `bbsc-scout-N.jpg` (N=0,
2, ..., 20) shown in fig. 10 in S2 Appendix.
