## Supplementary material for "EMULSION: transparent and flexible multiscale stochastic models in human, animal and plant epidemiology": S1 File: rvf.R

```
## This file reproduces figures from Cavalerie et al 2015
library(ggplot2)
library(dplyr)
library(tidyr)

theme_EMULSION <- theme_bw() +
    theme(text=element_text(family="Helvetica", size=20),
          axis.text=element_text(family="Helvetica", size=16),
          legend.text=element_text(family="Helvetica", size=18),
          )

t <- read.table('outputs/rvf/counts.csv', header=TRUE, sep=",")

REPET = max(t$simu_id) - min(t$simu_id) + 1


# Adult vector population / forcing
pop <- t %>% group_by(simu_id, step) %>%
    summarise(nbvect=EV + IV + SV)
forcing <- t %>% group_by(step) %>%
    summarise(phi=first(phi))

p <- ggplot(pop %>% filter(step > 365, step <= 365*3),
             aes(x=(step-365)/30, y=nbvect)) +
    geom_line(aes(group=factor(simu_id), colour="adult vectors"),
              alpha=min(1,50/REPET)) +
    geom_line(data=forcing, aes(y=phi*4e06, colour="emergence rate"), size=1) +
    scale_y_continuous(breaks=seq(0, 4e06, 1e06), limits=c(0, 4e06),
                       sec.axis = sec_axis(~ . / 4e06, name="Vector emergence rate")) +
    scale_x_continuous(breaks=seq(0, max(t$step)/30+1, 3), limits=c(0, 24)) +
    scale_colour_manual(values=c("adult vectors"="black", "emergence rate"="red")) +
    guides(colour = guide_legend(title="", keywidth=3,
                                 override.aes=list(alpha=1))) +
    xlab('Time (months)') + ylab('Adult vector population') +
    theme(legend.position="bottom",
          legend.direction="horizontal") +
    theme_EMULSION

jpeg("rvf2.jpg", width=1200, height=900, quality=100, res=150)
print(p)
dev.off()

# Persistence
pers <- t %>%
    filter(step > 365) %>%
    group_by(simu_id, step) %>%
    summarise(pers=as.integer(EH + IH + EV + IV + IA > 0)) %>%
    group_by(step) %>%
    summarise(pers=sum(pers))
pers$step <- pers$step - 365

p <- ggplot(pers, aes(x=step/365, y=pers/REPET)) +
    geom_line(size=1) +
    scale_y_continuous(breaks=seq(0, 1, .2), limits=c(0, 1)) +
    scale_x_continuous(breaks=seq(0, max(t$step)/365, 1), limits=c(0, 5.05)) +
    xlab('Time (years)') + ylab('Persistence') +
    theme_EMULSION

jpeg("rvf4-3.jpg", width=1200, height=900, quality=100, res=150)
print(p)
dev.off()


# Host states
hdat <- t %>%
    select(simu_id, EH, IH, RH, SH, step) %>%
    gather("state", "nb", 2:5)
vdat <- t %>%
    select(simu_id, EV, IV, IA, SV, SA, step)  %>%
    gather("state", "nb", 2:6)

hdat$state <- factor(hdat$state, levels=c("SH", "EH", "IH", "RH"))
vdat$state <- factor(vdat$state, levels=c("SV", "EV", "IV", "SA", "IA"))

p <- ggplot(dat=hdat %>% filter(step > 365), # %>% filter(simu_id < 5, step < 100),
            aes(x=(step-365)/365, y=nb, colour=state)) +
    geom_line(aes(colour=state, group=interaction(simu_id, state)),
              alpha=min(1, 50/REPET)) +
    scale_colour_manual(values=c('SH'='green', 'EH'='orange', 'IH'='red', 'RH'='blue')) +
    scale_x_continuous(breaks=seq(0, max(t$step)/365, 1), limits=c(0, 5.05)) +
    guides(colour=guide_legend(override.aes=list(alpha=1, size=3), keywidth=2),
           linetype=FALSE) +
    theme(legend.position="bottom", legend.direction="horizontal") +
    xlab('Time (years)') + ylab('Number of hosts') +
    theme_EMULSION

jpeg("rvf4-1.jpg", width=1200, height=900, quality=100, res=150)
print(p)
dev.off()

# Infected vectors
p <- ggplot(dat=vdat %>% filter(state=='IV', step > 365),
       aes(x=(step-365)/365, y=nb, colour=state)) +
    geom_line(aes(colour=state, group=factor(simu_id)), alpha=min(1, 50/REPET)) +
    scale_colour_manual(values=c('SV'='darkgreen', 'EV'='orange', 'IV'='red', 'RV'='blue')) +
    scale_x_continuous(breaks=seq(0, max(t$step)/365, 1), limits=c(0, 5.05)) +
    guides(colour=FALSE, linetype=FALSE) +
    theme_bw() +
    xlab('Time (years)') + ylab('Number of infected vectors') +
    theme_EMULSION

jpeg("rvf4-2.jpg", width=1200, height=900, quality=100, res=150)
print(p)
dev.off()
```
