## Supplementary material for "EMULSION: transparent and flexible multiscale stochastic models in human, animal and plant epidemiology": S1 File: SimInf_code.R

```
library(SimInf)
library(ggplot2)
library(tidyr)
library(dplyr)

args = commandArgs(trailingOnly=TRUE)
NBRUNS = as.integer(args[1])

model <- SIR(u0=data.frame(S=rep(90, NBRUNS), I=10, R=0),
             beta=0.5, gamma=0.1,
             tspan=0:100)
result <- run(model)

traj <- trajectory(model = result) %>%
    rename(simu_id=node, step=time) %>%
    mutate(percentage_prevalence=100*I/(S+I+R))
write.table(traj, file="SimInf-compartment_SIR.csv", sep=",", row.names=FALSE, col.names=TRUE)

model <- mparse(
    transitions = c("S -> beta*S*I/(S+I+R) -> I",
                    "S -> mu*S -> @",
                    "@ -> b*(S+I+R) -> S",
                    "I -> gamma*I -> R",
                    "I -> mu*I -> @",
                    "R -> mu*R -> @"),
    compartments = c("S", "I", "R"),
    gdata =  c(beta=0.5, gamma=0.1, b=0.011, mu=0.01),
    u0=data.frame(S=rep(90, NBRUNS), I=10, R=0),
    tspan=0:100
)

result <- run(model)

traj <- trajectory(model = result) %>%
    rename(simu_id=node, step=time) %>%
    mutate(percentage_prevalence=100.0*I/(S+I+R))

write.table(traj, file="SimInf-compartment_SIR_demo.csv", sep=",",
            row.names=FALSE, col.names=TRUE)
```
