## Supplementary material for "EMULSION: transparent and flexible multiscale stochastic models in human, animal and plant epidemiology": S2 File: functions.html

emulsion.tools.functions — EMULSION (Epidemiological Multi-Level Simulation framework)

### Source code for emulsion.tools.functions

```
""".. module:: emulsion.tools.functions

Additional functions for symbolic computing in YAML model definition files.

All functions in this module can be used in EMULSION models.
"""

# [HEADER]


import numpy as np

[docs]def IfThenElse(condition, val_if_true, val_if_false):
    """Ternary conditional function: return either *val_if_true* or
    *val_if_false* depending on *condition*.

    Example
    -------
    Here in a parameter definition (assuming that ``summer_period`` is
    defined e.g. in the *calendars* section of the model)::

      average_temperature:
        desc: 'average temperature for the season'
        value: 'IfThenElse(summer_period, 25, 8)'

    Parameters
    ----------
    condition: bool
        a boolean expression
    val_if_true: number
        value to return if the expression is True
    val_if_false: number
        value to return if the expression is False

    Returns
    -------
    number:
        One of the two values, depending on *condition*.

    """
    return val_if_true if condition else val_if_false

IfThenElse.__USER_FUNCTION__ = ['Functions Available for Models']


# BEWARE: using lambdify, by default And and Or fall back to numpy's
# *binary* operators, so that more than 3 conditions linked by And or
# Or trigger a TypeError ('return arrays must be of ArrayType')
# see topic addressed there:
# https://stackoverflow.com/questions/42045906/typeerror-return-arrays-must-be-of-arraytype-using-lambdify-of-sympy-in-python

# To avoid problems, please use AND / OR (fully in UPPERCASE) instead
# of And / Or (Capitalized) in the conditions of Emulsion models

[docs]def AND(*values):
    """Return a logical AND (conjunction) between all the values.

    Example
    -------
    To define symptomatic individuals as infected for at least 5 days::

      is_symptomatic:
        desc: 'test if individuals has symptoms'
        value: 'AND(is_I, duration_in_health_state >= 5)'

    Parameters
    ----------
    *values: list
        boolean values separated by commas

    Returns
    -------
    bool:
        True if **all** values are True, False otherwise.
    """
    return all(values)

AND.__USER_FUNCTION__ = ['Functions Available for Models']

[docs]def OR(*values):
    """Return a logical OR (disjunction) between all the values.

    Example
    -------
    To sell animals depending on either their weight or their age::

      transitions:
        ...
        - from: Fattening
          to: Sold
          proba: 1
          cond: 'OR(weight >= weight_thr, age >= age_thr)'

    Parameters
    ----------
    *values: list
        boolean values separated by commas

    Returns
    -------
    bool:
        True if **at least one** of the values is True, False otherwise.
    """
    return any(values)

OR.__USER_FUNCTION__ = ['Functions Available for Models']

[docs]def random_bool(proba_success: float) -> int:
    """Return a random boolean value (actually, 0 or 1) depending on
    *proba_success*.

    Example
    -------
    Set the value of a state variable ``has_symptoms`` when entering
    in the infectious state::

      states:
        ...
        - I:
          ...
          on_enter:
            - set_var: has_symptoms
              value: 'random_bool(proba_symptomatic)'

    Parameters
    ----------
    proba_success: float in [0,1]
        probability of returning 1 (True)

    Returns
    -------
    int:
        either 1 with probability *proba_success*, or 0 with
        probability 1-*proba_success*

    """
    return np.random.binomial(1, proba_success)

random_bool.__USER_FUNCTION__ = ['Functions Available for Models']

[docs]def random_choice(*values):
    """Return a value chosen randomly among those provided (equiprobable
    sampling).

    Example
    -------
    To init ``age`` among three typical values::

      prototypes:
        individuals:
          init_individual:
            age: 'random_choice(10, 50, 200)'

    Parameters
    ----------
    *values: list
        the possible values, separated by commas

    Returns
    -------
    val:
        one of the values (equiprobable choice)

    """
    return np.random.choice(values)

random_choice.__USER_FUNCTION__ = ['Functions Available for Models']

[docs]def random_choice_weighted(*values):
    """Return a value chosen randomly among those provided (but not
    equiprobably).

    Example
    -------
    To init ``age`` among three typical values which respectively
    represent 10%, 20% and 70% of the population::

      prototypes:
        individuals:
          init_individual:
            age: 'random_choice_weighted(10, 0.1, 50, 0.2, 200, 0.7)'

    Parameters
    ----------
    values: list
        possibles choices and their weight, alternatively, separated
        by commas. Weights are normalized to be used as probabilities.

    Returns
    -------
    One of the choices

    """
    choices_and_values = zip(*[iter(values)]*2)
    choices, weights = zip(*choices_and_values)
    total = sum(weights)
    probas = [w / total for w in weights]
    return np.random.choice(choices, p=probas)

random_choice_weighted.__USER_FUNCTION__ = ['Functions Available for Models']

[docs]def random_multinomial(number_of_samples, *probas):
    """Return a multinomial sample based on the specified probabilities.

    Parameters
    ----------
    number_of_samples: int
        number of experiments
    probas: list
        list of probabilities

    Returns
    -------
    list:
        the drawn samples
    """
    return list(np.random.multinomial(number_of_samples, probas))

random_multinomial.__USER_FUNCTION__ = ['Functions Available for Models']


#   _____ _                _             _         _
#  / ____| |              | |           | |       | |
# | (___ | |__   ___  _ __| |_ ___ _   _| |_ ___  | |_ ___
#  \___ \| '_ \ / _ \| '__| __/ __| | | | __/ __| | __/ _ \
#  ____) | | | | (_) | |  | || (__| |_| | |_\__ \ | || (_) |
# |_____/|_| |_|\___/|_|   \__\___|\__,_|\__|___/  \__\___/
#                                                          _
#                                                         | |
#  _ __  _   _ _ __ ___  _ __  _   _   _ __ __ _ _ __   __| | ___  _ __ ___
# | '_ \| | | | '_ ` _ \| '_ \| | | | | '__/ _` | '_ \ / _` |/ _ \| '_ ` _ \
# | | | | |_| | | | | | | |_) | |_| |_| | | (_| | | | | (_| | (_) | | | | | |
# |_| |_|\__,_|_| |_| |_| .__/ \__, (_)_|  \__,_|_| |_|\__,_|\___/|_| |_| |_|
#                       | |     __/ |
#                       |_|    |___/

[docs]def random_uniform(low: float, high: float) -> float:
    """Return a random value drawn from a uniform distribution between
    *low* (inclusive) and *high* (exclusive).

    Example
    -------
    In a prototype definition::

      age: random_uniform(min_age, max_age)

    Parameters
    ----------
    low: float
        lower boundary of the sample interval (inclusive)
    high: float
        upper boundary of the sample interval (exclusive)

    Returns
    -------
    float:
        a random value sampled according to a uniform distribution
        between *low* and *high*

    See also
    --------
    numpy.random.uniform

    """
    return np.random.uniform(low, high)

random_uniform.__USER_FUNCTION__ = ['Functions Available for Models']

[docs]def random_integers(low: int, high: int) -> int:
    """Return a random integer value drawn from a discrete uniform
    distribution between *low* and *high* (both inclusive).

    Example
    -------
    In a prototype definition::

      age: random_integers(min_age, max_age)

    Parameters
    ----------
    low: int
        lower boundary of the sample interval (inclusive)
    high: int
        upper boundary of the sample interval (inclusive)

    Returns
    -------
    int:
        a random integer value sampled according to a discrete uniform
        distribution between *low* and *high*

    See also
    --------
    numpy.random.random_integers
    """
    return np.random.random_integers(low, high)

random_integers.__USER_FUNCTION__ = ['Functions Available for Models']

[docs]def random_exponential(scale: float) -> float:
    """Return a random value drawn from an exponential distribution of
    rate 1/*scale* (thus of mean *scale*).

    Example
    -------
    In a prototype definition::

      time_to_live: random_exponential(mean_duration)

    Parameters
    ----------
    scale: float
        the scale parameter of the distribution, i.e. the inverse of the rate

    Returns
    -------
    float:
        a random value sampled according to an exponential
        distribution of rate 1/*scale*

    See also
    --------
    numpy.random.exponential

    """
    return np.random.exponential(scale)

random_exponential.__USER_FUNCTION__ = ['Functions Available for Models']

[docs]def random_beta(a: float, b: float) -> float:
    """Return a random value drawn from a beta distribution of parameters
    *a* and *b*.

    Example
    -------
    In a prototype definition::

      age: 'random_beta(2, 5) * age_max'

    Parameters
    ----------
    a: float, positive (>0)
        first shape parameter of the beta distribution
    b: float, positive (>0)
        second shape parameter of the beta distribution

    Returns
    -------
    float:
        a random value sampled according to a beta distribution of
        parameters *a* and *b*

    See also
    --------
    numpy.random.beta
    """
    return np.random.beta(a, b)

random_beta.__USER_FUNCTION__ = ['Functions Available for Models']

[docs]def random_gamma(shape: float, scale: float) -> float:
    """Return a random value drawn from a gamma distribution of parameters
    *shape* and *scale*.

    Example
    -------
    In a prototype definition::

      age: 'random_gamma(3, 2)'

    Parameters
    ----------
    shape: float, positive (>0)
        the shape of the gamma distribution
    scale: float, positive (>0)
        the scale of the gamma distribution

    Returns
    -------
    float:
        a random value sampled according to a gamma distribution of
        parameters *shape* and *scale*

    See also
    --------
    numpy.random.gamma
    """
    return np.random.gamma(shape, scale)

random_gamma.__USER_FUNCTION__ = ['Functions Available for Models']

[docs]def random_normal(mn: float, sd: float) -> float:
    """Return a random value drawn from a normal distribution of mean *mn*
    and standard deviation *sd*.

    Example
    -------
    In a prototype definition::

      age: 'random_normal(100, 5)'

    Parameters
    ----------
    mn: float
        the mean of the normal distribution
    sd: float, positive (>=0)
        the standard deviation of the normal distribution

    Returns
    -------
    float:
        a random value sampled according to a normal distribution of
        mean *mn* and standard deviation *sd*

    See also
    --------
    numpy.random.normal

    """
    return np.random.normal(mn, sd)

random_normal.__USER_FUNCTION__ = ['Functions Available for Models']

[docs]def random_poisson(lam: float) -> int:
    """Return an integer random value drawn from a Poisson distribution of
    mean *lam*.

    Example
    -------
    In an action, e.g. here when computing how many newborn
    individuals will be produced::

      - from: Gestating
        to: NonGestating
        on_cross:
          - produce_offspring: newborn
            amount: 'random_poisson(1.05)'

    Parameters
    ----------
    lam: float
        the mean of the Poisson distribution

    Returns
    -------
    int:
        a random integer value sampled according to a Poisson distribution of
        mean *lam*

    See also
    --------
    numpy.random.poisson

    """
    return np.random.poisson(lam)

random_poisson.__USER_FUNCTION__ = ['Functions Available for Models']
```

### EMULSION

Epidemiological Multi-Level Simulation Framework

##### Navigation

- 1. Installation
- 2. Getting started with EMULSION
- 3. Modelling principles
- 4. Modelling language (basics)
- 5. Modelling language (advanced)
- 6. Feature examples
- 7. Information
- 8. License
- 9. High-level functions for model designers
- 10. emulsion package

##### Related Topics

- Documentation overview
  - Module code

##### Quick search

©2016, INRA and Univ. Lille.
|
Powered by Sphinx 1.8.5
& Alabaster 0.7.10
