## Supplementary material for "EMULSION: transparent and flexible multiscale stochastic models in human, animal and plant epidemiology": S2 File: simulation.html

emulsion.tools.simulation — EMULSION (Epidemiological Multi-Level Simulation framework)

### Source code for emulsion.tools.simulation

```
""".. module:: emulsion.tools.simulation

Tools for providing generic simulation classes.

"""

#[HEADER]


import abc
from   pathlib        import Path
from   os.path        import exists
from   typing         import List, Iterable, Tuple, Union

import numpy          as     np
import pandas         as     pd

from   tqdm           import trange, tqdm   # a nice replacement for progressbar2

from   sqlalchemy     import create_engine

from   emulsion.model import EmulsionModel

#   ____        _               _   __  __
#  / __ \      | |             | | |  \/  |
# | |  | |_   _| |_ _ __  _   _| |_| \  / | __ _ _ __   __ _  __ _  ___ _ __
# | |  | | | | | __| '_ \| | | | __| |\/| |/ _` | '_ \ / _` |/ _` |/ _ \ '__|
# | |__| | |_| | |_| |_) | |_| | |_| |  | | (_| | | | | (_| | (_| |  __/ |
#  \____/ \__,_|\__| .__/ \__,_|\__|_|  |_|\__,_|_| |_|\__,_|\__, |\___|_|
#                  | |                                        __/ |
#                  |_|                                       |___/

[docs]class OutputManager(object):
    """Manager to handle different outputs (csv, database,... etc)

    """

[docs]    def __init__(self, model=None, output_dir='', output_file='counts.csv', log_file='log.txt'):
        """Initialize the output manager, specifying the model and the
        directory where outputs will be stored.

        """
        self.model = model
        self.output_dir = output_dir
        self.output_file = output_file
        self.log_file = log_file

        # database engine
        self.engine = None
        self.counts_path = None
        self.log_path = str(Path(self.output_dir, self.log_file))
        # We choose csv file by default
        self.output_type = 'csv'
        self.update_output_type()
        self.update_output_information()


[docs]    def update_output_type(self):
        """Update output type if specified in model

        """
        if 'type' in self.model.outputs:
            self.output_type = self.model.outputs['type']


[docs]    def update_output_information(self):
        """Update csv file path or database connection engine

        """
        if self.output_type == 'database':
            database_information = self.model.outputs['database_information']

            host = database_information['server_name']
            port = database_information['port']
            host = '{}:{}'.format(host, port) if port else host

            # dialect+driver://username:password@servername:port/database
            connection = '{}+{}://{}:{}@{}/{}'.format(
                database_information['dialect'],
                database_information['driver'],
                database_information['username'],
                database_information['password'],
                host,
                database_information['database'])
            self.engine = create_engine(connection)
            # self.engine.raw_connection()
            self.output_dir = self.output_dir.replace('/', '_')

        elif self.output_type == 'csv':
            self.counts_path = str(Path(self.output_dir, self.output_file))
        else:
            print('unknown output type !!!{]!!!')


[docs]    def update_outputs(self, df=None):
        """Update outputs: writing in csv file or in database

        """
        if self.output_type == 'csv':
            header = not exists(self.counts_path)
            with open(self.counts_path, 'a') as f:
                df.to_csv(f, header=header, index=False)
        elif self.output_type == 'database':
            df.to_sql(self.output_dir, self.engine, if_exists='append',
                      index=False, chunksize=10000)

#           _         _                  _    _____ _                 _
#     /\   | |       | |                | |  / ____(_)               | |
#    /  \  | |__  ___| |_ _ __ __ _  ___| |_| (___  _ _ __ ___  _   _| | __ _
#   / /\ \ | '_ \/ __| __| '__/ _` |/ __| __|\___ \| | '_ ` _ \| | | | |/ _` |
#  / ____ \| |_) \__ \ |_| | | (_| | (__| |_ ____) | | | | | | | |_| | | (_| |
# /_/    \_\_.__/|___/\__|_|  \__,_|\___|\__|_____/|_|_| |_| |_|\__,_|_|\__,_|

[docs]class AbstractSimulation(object):
    """Abstract class from which any simulation class inherits.

    """

[docs]    def __init__(self, start_id: int = 0, model=None, model_path: str = '',
                 stock_agent: bool = True, output_dir: str = 'outputs/',
                 target_agent_class=None, save_results: bool = True,
                 input_dir: str=None,
                 load_from_file: str = None, save_to_file: str = None, **_):
        """Initialize the simulation.

        Args:
            start_id: ID of the (first) simulation
            model: instance of the model to run
            model_path: path to the filename holding the description of the
              model, used if *model* is None
            stock_agent: TODO
            output_dir: name of the directory for simulation outputs
            target_agent_class: agent class representing the top level in the
              simulation
            save_results: True if simulation outputs have to be saved,
              False otherwise. TODO: should be removed, and replaced
              by a set of OutputManagers dedicated to the specific
              expected outputs.
            load_from_file: a filename from which the initial state of
              the simulation (agents corresponding to levels with
              their state) is read (instead of running the
              `initialize_level` method)
            save_to_file: a filename in which the final state of the
              simulation (agents corresponding to levels with their
              state) is written (after running the `finalize_level`
              method)

        """
        # ID of simulation
        self.start_id = start_id
        self.model = EmulsionModel(filename=model_path) if model is None else model
        self.model.input_dir = input_dir
        self.stock_agent = stock_agent
        self.target_agent_class = target_agent_class
        self.output_manager = OutputManager(model=self.model, output_dir=output_dir)
        self.save_results = save_results
        self.save_to_file = save_to_file
        self.load_from_file = load_from_file

# filename = 'counts.csv'
        # self.counts_path = str(Path(output_dir, filename))

[docs]    @abc.abstractmethod
    def evolve(self, steps: int = 1):
        """Operations to perform at each time step. Should be defined in
        subclasses.

        Args:
            steps:

        """
        pass


[docs]    @abc.abstractmethod
    def run(self):
        """Entry point to simulation execution. Should be defined in
        subclasses.

        """
        pass


[docs]    def update_csv_counts(self, df, dparams: dict = {}):
        """Update the CSV recording of populations in each state. """
        if self.save_results:
            for name, value in dparams.items():
                df.insert(0, name, value)
            # with open(self.counts_path, 'a') as f:
            #     df.to_csv(f, header=header, index=False)
            self.output_manager.update_outputs(df=df)

# def counts_to_csv(self):
    #     """Record the counts into a CSV file."""
    #     self.counts.to_csv(self.counts_path, index=False)

#   _____ _                 _       _   _
#  / ____(_)               | |     | | (_)
# | (___  _ _ __ ___  _   _| | __ _| |_ _  ___  _ __
#  \___ \| | '_ ` _ \| | | | |/ _` | __| |/ _ \| '_ \
#  ____) | | | | | | | |_| | | (_| | |_| | (_) | | | |
# |_____/|_|_| |_| |_|\__,_|_|\__,_|\__|_|\___/|_| |_|

[docs]class Simulation(AbstractSimulation):
    """Simulation class is aimed at running one repetition of a given
    model (for several repetitions, use `MultiSimulation`).

    """

[docs]    def __init__(self, steps: int = 100, simu_id: int = 0,
                 silent: bool = False, quiet: bool = False, **others):
        """Create an instance of simulation.

        Args:
            steps: number of time steps to run
            simu_id: ID of the simulation
            silent: if False, show a progress bar during simulation execution
            quiet: if True, show no progressbar at all

        See Also:
            `emulsion.tools.simulation.AbstractSimulation`_
        """
        super().__init__(**others)
        self.simu_id = simu_id
        self.steps = steps
        self.silent = silent
        self.quiet = quiet

        del(others['target_agent_class'])
        others['simu_id'] = self.simu_id
        others['simulation'] = self
        self.agent = self.init_agent(**others) ##self.target_agent_class(**others)

        self.outputs_period = self.model.outputs[self.agent.level]['period']\
                                if self.agent.level in self.model.outputs else 1


[docs]    def log_path(self):
        """Return the log path used by current simulation"""
        return self.output_manager.log_path


[docs]    def init_agent(self, **others):
        """Create an agent from the target class."""
        toplevel_agent = self.target_agent_class(**others)
        # if self.load_from_file:
        #     toplevel_agent.load_state_from_file(self.simu_id, self.load_from_file)
        # else:
        #     toplevel_agent.initialize_level()
        # # print(toplevel_agent.statevars)
        return toplevel_agent


[docs]    def evolve(self, steps: int = 1):
        """Make the target agent evolve.

        Args:
            steps: the number of time steps to run
        """
        for _ in range(steps):
            self.agent.evolve()


[docs]    def run(self, dparams: dict = {}):
        """Make the simulation progress."""
        if self.silent or self.quiet:
            progress = range(self.steps)
        else:
            progress = trange(self.steps, desc=f'[Run {self.simu_id}]',
                              bar_format='{l_bar}{bar}| {n_fmt}/{total_fmt}')
        self.update_csv_counts(self.counts, dparams=dparams)
        for step in progress:
            self.agent.evolve()
            if step % self.outputs_period == 0:
                self.update_csv_counts(self.counts,
                                       # header=not exists(self.counts_path),
                                       dparams=dparams)

        ## make main level agent run a "finalization" procedure if any
        self.agent.finalize_level(simulation=self)
        if self.save_to_file:
            self.agent.save_state_to_file(self.simu_id, self.save_to_file)

@property
    def counts(self):
        """Return a pandas DataFrame contains counts of each process if existing.
        TODO: column steps need to be with one of process
        and NO column steps for inter herd

        """
        res = self.agent.counts
        # insert explcitily modified parameters if asked for log
        to_log = self.model.params_to_log
        for name in sorted(to_log.keys()):
            res.insert(0, name, to_log[name])
        # insert ID for the simulation
        res.insert(0, 'simu_id', self.simu_id)
        return res

# res = None
        # for comp in self.agent:
        #     try:
        #         counts = pd.DataFrame(comp.counts)
        #         res = res.join(counts, lsuffix='res', rsuffix='counts')\
        #                                         if not res is None else counts
        #     except AttributeError:
        #         pass
        #     except Exception as e:
        #         raise e
        # if not res is None:
        #     res.insert(0, 'steps', res.index)
        #     res.insert(0, 'simu_id', self.simu_id)
        # return res

#  __  __       _ _   _  _____ _                 _       _   _
# |  \/  |     | | | (_)/ ____(_)               | |     | | (_)
# | \  / |_   _| | |_ _| (___  _ _ __ ___  _   _| | __ _| |_ _  ___  _ __
# | |\/| | | | | | __| |\___ \| | '_ ` _ \| | | | |/ _` | __| |/ _ \| '_ \
# | |  | | |_| | | |_| |____) | | | | | | | |_| | | (_| | |_| | (_) | | | |
# |_|  |_|\__,_|_|\__|_|_____/|_|_| |_| |_|\__,_|_|\__,_|\__|_|\___/|_| |_|

[docs]class MultiSimulation(AbstractSimulation):
    """MultiSimulation can handle multiple repetitions of a given model.
    For sensibility study (same model with different values of variables),
    please check out SensitivitySimulation.

    """

[docs]    def __init__(self, multi_id: int = 0, nb_simu: int = 100,
                 set_seed: bool = False, silent: bool = False,
                 quiet: bool = False, dparams: dict = {}, **others):
        """Initialize a simulation with multiple repetitions of the same
        model.
        """
        super().__init__(**others)
        self.multi_id = multi_id
        self.nb_simu = nb_simu
        self.set_seed = set_seed
        self.others = others
        self.silent = silent
        self.quiet = quiet
        self.dparams = dparams

        self.d_simu = dict()

def __iter__(self):
        return self.d_simu.values().__iter__()

[docs]    def evolve(self, steps=1):
        for simu in self.d_simu.values():
            simu.evolve(steps=steps)


[docs]    def run(self, update=True):
        """Run all repetitions one by one.

        """
        if self.silent and not self.quiet:
            progress = tqdm(range(self.start_id, self.start_id+self.nb_simu),
                            bar_format='{l_bar}{bar}| {n_fmt}/{total_fmt}')
        else:
            progress = range(self.start_id, self.start_id+self.nb_simu)
        for simu_id in progress:
            if self.set_seed:
                np.random.seed(simu_id)
            if 'model' not in self.others:
                self.others['model'] = self.model
            simu = Simulation(simu_id=simu_id, silent=self.silent,
                              quiet=self.quiet, dparams=self.dparams,
                              **self.others)

            simu.run()

            if self.stock_agent:
                self.d_simu[simu_id] = simu

# if update:
            #     header = not exists(self.counts_path)
            #     self.update_csv_counts(simu.counts, header=header, dparams=dparams)

    @property
    def counts(self):
        l_counts = [simu.counts for simu in self]
        return pd.concat(l_counts).reset_index(drop=True)

[docs]    def write_dot(self):
        self.model.write_dot(self.others['output_dir'])

#   _____                _ _   _       _ _          _____ _                 _
#  / ____|              (_) | (_)     (_) |        / ____(_)               | |
# | (___   ___ _ __  ___ _| |_ ___   ___| |_ _   _| (___  _ _ __ ___  _   _| |
#  \___ \ / _ \ '_ \/ __| | __| \ \ / / | __| | | |\___ \| | '_ ` _ \| | | | |
#  ____) |  __/ | | \__ \ | |_| |\ V /| | |_| |_| |____) | | | | | | | |_| | |
# |_____/ \___|_| |_|___/_|\__|_| \_/ |_|\__|\__, |_____/|_|_| |_| |_|\__,_|_|
#                                             __/ |
#                                            |___/
#        _   _
#       | | (_)
#   __ _| |_ _  ___  _ __
#  / _` | __| |/ _ \| '_ \
# | (_| | |_| | (_) | | | |
#  \__,_|\__|_|\___/|_| |_|

[docs]class SensitivitySimulation(AbstractSimulation):
    """SensitivitySimulation can handle sensibility study with a given
    pandas DataFrame of parameters or a path linked with file which contains
    scenarios of parameters. Then it will be transformed to a dictionary of
    scenario in the ```d_scenario``` attribute.

    For instance, d_scenario could be the form (QFever model example) :
        {0: {'m': 0.7, 'q': 0.02 ...},
         1: {'m': 0.5, 'q': 0.02 ...},
         2: ...,
         ... }
    """

[docs]    def __init__(self, scenario_path=None, df=None, nb_multi=None, **others):
        super().__init__(**others)
        self.others = others
        self.others['start_id'] = 0

        # Retrieving DataFrame and creation of dictionnary of simulation/scenario
        df = pd.read_csv(scenario_path) if df is None else df
        self.nb_multi = len(df) if nb_multi is None else nb_multi
        self.d_scenario = df.to_dict(orient='index')
        self.d_multi = dict()


[docs]    def run(self):
        """Make the simulation advance."""
        bar_sens = tqdm(range(self.start_id, self.start_id+self.nb_multi),
                        desc='[Sensitivity Simulation]')

        # for multi_id, scenario in bar_sens(self.d_scenario.items()):
        for multi_id in bar_sens:
            scenario = self.d_scenario[multi_id]
            # Copy a model
            model = self.model.copy()
            # Modify model
            for name, value in scenario.items():
                try:
                    model.set_value(name, value)
                except Exception:
                    self.others[name] = value

            # Instantiate MultiSimulation and execution
            multi = MultiSimulation(multi_id=multi_id, model=model, **self.others)
            multi.run(dparams=scenario)

            if self.stock_agent:
                self.d_multi[multi_id] = multi

# if update:
            #     header = not exists(self.counts_path)

            #     counts = multi.counts
            #     for name, value in self.d_scenario[multi_id].items():
            #         counts.insert(0, name, value)

            #     self.update_csv_counts(counts, header=header)

    @property
    def counts(self):
        l_counts = []
        for multi_id, multi in self.d_multi.items():
            counts = multi.counts
            for name, value in self.d_scenario[multi_id].items():
                counts.insert(0, name, value)

            counts.insert(0, 'scenario_id', multi_id)
            l_counts.append(counts)
        print(counts)
        return pd.concat(l_counts).reset_index(drop=True)

[docs]    def write_dot(self):
        self.model.write_dot(self.others['output_dir'])
```

### EMULSION

Epidemiological Multi-Level Simulation Framework

##### Navigation

- 1. Installation
- 2. Getting started with EMULSION
- 3. Modelling principles
- 4. Modelling language (basics)
- 5. Modelling language (advanced)
- 6. Feature examples
- 7. Information
- 8. License
- 9. High-level functions for model designers
- 10. emulsion package

##### Related Topics

- Documentation overview
  - Module code

##### Quick search

©2016, INRA and Univ. Lille.
|
Powered by Sphinx 1.8.5
& Alabaster 0.7.10
