## Supplementary material for "EMULSION: transparent and flexible multiscale stochastic models in human, animal and plant epidemiology": S2 File: index.html

Overview: module code — EMULSION (Epidemiological Multi-Level Simulation framework)

### All modules for which code is available

- emulsion.agent.action
- emulsion.agent.atoms
- emulsion.agent.comparts
- emulsion.agent.core.abstract\_agent
- emulsion.agent.core.emulsion\_agent
- emulsion.agent.core.groups
- emulsion.agent.exceptions
- emulsion.agent.managers.abstract\_process\_manager
- emulsion.agent.managers.compart\_process\_manager
- emulsion.agent.managers.functions
- emulsion.agent.managers.group\_manager
- emulsion.agent.managers.ibm\_process\_manager
- emulsion.agent.managers.metapop\_process\_manager
- emulsion.agent.managers.multi\_process\_manager
- emulsion.agent.meta
- emulsion.agent.process
- emulsion.agent.views
- emulsion.init\_emulsion
- emulsion.model.emulsion\_model
- emulsion.model.exceptions
- emulsion.model.functions
- emulsion.model.state\_machines
- emulsion.tools.calendar
- emulsion.tools.functions
- emulsion.tools.graph
- emulsion.tools.misc
- emulsion.tools.parallel
- emulsion.tools.plot
- emulsion.tools.simulation
- emulsion.tools.state
- emulsion.tools.timing
- emulsion.tools.view
- sortedcontainers.sortedset

### EMULSION

Epidemiological Multi-Level Simulation Framework

##### Navigation

- 1. Installation
- 2. Getting started with EMULSION
- 3. Modelling principles
- 4. Modelling language (basics)
- 5. Modelling language (advanced)
- 6. Feature examples
- 7. Information
- 8. License
- 9. High-level functions for model designers
- 10. emulsion package

##### Related Topics

- Documentation overview

##### Quick search

©2016, INRA and Univ. Lille.
|
Powered by Sphinx 1.8.5
& Alabaster 0.7.10
