## Supplementary material for "EMULSION: transparent and flexible multiscale stochastic models in human, animal and plant epidemiology": S2 File: sortedset.html

sortedcontainers.sortedset — EMULSION (Epidemiological Multi-Level Simulation framework)

### Source code for sortedcontainers.sortedset

```
"""Sorted set implementation.

"""

from collections import Set, MutableSet, Sequence
from itertools import chain
import operator as op

from .sortedlist import SortedList, recursive_repr, SortedListWithKey

class SortedSet(MutableSet, Sequence):
    """
    A `SortedSet` provides the same methods as a `set`.  Additionally, a
    `SortedSet` maintains its items in sorted order, allowing the `SortedSet` to
    be indexed.

    Unlike a `set`, a `SortedSet` requires items be hashable and comparable.
    """
    def __init__(self, iterable=None, key=None, load=1000, _set=None):
        """
        A `SortedSet` provides the same methods as a `set`.  Additionally, a
        `SortedSet` maintains its items in sorted order, allowing the
        `SortedSet` to be indexed.

        An optional *iterable* provides an initial series of items to populate
        the `SortedSet`.

        An optional *key* argument defines a callable that, like the `key`
        argument to Python's `sorted` function, extracts a comparison key from
        each set item. If no function is specified, the default compares the
        set items directly.

        An optional *load* specifies the load-factor of the set. The default
        load factor of '1000' works well for sets from tens to tens of millions
        of elements.  Good practice is to use a value that is the cube root of
        the set size.  With billions of elements, the best load factor depends
        on your usage.  It's best to leave the load factor at the default until
        you start benchmarking.
        """
        # pylint: disable=redefined-variable-type
        self._key = key
        self._load = load

        self._set = set() if _set is None else _set

        _set = self._set
        self.isdisjoint = _set.isdisjoint
        self.issubset = _set.issubset
        self.issuperset = _set.issuperset

        if key is None:
            self._list = SortedList(self._set, load=load)
        else:
            self._list = SortedListWithKey(self._set, key=key, load=load)

        _list = self._list
        self.bisect_left = _list.bisect_left
        self.bisect = _list.bisect
        self.bisect_right = _list.bisect_right
        self.index = _list.index
        self.irange = _list.irange
        self.islice = _list.islice

        if key is not None:
            self.bisect_key_left = _list.bisect_key_left
            self.bisect_key_right = _list.bisect_key_right
            self.bisect_key = _list.bisect_key
            self.irange_key = _list.irange_key

        if iterable is not None:
            self._update(iterable)

    def __contains__(self, value):
        """Return True if and only if *value* is an element in the set."""
        return value in self._set

    def __getitem__(self, index):
        """
        Return the element at position *index*.

        Supports slice notation and negative indexes.
        """
        return self._list[index]

    def __delitem__(self, index):
        """
        Remove the element at position *index*.

        Supports slice notation and negative indexes.
        """
        _set = self._set
        _list = self._list
        if isinstance(index, slice):
            values = _list[index]
            _set.difference_update(values)
        else:
            value = _list[index]
            _set.remove(value)
        del _list[index]

    def _make_cmp(self, set_op, doc):
        "Make comparator method."
        def comparer(self, that):
            "Compare method for sorted set and set-like object."
            # pylint: disable=protected-access
            if isinstance(that, SortedSet):
                return set_op(self._set, that._set)
            elif isinstance(that, Set):
                return set_op(self._set, that)
            else:
                return NotImplemented

        comparer.__name__ = '__{0}__'.format(set_op.__name__)
        doc_str = 'Return True if and only if Set is {0} `that`.'
        comparer.__doc__ = doc_str.format(doc)

        return comparer

    __eq__ = _make_cmp(None, op.eq, 'equal to')
    __ne__ = _make_cmp(None, op.ne, 'not equal to')
    __lt__ = _make_cmp(None, op.lt, 'a proper subset of')
    __gt__ = _make_cmp(None, op.gt, 'a proper superset of')
    __le__ = _make_cmp(None, op.le, 'a subset of')
    __ge__ = _make_cmp(None, op.ge, 'a superset of')

    def __len__(self):
        """Return the number of elements in the set."""
        return len(self._set)

    def __iter__(self):
        """
        Return an iterator over the Set. Elements are iterated in their sorted
        order.

        Iterating the Set while adding or deleting values may raise a
        `RuntimeError` or fail to iterate over all entries.
        """
        return iter(self._list)

    def __reversed__(self):
        """
        Return an iterator over the Set. Elements are iterated in their reverse
        sorted order.

        Iterating the Set while adding or deleting values may raise a
        `RuntimeError` or fail to iterate over all entries.
        """
        return reversed(self._list)

    def add(self, value):
        """Add the element *value* to the set."""
        _set = self._set
        if value not in _set:
            _set.add(value)
            self._list.add(value)

    def clear(self):
        """Remove all elements from the set."""
        self._set.clear()
        self._list.clear()

    def copy(self):
        """Create a shallow copy of the sorted set."""
        return self.__class__(key=self._key, load=self._load, _set=set(self._set))

    __copy__ = copy

    def count(self, value):
        """Return the number of occurrences of *value* in the set."""
        return 1 if value in self._set else 0

    def discard(self, value):
        """
        Remove the first occurrence of *value*.  If *value* is not a member,
        does nothing.
        """
        _set = self._set
        if value in _set:
            _set.remove(value)
            self._list.discard(value)

    def pop(self, index=-1):
        """
        Remove and return item at *index* (default last).  Raises IndexError if
        set is empty or index is out of range.  Negative indexes are supported,
        as for slice indices.
        """
        # pylint: disable=arguments-differ
        value = self._list.pop(index)
        self._set.remove(value)
        return value

    def remove(self, value):
        """
        Remove first occurrence of *value*.  Raises ValueError if
        *value* is not present.
        """
        self._set.remove(value)
        self._list.remove(value)

    def difference(self, *iterables):
        """
        Return a new set with elements in the set that are not in the
        *iterables*.
        """
        diff = self._set.difference(*iterables)
        new_set = self.__class__(key=self._key, load=self._load, _set=diff)
        return new_set

    __sub__ = difference
    __rsub__ = __sub__

    def difference_update(self, *iterables):
        """
        Update the set, removing elements found in keeping only elements
        found in any of the *iterables*.
        """
        _set = self._set
        values = set(chain(*iterables))
        if (4 * len(values)) > len(_set):
            _list = self._list
            _set.difference_update(values)
            _list.clear()
            _list.update(_set)
        else:
            _discard = self.discard
            for value in values:
                _discard(value)
        return self

    __isub__ = difference_update

    def intersection(self, *iterables):
        """
        Return a new set with elements common to the set and all *iterables*.
        """
        comb = self._set.intersection(*iterables)
        new_set = self.__class__(key=self._key, load=self._load, _set=comb)
        return new_set

    __and__ = intersection
    __rand__ = __and__

    def intersection_update(self, *iterables):
        """
        Update the set, keeping only elements found in it and all *iterables*.
        """
        _set = self._set
        _list = self._list
        _set.intersection_update(*iterables)
        _list.clear()
        _list.update(_set)
        return self

    __iand__ = intersection_update

    def symmetric_difference(self, that):
        """
        Return a new set with elements in either *self* or *that* but not both.
        """
        diff = self._set.symmetric_difference(that)
        new_set = self.__class__(key=self._key, load=self._load, _set=diff)
        return new_set

    __xor__ = symmetric_difference
    __rxor__ = __xor__

    def symmetric_difference_update(self, that):
        """
        Update the set, keeping only elements found in either *self* or *that*,
        but not in both.
        """
        _set = self._set
        _list = self._list
        _set.symmetric_difference_update(that)
        _list.clear()
        _list.update(_set)
        return self

    __ixor__ = symmetric_difference_update

    def union(self, *iterables):
        """
        Return a new SortedSet with elements from the set and all *iterables*.
        """
        return self.__class__(chain(iter(self), *iterables), key=self._key, load=self._load)

    __or__ = union
    __ror__ = __or__

    def update(self, *iterables):
        """Update the set, adding elements from all *iterables*."""
        _set = self._set
        values = set(chain(*iterables))
        if (4 * len(values)) > len(_set):
            _list = self._list
            _set.update(values)
            _list.clear()
            _list.update(_set)
        else:
            _add = self.add
            for value in values:
                _add(value)
        return self

    __ior__ = update
    _update = update

    def __reduce__(self):
        return (self.__class__, ((), self._key, self._load, self._set))

    @recursive_repr
    def __repr__(self):
        temp = '{0}({1}, key={2}, load={3})'
        return temp.format(
            self.__class__.__name__,
            repr(list(self)),
            repr(self._key),
            repr(self._load)
        )

    def _check(self):
        # pylint: disable=protected-access
        self._list._check()
        assert len(self._set) == len(self._list)
        _set = self._set
        assert all(val in _set for val in self._list)
```

### EMULSION

Epidemiological Multi-Level Simulation Framework

##### Navigation

- 1. Installation
- 2. Getting started with EMULSION
- 3. Modelling principles
- 4. Modelling language (basics)
- 5. Modelling language (advanced)
- 6. Feature examples
- 7. Information
- 8. License
- 9. High-level functions for model designers
- 10. emulsion package

##### Related Topics

- Documentation overview
  - Module code

##### Quick search

©2016, INRA and Univ. Lille.
|
Powered by Sphinx 1.8.5
& Alabaster 0.7.10
