## Supplementary material for "EMULSION: transparent and flexible multiscale stochastic models in human, animal and plant epidemiology": S2 File: emulsion.agent.html

emulsion.agent package — EMULSION (Epidemiological Multi-Level Simulation framework)

### emulsion.agent package¶

A Python implementation of EMULSION framework (Epidemiologic
MUlti-Level SImulatiONs).

This module contains classes and functions for handling all kinds of
entities involved in epidemiologic models.

#### Subpackages¶

- emulsion.agent.core package
  - Submodules
  - emulsion.agent.core.abstract\_agent module
  - emulsion.agent.core.emulsion\_agent module
  - emulsion.agent.core.groups module
- emulsion.agent.managers package
  - Submodules
  - emulsion.agent.managers.abstract\_process\_manager module
  - emulsion.agent.managers.compart\_process\_manager module
  - emulsion.agent.managers.functions module
  - emulsion.agent.managers.group\_manager module
  - emulsion.agent.managers.ibm\_process\_manager module
  - emulsion.agent.managers.metapop\_process\_manager module
  - emulsion.agent.managers.multi\_process\_manager module

#### Submodules¶

#### emulsion.agent.action module¶

A Python implementation of the EMuLSion framework (Epidemiologic
MUlti-Level SImulatiONs).

Classes and functions for actions.

*class* `emulsion.agent.action.``AbstractAction`(*state\_machine=None*, *\*\*\_*)[source]¶
:   Bases: `object`

    AbstractActions are aimed at describing actions triggered by a
    state machine.

    `__init__`(*state\_machine=None*, *\*\*\_*)[source]¶
    :   Initialize self. See help(type(self)) for accurate signature.

    *classmethod* `build_action`(*action\_name*, *\*\*others*)[source]¶
    :   Return an instance of the appropriate Action subclass,
        depending on its name. The appropriate parameters for this
        action should be passed as a dictionary.

    `execute_action`(*unit*, *\*\*others*)[source]¶
    :   Execute the action on the specified unit.

*class* `emulsion.agent.action.``BecomeAction`(*prototypes=[]*, *probas=None*, *model=None*, *\*\*others*)[source]¶
:   Bases: `emulsion.agent.action.AbstractAction`

    A BecomeAction is aimed at making an agent change its state
    according to one ore more specified prototypes.

    `__init__`(*prototypes=[]*, *probas=None*, *model=None*, *\*\*others*)[source]¶
    :   Initialize self. See help(type(self)) for accurate signature.

    `execute_action`(*unit*, *agents=None*, *\*\*others*)[source]¶
    :   Execute the action in the specified unit. If the agents parameter
        is specified (as a list), each agent of this list will execute
        the action. If changes of state variables in relation to a
        state machine occur, the corresponding actions (if any) are
        executed: on\_exit from the current state, and on\_enter for the
        new state.

*class* `emulsion.agent.action.``CloneAction`(*prototypes=[]*, *amount=None*, *probas=None*, *model=None*, *\*\*others*)[source]¶
:   Bases: `emulsion.agent.action.AbstractAction`

    A CloneAction produces several copies of the agent with a given
    prototype.

    `__init__`(*prototypes=[]*, *amount=None*, *probas=None*, *model=None*, *\*\*others*)[source]¶
    :   Initialize self. See help(type(self)) for accurate signature.

    `execute_action`(*unit*, *agents=None*, *\*\*others*)[source]¶
    :   Execute the action in the specified unit. If the agents parameter
        is specified (as a list), each agent of this list will execute
        the action. If changes of state variables in relation to a
        state machine occur, the corresponding actions (if any) are
        executed: on\_exit from the current state, and on\_enter for the
        new state.

*class* `emulsion.agent.action.``FunctionAction`(*function=None*, *\*\*others*)[source]¶
:   Bases: `emulsion.agent.action.MethodAction`

    A FunctionAction is aimed at making an agent perform an action
    on a specific population. It requires a function, and optionnally
    a list and a dictionary of parameters. A FunctionAction runs
    faster than a MethodAction since it does not require to retrieve
    the method in each agent.

    `__init__`(*function=None*, *\*\*others*)[source]¶
    :   Initialize self. See help(type(self)) for accurate signature.

    `execute_action`(*unit*, *agents=None*, *\*\*others*)[source]¶
    :   Execute the action using the specified unit. If the
        agents parameter is a list of units, each unit of this list
        will execute the action.

*exception* `emulsion.agent.action.``InvalidActionException`(*message*)[source]¶
:   Bases: `Exception`

    Exception raised when a semantic error occurs in action definition.

    `__init__`(*message*)[source]¶
    :   Initialize self. See help(type(self)) for accurate signature.

*class* `emulsion.agent.action.``LogVarsAction`(*parameter=None*, *l\_params=[]*, *d\_params={}*, *\*\*others*)[source]¶
:   Bases: `emulsion.agent.action.StringAction`

    A LogVarsAction is aimed at making an agent print a list of
    variables into a file. Output is formatted in several
    comma-separated fields: the simulation ID, the time step when the
    message was produced, the class and ID of agent speaking, and the
    values of variables (in the order they were defined in the
    action).

    `execute_action`(*unit*, *agents=None*, *\*\*others*)[source]¶
    :   Execute the action in the specified unit. If the agents parameter
        is specified (as a list), each agent of this list will execute
        the action.

*class* `emulsion.agent.action.``MessageAction`(*parameter=None*, *l\_params=[]*, *d\_params={}*, *\*\*others*)[source]¶
:   Bases: `emulsion.agent.action.StringAction`

    A MessageAction is aimed at making an agent print a given
    string. It requires a string message. This string can contain one
    reference to a variable or method of the agent, using Python’s
    formatting syntax.

    For instance, ‘My state is {.statevars.health\_state}’ will print
    the current health state of the agent.

    Output is formatted in four comma-separated fields: the simulation
    ID, the time step when the message was produced, the agent
    speaking, and the message itself.

    `execute_action`(*unit*, *agents=None*, *\*\*others*)[source]¶
    :   Execute the action in the specified unit. If the agents parameter
        is specified (as a list), each agent of this list will execute
        the action.

*class* `emulsion.agent.action.``MethodAction`(*method=None*, *l\_params=[]*, *d\_params={}*, *\*\*others*)[source]¶
:   Bases: `emulsion.agent.action.AbstractAction`

    A MethodAction is aimed at making an agent perform an action on
    a specific population. It requires a method name, and optionnally
    a list and a dictionary of parameters.

    `__init__`(*method=None*, *l\_params=[]*, *d\_params={}*, *\*\*others*)[source]¶
    :   Initialize self. See help(type(self)) for accurate signature.

    `execute_action`(*unit*, *agents=None*, *\*\*others*)[source]¶
    :   Execute the action using the specified unit. If the
        agents parameter is a list of units, each unit of this list
        will execute the action.

*class* `emulsion.agent.action.``RateAdditiveAction`(*sign=1*, *\*\*others*)[source]¶
:   Bases: `emulsion.agent.action.ValueAction`

    A RateChangeAction is aimed at increasing or decreasing a
    specific state variable or attribute, according to a specific rate
    (i.e. the actual increase or decrease is the product of the
    parameter attribute and a population size).

    `__init__`(*sign=1*, *\*\*others*)[source]¶
    :   Create a ValueAction aimed at modifying the specified
        statevar according to the parameter.

    `execute_action`(*unit*, *population=None*, *agents=None*)[source]¶
    :   Execute the action on the specified unit, with the
        specified population size.

*class* `emulsion.agent.action.``RateDecreaseAction`(*\*\*others*)[source]¶
:   Bases: `emulsion.agent.action.RateAdditiveAction`

    A RateDecreaseAction is aimed at decreasing a specific state
    variable or attribute, according to a specific rate (i.e. the
    actual decrease is the product of the parameter attribute and a
    population size).

    `__init__`(*\*\*others*)[source]¶
    :   Create a ValueAction aimed at modifying the specified
        statevar according to the parameter.

*class* `emulsion.agent.action.``RateIncreaseAction`(*\*\*others*)[source]¶
:   Bases: `emulsion.agent.action.RateAdditiveAction`

    A RateIncreaseAction is aimed at increasing a specific state
    variable or attribute, according to a specific rate (i.e. the
    actual increase is the product of the parameter attribute and a
    population size).

    `__init__`(*\*\*others*)[source]¶
    :   Create a ValueAction aimed at modifying the specified
        statevar according to the parameter.

*class* `emulsion.agent.action.``RecordChangeAction`(*parameter=None*, *model=None*, *\*\*others*)[source]¶
:   Bases: `emulsion.agent.action.ValueAction`

    RecordChangeAction allows to record how many agents performed an
    action set. The corresponding value is added to a variable assumed
    to be defined at the upper level.

    `__init__`(*parameter=None*, *model=None*, *\*\*others*)[source]¶
    :   Create a RecordChangeAction aimed at modifying the specified
        parameter, (assumed to be a statevar defined at the upper
        level).

    `execute_action`(*unit*, *population=None*, *agents=None*)[source]¶
    :   Execute the action on the specified unit, with the
        specified population size.

*class* `emulsion.agent.action.``SetVarAction`(*statevar\_name=None*, *parameter=None*, *model=None*, *\*\*others*)[source]¶
:   Bases: `emulsion.agent.action.ValueAction`

    SetVarAction allows to set the variable of the agent.

    `__init__`(*statevar\_name=None*, *parameter=None*, *model=None*, *\*\*others*)[source]¶
    :   Create a SetVarAction aimed at modifying the specified statevar
        according to the parameter.

    `execute_action`(*unit*, *agents=None*, *\*\*others*)[source]¶
    :   Execute the action in the specified unit. If the agents parameter
        is specified (as a list), each agent of this list will execute
        the action. If changes of state variables in relation to a
        state machine occur, the corresponding actions (if any) are
        executed: on\_exit from the current state, and on\_enter for the
        new state.

*class* `emulsion.agent.action.``StochAdditiveAction`(*sign=1*, *\*\*others*)[source]¶
:   Bases: `emulsion.agent.action.ValueAction`

    A StochAdditiveAction is aimed at increasing or decreasing a
    specific state variable or attribute, according to a specific
    rate, using a *binomial sampling*.

    `__init__`(*sign=1*, *\*\*others*)[source]¶
    :   Create a ValueAction aimed at modifying the specified
        statevar according to the parameter.

    `execute_action`(*unit*, *population=None*, *agents=None*)[source]¶
    :   Execute the action on the specified unit, with the
        specified population size.

*class* `emulsion.agent.action.``StochDecreaseAction`(*\*\*others*)[source]¶
:   Bases: `emulsion.agent.action.StochAdditiveAction`

    A StochDecreaseAction is aimed at decreasing a specific state
    variable or attribute, according to a specific rate, using a
    *binomial sampling*.

    `__init__`(*\*\*others*)[source]¶
    :   Create a ValueAction aimed at modifying the specified
        statevar according to the parameter.

*class* `emulsion.agent.action.``StochIncreaseAction`(*\*\*others*)[source]¶
:   Bases: `emulsion.agent.action.StochAdditiveAction`

    A StochIncreaseAction is aimed at increasing a specific state
    variable or attribute, according to a specific rate, using a
    *binomial sampling*.

    `__init__`(*\*\*others*)[source]¶
    :   Create a ValueAction aimed at modifying the specified
        statevar according to the parameter.

*class* `emulsion.agent.action.``StringAction`(*parameter=None*, *l\_params=[]*, *d\_params={}*, *\*\*others*)[source]¶
:   Bases: `emulsion.agent.action.AbstractAction`

    A StringAction is based on the specification of a string
    parameter.

    `__init__`(*parameter=None*, *l\_params=[]*, *d\_params={}*, *\*\*others*)[source]¶
    :   Initialize self. See help(type(self)) for accurate signature.

*class* `emulsion.agent.action.``ValueAction`(*statevar\_name=None*, *parameter=None*, *delta\_t=1*, *\*\*others*)[source]¶
:   Bases: `emulsion.agent.action.AbstractAction`

    ValueActions represent modifications of state variables or
    attributes.

    `__init__`(*statevar\_name=None*, *parameter=None*, *delta\_t=1*, *\*\*others*)[source]¶
    :   Create a ValueAction aimed at modifying the specified
        statevar according to the parameter.

#### emulsion.agent.atoms module¶

A Python implementation of the EMuLSion framework (Epidemiologic
MUlti-Level SImulatiONs).

Classes and functions for entities management.

*class* `emulsion.agent.atoms.``AtomAgent`(*\*\*others*)[source]¶
:   Bases: `emulsion.agent.core.emulsion_agent.EmulsionAgent`

    The AtomAgent is aimed at representing an ‘individual’, i.e. the
    smallest organization level to be modeled as an entity in the
    simulation. An AtomAgent may be situated in several hosts, each one
    associated with a specific tuple of state variables.

    `__init__`(*\*\*others*)[source]¶
    :   Initialize the unit with a health state and a name.

    `add_host`(*host*)[source]¶
    :   Add the specified host to the current AtomAgent, associated
        with the specified key.

    `agcount` *= 0*¶

    `agdict` *= {}*¶

    `clone`(*prototype=None*, *custom\_prototype=None*, *\*\*others*)[source]¶
    :   Make a copy of the current compartment with the specified
        observable/value settings. If a prototype is provided, it is
        applied to the new atom.

    `families` *= SortedSet(['AtomAgent'], key=None, load=1000)*¶

    `get_content`()[source]¶
    :   Return the population (1) of the current unit.

    `get_host`(*key='MASTER'*)[source]¶
    :   Retrieve the host of the current AtomAgent identified by the
        specific key.

    `members` *= ('agcount', 'agdict', 'families', '\_\_module\_\_', '\_\_qualname\_\_', '\_\_doc\_\_', '\_\_init\_\_', '\_\_len\_\_', 'get\_content', 'add\_host', 'remove\_host', 'get\_host', 'clone', '\_\_classcell\_\_')*¶

    `remove_host`(*host*, *keys=None*)[source]¶
    :   Remove the specified host from the current AtomAgent,
        associated with the specified key.

*class* `emulsion.agent.atoms.``EvolvingAtom`(*statemachines=[]*, *\*\*others*)[source]¶
:   Bases: `emulsion.agent.atoms.AtomAgent`

    An EvolvingAtom is able to change state according to its
    own statemachines.

    `__init__`(*statemachines=[]*, *\*\*others*)[source]¶
    :   Initialize the unit with a health state and a name.

    `add_method_process`(*process\_name*, *method=None*)[source]¶
    :   Add a process based on a method name.

    `agcount` *= 0*¶

    `agdict` *= {}*¶

    `evolve`(*machine=None*)[source]¶
    :   This method is aimed at defining what has systematically to
        be done in the unit at each time step (e.g. age change…). It
        has to be overriden if needed in subclasses.

    `evolve_states`(*machine=None*)[source]¶
    :   Change the state of the current unit according to the
        specified state machine name. If no special state machine is
        provided, executes all the machines.

    `families` *= SortedSet(['EvolvingAtom'], key=None, load=1000)*¶

    `get_machine`(*name*)[source]¶
    :   Return the state machine with the specified name.

    `init_level_processes`()[source]¶
    :   Initialize the level of the agent.

    `members` *= ('agcount', 'agdict', 'families', '\_\_module\_\_', '\_\_qualname\_\_', '\_\_doc\_\_', '\_\_init\_\_', 'set\_statemachines', 'init\_level\_processes', 'add\_method\_process', 'get\_machine', 'evolve', 'evolve\_states', '\_\_classcell\_\_')*¶

    `set_statemachines`(*statemachines*)[source]¶
    :   Define the state machines that this agent is able to execute.

#### emulsion.agent.comparts module¶

A Python implementation of the EMuLSion framework (Epidemiologic
MUlti-Level SImulatiONs).

Classes and functions for entities management.

*class* `emulsion.agent.comparts.``Compartment`(*population=0*, *stochastic=True*, *\*\*others*)[source]¶
:   Bases: `emulsion.agent.core.groups.GroupAgent`

    An Compartment is a compartment which does not
    represent the underlying level but with aggregate information such
    as the total population (‘individuals’ are not represented).

    `__init__`(*population=0*, *stochastic=True*, *\*\*others*)[source]¶
    :   Create an Compartment with an initial population.

    `add`(*population*)[source]¶
    :   Add the specified population to the current population of
        the compartment.

    `agcount` *= 0*¶

    `agdict` *= {}*¶

    `clone`(*\*\*others*)[source]¶
    :   Make a copy of the current compartment with the specified
        observable/value settings. The new content is empty.

    `families` *= SortedSet(['Compartment'], key=None, load=1000)*¶

    `get_content`()[source]¶
    :   Return the population of the current unit.

    `members` *= ('agcount', 'agdict', 'families', '\_\_module\_\_', '\_\_qualname\_\_', '\_\_doc\_\_', '\_\_init\_\_', '\_\_len\_\_', 'get\_content', 'add', 'remove', '\_base\_move', 'move\_to', 'population', 'clone', 'next\_states', '\_\_classcell\_\_')*¶

    `move_to`(*other\_unit*, *population*, *state\_machine=None*, *\*\*others*)[source]¶
    :   Move the specified population from the current population
        of the compartment (the population is kept positive) to the
        other unit. If a state machine is provided, executes the
        corresponding actions when entering/exiting nodes and crossing
        edges if needed.

    `next_states`(*states*, *values*, *populations*, *actions*, *method=None*)[source]¶
    :   Compute the population moving from the current compartment to each
        of the destination states, handling the values according the
        the specified method. Values can be handled either as absolute
        amounts (‘amount’ method), as proportions (‘rate’, in a
        deterministic approach) or as probabilities (‘proba’, in a
        stochastic approach). Actions are to be performed when
        changing state. The actual population affected by the
        transitions is stored in the first element of the
        populations parameter, as a dictionary: {‘population’:
        number, ‘actions’: actions}. Several edges can lead to the
        same state.

        Return a list of tuples:
        :   (state, {‘population’: qty, ‘actions:’ list of actions})

    `population`¶

    `remove`(*population*)[source]¶
    :   Remove the specified population from the current population
        of the compartment (the population is kept positive).

#### emulsion.agent.exceptions module¶

A Python implementation of the EMuLSion framework (Epidemiologic
MUlti-Level SImulatiONs).

Exceptions raised by agents.

*exception* `emulsion.agent.exceptions.``InvalidCompartmentOperation`(*source*, *operation*, *params*)[source]¶
:   Bases: `Exception`

    Exception raised when a compartiment is asked for impossible
    operations, such as adding numbers to a list of units.

    `__init__`(*source*, *operation*, *params*)[source]¶
    :   Initialize self. See help(type(self)) for accurate signature.

*exception* `emulsion.agent.exceptions.``LevelException`(*cause*, *level*)[source]¶
:   Bases: `Exception`

    Exception raised when a semantic error occurs during model parsing.

    `__init__`(*cause*, *level*)[source]¶
    :   Initialize self. See help(type(self)) for accurate signature.

*exception* `emulsion.agent.exceptions.``StateVarNotFoundException`(*statevar*, *source*)[source]¶
:   Bases: `Exception`

    Exception raised when a semantic error occurs during model parsing.

    `__init__`(*statevar*, *source*)[source]¶
    :   Initialize self. See help(type(self)) for accurate signature.

#### emulsion.agent.meta module¶

A Python implementation of the EMuLSion framework (Epidemiologic
MUlti-Level SImulatiONs).

Classes and functions for abstract agent management.

Part of this code was adapted from the PADAWAN framework (S. Picault,
Univ. Lille).

*class* `emulsion.agent.meta.``MetaAgent`[source]¶
:   Bases: `abc.ABCMeta`

    The Metaclass definition for all agents. When created, agents
    are stored in a class-specific dictionaries of agents. They are
    given an ID value (unique value within each class) and can be
    assigned to several agents families (by default, each agent is
    assigned to its own class).

#### emulsion.agent.process module¶

A Python implementation of the EMuLSion framework (Epidemiologic
MUlti-Level SImulatiONs).

Classes and functions for process management in MultiProcessCompartments.

*class* `emulsion.agent.process.``AbstractProcess`(*name*)[source]¶
:   Bases: `object`

    An AbstractProcess is aimed at controlling a specific activity
    in a compartment, and is identified by its name.

    `__init__`(*name*)[source]¶
    :   Initialize self. See help(type(self)) for accurate signature.

    `evolve`()[source]¶
    :   Define the actions that the process must perform.

*class* `emulsion.agent.process.``MethodProcess`(*name*, *method*, *lparams=[]*, *dparams={}*)[source]¶
:   Bases: `emulsion.agent.process.AbstractProcess`

    A MethodProcess is aimed at running a specific method (and
    possibly any function or even any callable object).

    `__init__`(*name*, *method*, *lparams=[]*, *dparams={}*)[source]¶
    :   Initialize self. See help(type(self)) for accurate signature.

    `evolve`()[source]¶
    :   Define the actions that the process must perform. In a
        MethodProcess, those actions consist in running the method of
        the target compartment.

*class* `emulsion.agent.process.``StateMachineProcess`(*name*, *agent*, *state\_machine*)[source]¶
:   Bases: `emulsion.agent.process.AbstractProcess`

    A StateMachineProcess is aimed at running a specific state machine
    within the agent (not within a grouping).

    `__init__`(*name*, *agent*, *state\_machine*)[source]¶
    :   Initialize self. See help(type(self)) for accurate signature.

    `evolve`()[source]¶
    :   Define the actions that the process must perform. In a
        StateMachineProcess, those actions consist in 1) executing the
        transitions of the state machine to change the agent’s states
        and 2) computing productions and transmit them to the upper
        level of the agent.

    `evolve_productions`()[source]¶

    `evolve_transitions`()[source]¶

#### emulsion.agent.views module¶

A Python implementation of the EMuLSion framework (Epidemiologic
MUlti-Level SImulatiONs).

Classes and functions for entities management.

*class* `emulsion.agent.views.``AdaptiveView`(*observables=(None*, *)*, *values=(None*, *)*, *\*\*others*)[source]¶
:   Bases: `emulsion.agent.views.SimpleView`

    An AdaptiveView is able to evaluate conditions on individuals
    (AtomAgents) for state transitions, and to detect changes in its
    content and automatically report it to the upper level. Each
    compartment is associated with a particular value of a specific
    state variable or attribute of the units in its content. At the
    end of the evolve step, the compartment checks the values of the
    units in its content and reports units where changes occured.

    `__init__`(*observables=(None*, *)*, *values=(None*, *)*, *\*\*others*)[source]¶
    :   Specify the state variables or attributes associated with
        this compartment (`observables`) and the expected values.

    `agcount` *= 0*¶

    `agdict` *= {}*¶

    `check_consistency`()[source]¶
    :   Check the value of the observables of the current
        compartment. Units which do not have the expected values are
        notified to the upper level.

    `clone`(*\*\*others*)[source]¶
    :   Make a copy of the current compartment with the specified
        observable/value settings. The new content is empty.

    `evaluate_condition`(*condition*)[source]¶
    :   Return the population of the compartment if the condition
        is fulfilled, 0 otherwise.

    `evolve`(*machine=None*)[source]¶
    :   After the ordinary `evolve` step, check units which have
        changed their value(s) of the compartment-specific
        observable(s). The corresponding units are reported to the
        upper level.

    `families` *= SortedSet(['AdaptiveView'], key=None, load=1000)*¶

    `members` *= ('agcount', 'agdict', 'families', '\_\_module\_\_', '\_\_qualname\_\_', '\_\_doc\_\_', '\_\_init\_\_', 'evaluate\_condition', 'next\_states', 'evolve', 'check\_consistency', 'clone', '\_\_classcell\_\_')*¶

    `next_states`(*states*, *values*, *populations*, *actions*, *method=None*)[source]¶
    :   Compute stochastically the population moving from each population
        (tuples (‘agents’: list\_of\_agents)) in the populations
        parameter, to each of the possible destination states,
        according to each of the values interpreted as
        probabilities. Actions are to be performed when changing
        state. Several edges can lead to the same state.

        Return a list of tuples:
        :   (state, {‘agent’: list of agents, ‘actions’: list of actions})

*class* `emulsion.agent.views.``AutoStructuredView`(*key\_variable=None*, *\*\*others*)[source]¶
:   Bases: `emulsion.agent.views.StructuredView`

    An AutoStructuredView stores agents in an OrderedDict, using a
    specific statevar as key.

    `__init__`(*key\_variable=None*, *\*\*others*)[source]¶
    :   Initialize the unit with a health state and a name.

    `add`(*population*)[source]¶
    :   Add the specified population (SortedSet or list) to the
        current view.

    `agcount` *= 0*¶

    `agdict` *= {}*¶

    `families` *= SortedSet(['AutoStructuredView'], key=None, load=1000)*¶

    `members` *= ('agcount', 'agdict', 'families', '\_\_module\_\_', '\_\_qualname\_\_', '\_\_doc\_\_', '\_\_init\_\_', 'add', 'remove', '\_\_classcell\_\_')*¶

    `remove`(*population*)[source]¶
    :   Remove the specified population from the current view. The
        population is expected to be a SortedSet or list.

*class* `emulsion.agent.views.``SimpleView`(*\*\*others*)[source]¶
:   Bases: `emulsion.agent.core.groups.Aggregation`

    A SimpleView uses a set to store the underlying
    units. It is rather aimed at storing AtomAgents. All units are
    considered in the same state, thus conditions in the state
    machines are evaluated for the whole compartment, not for
    individuals.

    `__init__`(*\*\*others*)[source]¶
    :   Initialize the unit with a health state and a name.

    `add`(*population*)[source]¶
    :   Add the specified population to the current compartment.

    `agcount` *= 0*¶

    `agdict` *= {}*¶

    `families` *= SortedSet(['SimpleView'], key=None, load=1000)*¶

    `get_content`()[source]¶
    :   Return the units contained in the current unit.

    `members` *= ('agcount', 'agdict', 'families', '\_\_module\_\_', '\_\_qualname\_\_', '\_\_doc\_\_', '\_\_init\_\_', '\_\_iter\_\_', 'get\_content', 'add', 'remove', 'next\_states', '\_\_classcell\_\_')*¶

    `next_states`(*states*, *values*, *populations*, *actions*, *method=None*)[source]¶
    :   Compute stochastically the population moving from the current
        compartment to each of the destination states, according to
        the values interpreted as probabilities. Actions are to be
        performed when changing state. The population affected by the
        transitions is stored in the first element of the
        populations parameter, as a tuple: (‘agents’,
        list\_of\_agents). Several edges can lead to the same state.

        Return a list of tuples:
        :   (state, {‘agent’: list of agents, ‘actions’: list of actions})

    `remove`(*population*)[source]¶
    :   Remove the specified population from the current
        compartment.

*class* `emulsion.agent.views.``StructuredView`(*keep\_history=True*, *\*\*others*)[source]¶
:   Bases: `emulsion.agent.core.groups.Aggregation`

    A StructuredView uses ad dict list to store the underlying
    units. It is rather aimed at storing other compartiments.

    `__init__`(*keep\_history=True*, *\*\*others*)[source]¶
    :   Initialize the unit with a health state and a name.

    `add`(*population*)[source]¶
    :   Add the specified population to the current
        compartment. The population is expected to be a dictionary
        with names as keys and compartments as values.

    `agcount` *= 0*¶

    `agdict` *= {}*¶

    `evolve`(*machine=None*)[source]¶
    :   Ask each unit in the current compartment to make its content evolve
        according to its own capabilities. A specific state machine
        can be specified if needed.

    `families` *= SortedSet(['StructuredView'], key=None, load=1000)*¶

    `get_content`()[source]¶
    :   Return the agents contained in the current view.

    `get_or_build`(*key*, *source=None*)[source]¶
    :   Return the compartment with the specified key if any, or
        build one by cloning the source if not found.

    `handle_notifications`()[source]¶
    :   Handle all notifications received during the time step.

    `members` *= ('agcount', 'agdict', 'families', '\_\_module\_\_', '\_\_qualname\_\_', '\_\_doc\_\_', '\_\_init\_\_', '\_\_iter\_\_', '\_\_getitem\_\_', 'get\_content', 'add', 'remove', 'notify\_changed\_units', 'handle\_notifications', 'get\_or\_build', 'evolve', '\_\_classcell\_\_')*¶

    `notify_changed_units`(*source\_compartment*, *units*)[source]¶
    :   Receive a change notification concerning several units
        which cannot be stored anymore in the source compartment.

    `remove`(*population*)[source]¶
    :   Remove the specified population from the current
        compartment. The population is expected to be a dictionary
        with names as keys and compartments as values.

*class* `emulsion.agent.views.``StructuredViewWithCounts`(*allowed\_values=None*, *\*\*others*)[source]¶
:   Bases: `emulsion.agent.views.StructuredView`

    Same as StructuredView, plus the capability of handling counts for
    key values.

    `__init__`(*allowed\_values=None*, *\*\*others*)[source]¶
    :   Initialize the unit with a health state and a name.

    `agcount` *= 0*¶

    `agdict` *= {}*¶

    `evolve`()[source]¶
    :   Ask each unit in the current compartment to make its content evolve
        according to its own capabilities. A specific state machine
        can be specified if needed.

    `families` *= SortedSet(['StructuredViewWithCounts'], key=None, load=1000)*¶

    `init_counts`(*index=0*)[source]¶
    :   Initialize the counts.

    `members` *= ('agcount', 'agdict', 'families', '\_\_module\_\_', '\_\_qualname\_\_', '\_\_doc\_\_', '\_\_init\_\_', 'init\_counts', 'update\_counts', '\_\_contains\_\_', 'evolve', '\_\_classcell\_\_')*¶

    `update_counts`(*index=0*)[source]¶
    :   Update the counts.

### EMULSION

Epidemiological Multi-Level Simulation Framework

##### Navigation

- 1. Installation
- 2. Getting started with EMULSION
- 3. Modelling principles
- 4. Modelling language (basics)
- 5. Modelling language (advanced)
- 6. Feature examples
- 7. Information
- 8. License
- 9. High-level functions for model designers
- 10. emulsion package
  - 10.1. Subpackages
    - emulsion.agent package
    - emulsion.model package
    - emulsion.tools package
  - 10.2. Submodules
  - 10.3. emulsion.init\_emulsion module

##### Related Topics

- Documentation overview
  - 10. emulsion package
    - Previous: 10. emulsion package
    - Next: emulsion.agent.core package

##### Quick search

©2016, INRA and Univ. Lille.
|
Powered by Sphinx 1.8.5
& Alabaster 0.7.10
|
Page source
