## Supplementary material for "EMULSION: transparent and flexible multiscale stochastic models in human, animal and plant epidemiology": S2 File: emulsion.agent.managers.html

emulsion.agent.managers package — EMULSION (Epidemiological Multi-Level Simulation framework)

### emulsion.agent.managers package¶

*Module author: Sébastien Picault <>*

#### Submodules¶

#### emulsion.agent.managers.abstract\_process\_manager module¶

*Module author: Sébastien Picault <>*

*class* `emulsion.agent.managers.abstract_process_manager.``AbstractProcessManager`(*model=None*, *master=None*, *level=None*, *stochastic=True*, *keep\_history=False*, *prototype=None*, *custom\_prototype=None*, *execute\_actions=False*, *\*\*others*)[source]¶
:   Bases: `emulsion.agent.views.StructuredView`

    An AbstractProcessManager is aimed handling several independent
    StructuredViews at the same time, to represent several
    concerns. It can automatically build compartments for state
    machines associated with a specific state variable or attribute.

    `__init__`(*model=None*, *master=None*, *level=None*, *stochastic=True*, *keep\_history=False*, *prototype=None*, *custom\_prototype=None*, *execute\_actions=False*, *\*\*others*)[source]¶
    :   Initialize the unit with a health state and a name.

    `add_compart_process`(*process\_name*, *key\_variables*, *compart\_manager=(<class 'emulsion.agent.managers.group\_manager.GroupManager'>*, *{})*, *machine\_name=None*, *allowed\_values=None*, *compart\_class=(<class 'emulsion.agent.views.AdaptiveView'>*, *{})*)[source]¶
    :   Add a process aimed at managing a ‘Compartment Manager’, i.e. an
        object aimed at managing a collection of compartments. This
        compartment manager is automatically initialized from the
        compart\_manager class (which should be a subclass of
        StructuredView or GroupManager). The compartment manager may
        be associated with a specific state machine, and MUST BE
        identified by a tuple of state variables names. Additionally,
        since a first compartment is also instantiated, a specific
        class to do so can be also specified.

    `add_method_process`(*process\_name*, *method=None*)[source]¶
    :   Add a process based on a method name.

    `add_new_population`(*process\_name*, *population*)[source]¶

    `add_statemachine_process`(*process\_name*)[source]¶
    :   Add a process based on the direct execution of a state machine.

    `agcount` *= 0*¶

    `agdict` *= {}*¶

    `apply_initial_conditions`()[source]¶
    :   Apply initial conditions (if any) as defined in the model.

    `apply_initial_prototype`(*name=None*, *prototype=None*, *execute\_actions=False*)[source]¶
    :   This method inherited from AbstractAgent and called by new\_atom
        is intentionnaly doing nothing in the current class to ensure
        that initial prototype of a MultiProcessManager is applied
        before creating the sublevels.

    `counts`¶
    :   Return a pandas DataFrame containing counts of each process if existing.
        TODO: column steps need to be with one of process

    `create_count_properties_for_state`(*process\_or\_machine\_name*, *state\_name*, *count\_function*, *aggregation\_function*)[source]¶
    :   Dynamically add properties of the form `total_S` where S can be
        any state of the state machine. Counts are expected to be
        computed by the grouping associated with the specified
        process. The access to counts is defined by
        *count\_function*. If aggregated variables such as `aggvar`
        are defined, corresponding properties of the form
        ``aggvar\_S``are also defined with the specified
        *aggregation\_function*.

    `create_properties_for_groups`(*process\_name*, *key\_variables*)[source]¶
    :   Dynamically add properties of the form total\_S\_T where S, T are a
        valid key in grouping associated with the specified
        process\_name.

    `evolve`(*\*\*others*)[source]¶
    :   Make the ProcessManager evolve, i.e. all the registered processes
        in order, starting with the evolution of the sublevels, and
        followed by the evolution inherited from superclasses.

    `families` *= SortedSet(['AbstractProcessManager'], key=None, load=1000)*¶

    `finalize_level`(*simulation=None*, *\*\*others*)[source]¶
    :   User-defined operations at the end of simulation for an instance of
        this level.

    `get_group_population`(*process\_name*, *group\_name*)[source]¶
    :   Return the size of the subgroup associated with the specified group
        name (state names).

    `init_processes`()[source]¶
    :   Init the processes that the ProcessManager will undergo during each
        time step, in order. Processes may be either ‘method’
        processes (based on the execution of the specified method
        name), or ‘group-based’ processes (defined by the evolution of
        a grouping (aggregation or compartment), possibly endowed with
        a state machine), or even a ‘state-machine-driven’ process,
        based on the direct execution of a state machine within the
        ProcessManager.

    `initialize_level`(*\*\*others*)[source]¶
    :   User-defined operations when creating an instance of this
        level. These operations are performed *after* the application of
        initial conditions possibly defined in the corresponding model
        section.

    `load_state_from_file`(*simu\_id*, *filename*)[source]¶

    `members` *= ('agcount', 'agdict', 'families', '\_\_module\_\_', '\_\_qualname\_\_', '\_\_doc\_\_', '\_\_init\_\_', 'load\_state\_from\_file', 'save\_state\_to\_file', 'apply\_initial\_prototype', 'apply\_initial\_conditions', 'initialize\_level', 'finalize\_level', 'add\_new\_population', 'init\_processes', 'evolve', 'add\_method\_process', 'add\_statemachine\_process', 'add\_compart\_process', 'create\_count\_properties\_for\_state', 'create\_properties\_for\_groups', 'get\_group\_population', 'counts', 'remove\_randomly', 'remove', 'population', '\_\_classcell\_\_')*¶

    `population`¶
    :   Return the total population of the compartment. It is
        calculated either using a true ‘population’ statevar if any,
        or as the sum of the population of each unit contained in the
        compartment.

        TAG: USER

    `remove`(*agents\_or\_population*)[source]¶
    :   Remove the specified population from the current
        compartment. The population is expected to be a dictionary
        with names as keys and compartments as values.

    `remove_randomly`(*proba=0*)[source]¶
    :   Remove randomly chosen atoms or population from this
        ProcessManager.

    `save_state_to_file`(*simu\_id*, *filename*)[source]¶

#### emulsion.agent.managers.compart\_process\_manager module¶

*Module author: Sébastien Picault <>*

*class* `emulsion.agent.managers.compart_process_manager.``CompartProcessManager`(*model=None*, *master=None*, *level=None*, *stochastic=True*, *keep\_history=False*, *prototype=None*, *custom\_prototype=None*, *execute\_actions=False*, *\*\*others*)[source]¶
:   Bases: `emulsion.agent.managers.abstract_process_manager.AbstractProcessManager`

    A CompartProcessManager is aimed handling several independent
    StructuredViews at the same time, for managing true compartments.
    It can automatically allocate compartments for state machines
    associated with a specific state variable or attribute.

    `add_compart_process`(*process\_name*, *key\_variables*, *compart\_manager=(<class 'emulsion.agent.managers.group\_manager.GroupManager'>*, *{})*, *machine\_name=None*, *compart\_class=(<class 'emulsion.agent.comparts.Compartment'>*, *{})*)[source]¶
    :   Add a process aimed at managing a ‘Compartment Manager’, i.e. an
        object aimed at managing a collection of compartments. This
        compartment manager is automatically initialized from the
        compart\_manager class (which should be a subclass of
        StructuredView or GroupManager). The compartment manager may
        be associated with a specific state machine, and MUST BE
        identified by a tuple of state variables names. Additionally,
        since a first compartment is also instantiated, a specific
        class to do so can be also specified.

    `add_host`(*host*)[source]¶
    :   Add the specified host to the current Multiprocessmanager, associated
        with the specified key.

    `add_new_population`(*process\_name*, *population*)[source]¶

    `add_population`(*population\_spec*, *init=False*)[source]¶
    :   Add the specified population specification to the current
        CompartProcessManager. population\_spec is a dictionary with
        process names as keys, each one associated with a dictionary
        (tuple of statevars) -> population. If init is True, the compartment
        managers counts the initial value of the populations in each
        compartment.

    `agcount` *= 0*¶

    `agdict` *= {}*¶

    `apply_initial_conditions`()[source]¶
    :   Initialize level with initial conditions specified in the model.

        As this agent is aimed at managing aggregated populations in
        the sub-levels, only ‘populations’ are taken into account to
        initialize the sub-levels.

    `complement_population`(*population\_to\_change*, *total=0*, *remove=False*)[source]¶
    :   Modifiy the population spec in a consistent way. Check is all
        populations in each process sum to the same total. If not, distribute
        the difference between total and sum at random between all available
        groups (FAIL if no available groups).

    `counts`¶
    :   Return a pandas DataFrame containing counts of each process if existing.
        TODO: column steps need to be with one of process

    `families` *= SortedSet(['CompartProcessManager'], key=None, load=1000)*¶

    `members` *= ('agcount', 'agdict', 'families', '\_\_module\_\_', '\_\_qualname\_\_', '\_\_doc\_\_', 'add\_compart\_process', 'add\_host', 'apply\_initial\_conditions', 'add\_new\_population', 'complement\_population', 'add\_population', 'remove', 'remove\_population', 'remove\_randomly', 'counts', '\_\_classcell\_\_')*¶

    `remove`(*agents\_or\_population*)[source]¶
    :   Remove the specified population from the current
        compartment. The population is expected to be a dictionary
        with names as keys and compartments as values.

    `remove_population`(*population\_spec*)[source]¶
    :   Remove the specified population spec from the current
        CompartProcessManager.

    `remove_randomly`(*proba=0*, *amount=None*, *process=None*)[source]¶
    :   Remove random amounts of populations from this ProcessManager. If
        amount is not None, a multinomial sampling is performed for
        each process. Otherwise: proba can be either a probability
        or a dictionary. In that case, the process parameter
        indicates the name of the process grouping which drives the
        probabilities, and the keys must be those of the
        grouping. Selected quantities are removed and returned by the
        method.

#### emulsion.agent.managers.functions module¶

*Module author: Sébastien Picault <>*

`emulsion.agent.managers.functions.``group_and_split_populations`(*transitions*)[source]¶
:   Transform a list of transitions into a dictionary based on the
    underlying populations.

    transitions is a list of tuples (state, flux, value,
    cond\_result, actions) where:
    - state is a possible state reachable from the current state flux
    - is either ‘rate’ or ‘proba’ or ‘amount’ or ‘amount-all-but’
    - value is the corresponding rate or probability or amount
    - cond\_result is a couple (either (‘population’, qty) or (‘agents’, list))

    > describing who fulfills the condition to cross the transition

    - actions is the list of actions on cross

    The goal of this function is to restructure all those elements for
    disjoint sub-populations.

    Return a list of tuples:
    :   either ((‘population’, qty), attributes)
        or ((‘agents’, list), attributes)

    where attributes is a list of tuples (state, flux, value, actions)

#### emulsion.agent.managers.group\_manager module¶

*Module author: Sébastien Picault <>*

*class* `emulsion.agent.managers.group_manager.``GroupManager`(*state\_machine=None*, *\*\*others*)[source]¶
:   Bases: `emulsion.agent.views.StructuredView`

    An GroupManager is able to make its content
    evolve according to a specific state machine, the state of each
    subcompartment being stored in a specific state variable or
    attribute.

    `__init__`(*state\_machine=None*, *\*\*others*)[source]¶
    :   Create an GroupManager based on the
        specified state machine. The state of each subcompartment can
        be retrieved in the specified statevar name (‘true’ statevar
        or attribute)

    `agcount` *= 0*¶

    `agdict` *= {}*¶

    `apply_changes`(*transitions*, *productions*)[source]¶
    :   Apply modifications to the compartments contained in the current
        StructuredView, according to transitions and
        productions. Dictionary transitions is keyed by a tuple of
        variables and associated with a list of dictionaries, either
        {‘population’: qty, ‘actions’: list} or {‘agents’: list,
        ‘actions’: list}. List productions contains tuples (target,
        {‘population’: qty}, None) or (target, {‘agents’: list},
        prototype).

    `evolve`(*machine=None*)[source]¶
    :   Ask each unit in the current compartment to make its content evolve
        according to its own capabilities. A specific state machine
        can be specified if needed.

    `evolve_states`(*machine=None*)[source]¶
    :   Ask each compartment to make its content evolve according
        to its current state and the specified state\_machine.

    `families` *= SortedSet(['GroupManager'], key=None, load=1000)*¶

    `init_counts`(*index=0*)[source]¶
    :   Initialize the counts.

    `members` *= ('agcount', 'agdict', 'families', '\_\_module\_\_', '\_\_qualname\_\_', '\_\_doc\_\_', '\_\_init\_\_', 'init\_counts', 'update\_counts', 'apply\_changes', 'evolve', 'evolve\_states', '\_evolve\_transitions', '\_evolve\_productions', '\_\_classcell\_\_')*¶

    `update_counts`(*index=0*)[source]¶
    :   Update the number of atoms for each state of the state
        machine (TODO: for each value of the key[index] enum).

#### emulsion.agent.managers.ibm\_process\_manager module¶

*Module author: Sébastien Picault <>*

*class* `emulsion.agent.managers.ibm_process_manager.``IBMProcessManager`(*\*\*others*)[source]¶
:   Bases: `emulsion.agent.managers.multi_process_manager.MultiProcessManager`

    An IBMProcessManager is a MultiProcessManager dedicated to the
    management of Individual-Based Models. This class is endowed with
    a counters attribute which is a dictionary {process -> counter of
    states in relation with the process}.

    `__init__`(*\*\*others*)[source]¶
    :   Initialize the unit with a health state and a name.

    `add_atoms`(*atom\_set*, *init=False*, *\*\*others*)[source]¶
    :   Add the specified set of atoms to the current
        MultiProcessManager. Atoms are especially added
        automatically to each of the compartment managers. If init
        is True, the compartment managers counts the initial value of
        the populations in each compartment.

    `agcount` *= 0*¶

    `agdict` *= {}*¶

    `counts`¶
    :   Return a pandas DataFrame containing counts of each process if
        existing.

    `evolve`(*\*\*others*)[source]¶
    :   Make the agent evolve and update counts based on sub-level
        agents.

    `families` *= SortedSet(['IBMProcessManager'], key=None, load=1000)*¶

    `find_sublevel_statemachines`()[source]¶
    :   Retrieve state machines used as processes by agents from the
        sub-level.

    `get_sublevels`()[source]¶
    :   Return the list of sublevels contained in this level.

    `members` *= ('agcount', 'agdict', 'families', '\_\_module\_\_', '\_\_qualname\_\_', '\_\_doc\_\_', '\_\_init\_\_', 'add\_atoms', 'get\_sublevels', 'find\_sublevel\_statemachines', 'evolve', 'update\_counts', 'counts', 'remove\_randomly', '\_\_classcell\_\_')*¶

    `remove_randomly`(*proba=0*, *statevar=None*)[source]¶
    :   Remove randomly chosen atoms from this ProcessManager. proba can
        be either a probability or a dictionary. In that case, the
        statevar parameter indicates the name of the state variable
        which drives the probabilities, and the keys must be valid
        values for this state variable. Selected atoms are removed and
        returned by the method.

    `update_counts`()[source]¶
    :   Update counters based on invdividual status.

#### emulsion.agent.managers.metapop\_process\_manager module¶

*Module author: Sébastien Picault <>*

*class* `emulsion.agent.managers.metapop_process_manager.``MetapopProcessManager`(*model=None*, *level=None*, *\*\*others*)[source]¶
:   Bases: `emulsion.agent.managers.multi_process_manager.MultiProcessManager`

    This class is in charge of handling multiple populations.

    `agcount` *= 0*¶

    `agdict` *= {}*¶

    `counts`¶
    :   Return a pandas DataFrame containing counts of each process if
        existing.

    `families` *= SortedSet(['MetapopProcessManager'], key=None, load=1000)*¶

    `get_populations`()[source]¶

    `members` *= ('agcount', 'agdict', 'families', '\_\_module\_\_', '\_\_qualname\_\_', '\_\_doc\_\_', 'get\_populations', 'counts')*¶

#### emulsion.agent.managers.multi\_process\_manager module¶

*Module author: Sébastien Picault <>*

*class* `emulsion.agent.managers.multi_process_manager.``MultiProcessManager`(*model=None*, *level=None*, *\*\*others*)[source]¶
:   Bases: `emulsion.agent.managers.abstract_process_manager.AbstractProcessManager`

    A MultiProcessManager is aimed handling several independent
    StructuredViews at the same time, together with a
    SimpleView containing all the atom units. It can
    automatically build compartments for:
    - state machines associated with a specific state variable or attribute
    - specific state variables or attributes with a limited number of
    values, such as booleans or enumerations

    `__init__`(*model=None*, *level=None*, *\*\*others*)[source]¶
    :   Initialize the unit with a health state and a name.

    `add_atoms`(*atom\_set*, *init=False*, *level=None*)[source]¶
    :   Add the specified set of atoms to the current
        MultiProcessManager. Atoms are especially added
        automatically to each of the compartment managers. If init
        is True, the compartment managers counts the initial value of
        the populations in each compartment.

    `add_compart_process`(*process\_name*, *key\_variables*, *compart\_manager=(<class 'emulsion.agent.managers.group\_manager.GroupManager'>*, *{})*, *machine\_name=None*, *allowed\_values=None*, *compart\_class=(<class 'emulsion.agent.views.AdaptiveView'>*, *{})*)[source]¶
    :   Add a process aimed at managing a ‘Compartment Manager’, i.e. an
        object aimed at managing a collection of compartments. This
        compartment manager is automatically initialized from the
        compart\_manager class (which should be a subclass of
        StructuredView or GroupManager). The compartment manager may
        be associated with a specific state machine, and MUST BE
        identified by a tuple of state variables names. Additionally,
        since a first compartment is also instantiated, a specific
        class to do so can be also specified.

    `add_host`(*host*)[source]¶
    :   Add the specified host to the current Multiprocessmanager, associated
        with the specified key.

    `add_new_population`(*process\_name*, *population*)[source]¶
    :   Create new atoms with the population information

    `agcount` *= 0*¶

    `agdict` *= {}*¶

    `apply_initial_conditions`()[source]¶
    :   Initialize level with initial conditions specified in the model.

        As this agent is aimed at managing individuals in the
        sub-levels, only ‘prototypes’ are taken into account to
        initialize the sub-levels.

    `families` *= SortedSet(['MultiProcessManager'], key=None, load=1000)*¶

    `get_agent_class_for_sublevel`(*sublevel*)[source]¶
    :   Return the agent class in charge of representing the specified
        sublevel.

    `get_default_sublevel`()[source]¶
    :   Return by default the first sublevel contained in this level, if
        any. If this level contains no sublevels, raise a
        LevelException.

    `get_group_atoms`(*process\_name*, *group\_name*)[source]¶
    :   Return all atoms which belong to the the specified group
        name (state names). (*group\_name* may be a subset of the grouping key)

    `make_all_consistent`()[source]¶
    :   Check all compartments to ensure their consistency

    `make_consistent`(*compartment*)[source]¶
    :   Make the specified dict compartment check and handle the
        consistency of its own sub-compartments.

    `members` *= ('agcount', 'agdict', 'families', '\_\_module\_\_', '\_\_qualname\_\_', '\_\_doc\_\_', '\_\_init\_\_', 'apply\_initial\_conditions', 'add\_host', 'add\_new\_population', 'add\_compart\_process', 'select\_atoms', 'get\_group\_atoms', 'get\_agent\_class\_for\_sublevel', 'get\_default\_sublevel', 'new\_atom', 'add\_atoms', 'make\_all\_consistent', 'make\_consistent', 'remove', 'remove\_atoms', 'select\_randomly', 'remove\_randomly', '\_\_classcell\_\_')*¶

    `new_atom`(*sublevel=None*, *prototype=None*, *custom\_prototype=None*, *execute\_actions=False*, *\*\*args*)[source]¶
    :   Instantiate a new atom for the specified sublevel, with the
        specified arguments. If the sublevel is not specified, the
        first one from the contains list is taken. If the name of a
        prototype is provided, it is applied to the new agent (using
        the execute\_actions parameter).

    `remove`(*agents\_or\_population*)[source]¶
    :   Remove the specified population from the current
        compartment. The population is expected to be a dictionary
        with names as keys and compartments as values.

    `remove_atoms`(*atom\_set*)[source]¶
    :   Remove the specified atoms from the current
        MultiProcessManager. Atoms are removed from each of the
        compartment managers (including the ‘MASTER’ set).

    `remove_randomly`(*proba=0*, *amount=None*, *process=None*)[source]¶
    :   Remove randomly chosen atoms from this ProcessManager. proba can
        be either a probability or a dictionary. In that case, the
        process parameter indicates the name of the process grouping
        which drives the probabilities, and the keys must be those of
        the grouping. Selected atoms are removed and returned by the
        method.

    `select_atoms`(*variable=None*, *state=None*, *value=None*, *process=None*)[source]¶
    :   Return a list of atoms selected by specific *value* or *state* of a
        *variable*.

        |  |  |
        | --- | --- |
        | Parameters: | - **variable** (*str*) – the variable to be compared to the *value* - if `None`,   all atoms are selected. - **state** (*str*) – a name of the state used for the selection (instead of *value*) - **value** (*object*) – the value to use for selection - if `None`, the value is   replaced with the model state specified in parameter   *state* - **process** (*str*) – name of a process associated to a specific grouping based   on the *variable* (to accelerate search) |
        | Returns: | *list* – the list of matching agents in the sublevel |

    `select_randomly`(*proba=0*, *amount=None*, *process=None*)[source]¶
    :   Select randomly chosen atoms from this ProcessManager. proba can
        be either a probability or a dictionary. In that case, the
        process parameter indicates the name of the process grouping
        which drives the probabilities, and the keys must be those of
        the grouping. Selected atoms are removed and returned by the
        method.

### EMULSION

Epidemiological Multi-Level Simulation Framework

##### Navigation

- 1. Installation
- 2. Getting started with EMULSION
- 3. Modelling principles
- 4. Modelling language (basics)
- 5. Modelling language (advanced)
- 6. Feature examples
- 7. Information
- 8. License
- 9. High-level functions for model designers
- 10. emulsion package
  - 10.1. Subpackages
    - emulsion.agent package
    - emulsion.model package
    - emulsion.tools package
  - 10.2. Submodules
  - 10.3. emulsion.init\_emulsion module

##### Related Topics

- Documentation overview
  - 10. emulsion package
    - emulsion.agent package
      - Previous: emulsion.agent.core package
      - Next: emulsion.model package

##### Quick search

©2016, INRA and Univ. Lille.
|
Powered by Sphinx 1.8.5
& Alabaster 0.7.10
|
Page source
