## Supplementary material for "EMULSION: transparent and flexible multiscale stochastic models in human, animal and plant epidemiology": S2 File: emulsion.tools.html

emulsion.tools package — EMULSION (Epidemiological Multi-Level Simulation framework)

### emulsion.tools package¶

A Python implementation of the EMuLSion framework (Epidemiologic
MUlti-Level SImulatiONs).

Misc. tools

#### Submodules¶

#### emulsion.tools.calendar module¶

Classes and functions for the definition of Emulsion calendars.

*class* `emulsion.tools.calendar.``EventCalendar`(*calendar\_name*, *step\_duration*, *origin*, *period=None*, *\*\*initial\_dates*)[source]¶
:   Bases: `object`

    The EventCalendar class is intended to handle events and
    periods of time. Dates/times are given to the calendar in a
    human-readable format, and converted to integers (simulation
    steps).

    `__init__`(*calendar\_name*, *step\_duration*, *origin*, *period=None*, *\*\*initial\_dates*)[source]¶
    :   Initialize the calendar using the specified information.

        |  |  |
        | --- | --- |
        | Parameters: | - **calendar\_name** (*string*) – the name of the calendar - **step\_duration** (`datetime.timedelta`) – actual duration of one simulation step - **origin** (`datetime.datetime`) – date/time value of the beginning of the calendar - **period** (`datetime.timedelta`) – if the calendar is periodic, actual duration of the   period, None otherwise - **initial\_dates** (*dict*) – initial dictionary of events (event name as key, dict as   value with either begin/end dates or one single date if   punctual event) |

    `add_event`(*name*, *begin\_end*)[source]¶
    :   Add the specified event to the calendar. An event is characterized
        by its *name* and a *begin\_end* tuple indicating the begin and
        end dates.

        |  |  |
        | --- | --- |
        | Parameters: | - **name** (*str*) – the name of the event - **begin\_end** (*tuple*) – a tuple composed of the begin date and the end date   (possibly identical for handling punctual events) |

    `date_to_step`(*date*)[source]¶
    :   Return the step corresponding the specified date.

        |  |  |
        | --- | --- |
        | Parameters: | **date** (`datetime.datetime`) – the date to convert into time steps |
        | Returns: | **t** (*int*) – the time step during which the specified date occurs (the date occurs between the returned time step *t* and time step *t+1*) |

    `get_events`()[source]¶
    :   Return the list of events contained in the current calendar.

        |  |  |
        | --- | --- |
        | Returns: | **l** (*list*) – the list of all event names |

    `increment`(*steps=1*)[source]¶
    :   Advance the current date by the specified number of
        simulation steps.

        |  |  |
        | --- | --- |
        | Parameters: | **steps** (*int*) – number of steps to ‘add’ to the current date |

    `step_to_date`(*step*)[source]¶
    :   Return the date when the specified step begins.

        |  |  |
        | --- | --- |
        | Parameters: | **step** (*int*) – the time step to convert to a date |
        | Returns: | **d** (`datetime.datetime`) – the corresponding date |

*exception* `emulsion.tools.calendar.``InvalidIntervalException`(*begin*, *end*)[source]¶
:   Bases: `Exception`

    Exception raised when trying to insert an inconsistent event in the
    calendar. An event is considered inconsistent if the begin date is
    posterior to the send date in a non-periodic calendar.

    `__init__`(*begin*, *end*)[source]¶
    :   Create the exception with the incorrect *begin* and *end* dates.

`emulsion.tools.calendar.``date_in`(*begin*, *end*, *period=None*)[source]¶
:   Build a function which tests if a date belongs to the specified
    interval (from *begin* to *end*). If a *period* is specified, the
    test relies upon the periodicity, otherwise dates are considered
    absolute.

    |  |  |
    | --- | --- |
    | Parameters: | - **begin** (`datetime.datetime`) – the date when the interval begins - **end** (`datetime.datetime`) – the date when the interval ends - **period** (`datetime.timedelta`) – the duration of the period after which the calendar cycles |
    | Returns: | *lambda* – a function which maps a date (`datetime.datetime`) to a `bool` to indicate whether or not the date belongs to the *begin*-*end* interval. |
    | Raises: | `InvalidIntervalexception` – if *begin* is posterior to *end* in a non-periodic calendar |

#### emulsion.tools.functions module¶

Additional functions for symbolic computing in YAML model definition files.

All functions in this module can be used in EMULSION models.

`emulsion.tools.functions.``AND`(*\*values*)[source]¶
:   Return a logical AND (conjunction) between all the values.

    Example

    To define symptomatic individuals as infected for at least 5 days:

    ```
    is_symptomatic:
      desc: 'test if individuals has symptoms'
      value: 'AND(is_I, duration_in_health_state >= 5)'
    ```

    |  |  |
    | --- | --- |
    | Parameters: | **\*values** (*list*) – boolean values separated by commas |
    | Returns: | *bool* – True if **all** values are True, False otherwise. |

`emulsion.tools.functions.``IfThenElse`(*condition*, *val\_if\_true*, *val\_if\_false*)[source]¶
:   Ternary conditional function: return either *val\_if\_true* or
    *val\_if\_false* depending on *condition*.

    Example

    Here in a parameter definition (assuming that `summer_period` is
    defined e.g. in the *calendars* section of the model):

    ```
    average_temperature:
      desc: 'average temperature for the season'
      value: 'IfThenElse(summer_period, 25, 8)'
    ```

    |  |  |
    | --- | --- |
    | Parameters: | - **condition** (*bool*) – a boolean expression - **val\_if\_true** (*number*) – value to return if the expression is True - **val\_if\_false** (*number*) – value to return if the expression is False |
    | Returns: | *number* – One of the two values, depending on *condition*. |

`emulsion.tools.functions.``OR`(*\*values*)[source]¶
:   Return a logical OR (disjunction) between all the values.

    Example

    To sell animals depending on either their weight or their age:

    ```
    transitions:
      ...
      - from: Fattening
        to: Sold
        proba: 1
        cond: 'OR(weight >= weight_thr, age >= age_thr)'
    ```

    |  |  |
    | --- | --- |
    | Parameters: | **\*values** (*list*) – boolean values separated by commas |
    | Returns: | *bool* – True if **at least one** of the values is True, False otherwise. |

`emulsion.tools.functions.``random_beta`(*a: float*, *b: float*) → float[source]¶
:   Return a random value drawn from a beta distribution of parameters
    *a* and *b*.

    Example

    In a prototype definition:

    ```
    age: 'random_beta(2, 5) * age_max'
    ```

    |  |  |
    | --- | --- |
    | Parameters: | - **a** (*float**,* *positive* *(**>0**)*) – first shape parameter of the beta distribution - **b** (*float**,* *positive* *(**>0**)*) – second shape parameter of the beta distribution |
    | Returns: | *float* – a random value sampled according to a beta distribution of parameters *a* and *b* |

    See also

    `numpy.random.beta()`

`emulsion.tools.functions.``random_bool`(*proba\_success: float*) → int[source]¶
:   Return a random boolean value (actually, 0 or 1) depending on
    *proba\_success*.

    Example

    Set the value of a state variable `has_symptoms` when entering
    in the infectious state:

    ```
    states:
      ...
      - I:
        ...
        on_enter:
          - set_var: has_symptoms
            value: 'random_bool(proba_symptomatic)'
    ```

    |  |  |
    | --- | --- |
    | Parameters: | **proba\_success** (*float in* *[**0**,**1**]*) – probability of returning 1 (True) |
    | Returns: | *int* – either 1 with probability *proba\_success*, or 0 with probability 1-*proba\_success* |

`emulsion.tools.functions.``random_choice`(*\*values*)[source]¶
:   Return a value chosen randomly among those provided (equiprobable
    sampling).

    Example

    To init `age` among three typical values:

    ```
    prototypes:
      individuals:
        init_individual:
          age: 'random_choice(10, 50, 200)'
    ```

    |  |  |
    | --- | --- |
    | Parameters: | **\*values** (*list*) – the possible values, separated by commas |
    | Returns: | *val* – one of the values (equiprobable choice) |

`emulsion.tools.functions.``random_choice_weighted`(*\*values*)[source]¶
:   Return a value chosen randomly among those provided (but not
    equiprobably).

    Example

    To init `age` among three typical values which respectively
    represent 10%, 20% and 70% of the population:

    ```
    prototypes:
      individuals:
        init_individual:
          age: 'random_choice_weighted(10, 0.1, 50, 0.2, 200, 0.7)'
    ```

    |  |  |
    | --- | --- |
    | Parameters: | **values** (*list*) – possibles choices and their weight, alternatively, separated by commas. Weights are normalized to be used as probabilities. |
    | Returns: | *One of the choices* |

`emulsion.tools.functions.``random_exponential`(*scale: float*) → float[source]¶
:   Return a random value drawn from an exponential distribution of
    rate 1/*scale* (thus of mean *scale*).

    Example

    In a prototype definition:

    ```
    time_to_live: random_exponential(mean_duration)
    ```

    |  |  |
    | --- | --- |
    | Parameters: | **scale** (*float*) – the scale parameter of the distribution, i.e. the inverse of the rate |
    | Returns: | *float* – a random value sampled according to an exponential distribution of rate 1/*scale* |

    See also

    `numpy.random.exponential()`

`emulsion.tools.functions.``random_gamma`(*shape: float*, *scale: float*) → float[source]¶
:   Return a random value drawn from a gamma distribution of parameters
    *shape* and *scale*.

    Example

    In a prototype definition:

    ```
    age: 'random_gamma(3, 2)'
    ```

    |  |  |
    | --- | --- |
    | Parameters: | - **shape** (*float**,* *positive* *(**>0**)*) – the shape of the gamma distribution - **scale** (*float**,* *positive* *(**>0**)*) – the scale of the gamma distribution |
    | Returns: | *float* – a random value sampled according to a gamma distribution of parameters *shape* and *scale* |

    See also

    `numpy.random.gamma()`

`emulsion.tools.functions.``random_integers`(*low: int*, *high: int*) → int[source]¶
:   Return a random integer value drawn from a discrete uniform
    distribution between *low* and *high* (both inclusive).

    Example

    In a prototype definition:

    ```
    age: random_integers(min_age, max_age)
    ```

    |  |  |
    | --- | --- |
    | Parameters: | - **low** (*int*) – lower boundary of the sample interval (inclusive) - **high** (*int*) – upper boundary of the sample interval (inclusive) |
    | Returns: | *int* – a random integer value sampled according to a discrete uniform distribution between *low* and *high* |

    See also

    `numpy.random.random_integers()`

`emulsion.tools.functions.``random_multinomial`(*number\_of\_samples*, *\*probas*)[source]¶
:   Return a multinomial sample based on the specified probabilities.

    |  |  |
    | --- | --- |
    | Parameters: | - **number\_of\_samples** (*int*) – number of experiments - **probas** (*list*) – list of probabilities |
    | Returns: | *list* – the drawn samples |

`emulsion.tools.functions.``random_normal`(*mn: float*, *sd: float*) → float[source]¶
:   Return a random value drawn from a normal distribution of mean *mn*
    and standard deviation *sd*.

    Example

    In a prototype definition:

    ```
    age: 'random_normal(100, 5)'
    ```

    |  |  |
    | --- | --- |
    | Parameters: | - **mn** (*float*) – the mean of the normal distribution - **sd** (*float**,* *positive* *(**>=0**)*) – the standard deviation of the normal distribution |
    | Returns: | *float* – a random value sampled according to a normal distribution of mean *mn* and standard deviation *sd* |

    See also

    `numpy.random.normal()`

`emulsion.tools.functions.``random_poisson`(*lam: float*) → int[source]¶
:   Return an integer random value drawn from a Poisson distribution of
    mean *lam*.

    Example

    In an action, e.g. here when computing how many newborn
    individuals will be produced:

    ```
    - from: Gestating
      to: NonGestating
      on_cross:
        - produce_offspring: newborn
          amount: 'random_poisson(1.05)'
    ```

    |  |  |
    | --- | --- |
    | Parameters: | **lam** (*float*) – the mean of the Poisson distribution |
    | Returns: | *int* – a random integer value sampled according to a Poisson distribution of mean *lam* |

    See also

    `numpy.random.poisson()`

`emulsion.tools.functions.``random_uniform`(*low: float*, *high: float*) → float[source]¶
:   Return a random value drawn from a uniform distribution between
    *low* (inclusive) and *high* (exclusive).

    Example

    In a prototype definition:

    ```
    age: random_uniform(min_age, max_age)
    ```

    |  |  |
    | --- | --- |
    | Parameters: | - **low** (*float*) – lower boundary of the sample interval (inclusive) - **high** (*float*) – upper boundary of the sample interval (exclusive) |
    | Returns: | *float* – a random value sampled according to a uniform distribution between *low* and *high* |

    See also

    `numpy.random.uniform()`

#### emulsion.tools.graph module¶

A Python implementation of the EMuLSion framework (Epidemiologic
MUlti-Level SImulatiONs).

Tools aimed at handling graphs for the state machines

*class* `emulsion.tools.graph.``EdgeTypes`¶
:   Bases: `emulsion.tools.state.EmulsionEnum`

    An enumeration.

    `PRODUCTION` *= 2*¶

    `TRANSITION` *= 1*¶

    `linestyle` *= 'solid'*¶

*class* `emulsion.tools.graph.``MultiDiGraph`(*\*\*attributes*)[source]¶
:   Bases: `object`

    An oriented multigraph (two nodes can be linked by several
    edges). Replacement for MultiDiGraph in networkx with a
    reproducible order in accessing edges and nodes.

    `__init__`(*\*\*attributes*)[source]¶
    :   Create an instance of MultiDiGraph. If keywords arguments
        are specified, they are considered attributes of the whole
        graph.

    `add_edge`(*from\_id*, *to\_id*, *type\_id=<EdgeTypes.TRANSITION>*, *key=None*, *\*\*attributes*)[source]¶
    :   Add the specified edge to the graph. If nodes are not
        already created, they are automatically added. Edge attributes
        can be specified. Since this graph allows multiple edges
        between pairs of nodes, two calls to this method lead to two
        distinct edges (even if the attributes are the same), unless a
        key is specified. The key is used to identify edges between a
        given pair of nodes. By default, keys are consecutive
        integers.

    `add_node`(*node\_id*, *\*\*attributes*)[source]¶
    :   Add the specified node to the graph. Node attributes can be
        specified. If the node is already present, updated the
        attributes.

    `edges`()[source]¶
    :   Return a list of edge tuples.

    `edges_from`(*from\_id*, *type\_id=<EdgeTypes.TRANSITION>*)[source]¶
    :   Return a list of tuples (to\_id, attributes) corresponding
        to all edges going out of the from\_id node.

#### emulsion.tools.misc module¶

Collection of various useful functions used in EMULSION framework,
especially regarding introspection.

`emulsion.tools.misc.``AGENTS` *= 1*¶
:   Constant value for the index where list of agents are stored in
    ‘populations’ tuples.

`emulsion.tools.misc.``POPULATION` *= 1*¶
:   Constant value for the index where population amounts are stored in
    ‘populations’ tuples.

`emulsion.tools.misc.``add_all_test_properties`(*agent*)[source]¶

`emulsion.tools.misc.``add_new_property`(*agent*, *property\_name*, *getter\_function*)[source]¶
:   Add a new property to an agent.

    Actually, the property is added to the class of the agent, as
    Python properties are descriptors. Yet, the dynamic attribution of
    properties must me done through instances rather than classes,
    since agents must add the name of the property to their
    `_mbr_cache` attribute.

    |  |  |
    | --- | --- |
    | Parameters: | - **agent** (*AbstractAgent*) – the agent to which the property must be added - **property\_name** (*str*) – the name of the property - **getter\_function** (*lambda*) – the function upon which the property is built |

`emulsion.tools.misc.``aggregate_probabilities`(*probability\_values: Iterable[float], delta\_t: float*) → Iterable[float][source]¶
:   From the specified *probability\_values*, intended to represent
    probabilities of events duting one time unit, compute the
    probabilities for the specified time step (*delta\_t*).

    |  |  |
    | --- | --- |
    | Parameters: | - **probability\_values** (*list*) – a list of probability values for several events (during 1 time unit) - **delta\_t** (*float*) – the value of the time step (expressed in time units) |
    | Returns: | *list* – the probabilities of the same events during *delta\_t* time units. |

`emulsion.tools.misc.``aggregate_probability`(*probability: float*, *delta\_t: float*) → float[source]¶
:   Transform the specified *probability* value, intended to represent
    a probability for events tested each time unit, into the
    probability for the specified time step (*delta\_t*).

    |  |  |
    | --- | --- |
    | Parameters: | - **probability** (*float*) – the probability value of an event (during 1 time unit) - **delta\_t** (*float*) – the value of the time step (expressed in time units) |
    | Returns: | *float* – the probability of the event during *delta\_t* time units. |

`emulsion.tools.misc.``count_population`()[source]¶
:   Return the amount of atoms represented in *agents\_or\_pop*.

    |  |  |
    | --- | --- |
    | Parameters: | **agents\_or\_pop** (*tuple*) – either (‘population’, qty) or (‘agents’, list of agents) |
    | Returns: | *int or float* – the amount corresponding to the population: generally, an int value, but deterministic models produce float values. |

`emulsion.tools.misc.``create_aggregator`(*sourcevar: str*, *operator: str*)[source]¶
:   Create an aggregator function to be used as property getter. This
    aggregator function has to collect all values of *sourcevar* for
    agents contained in a given host, and reduce them to one avalue
    using the specified *operator*.

    |  |  |
    | --- | --- |
    | Parameters: | - **sourcevar** (*str*) – the name of the variable to collect in the sublevel - **operator** (*str*) – the name of the operator to apply to the collected values |
    | Returns: | *lambda* – A function which can be applied to a MultiProcessManager agent (i.e. with explicit sublevel agents) which returns the aggregated values for the whole population. |

`emulsion.tools.misc.``create_atoms_aggregator`(*sourcevar: str*, *operator: str*, *machine\_name: str*, *state\_name: str*)[source]¶
:   Build a getter for property of the form `newvar_X` where `X` if
    the value of *state\_name* (a state of *state\_machine*), `newvar`
    is an aggregate variable based on collecting all values of
    *sourcevar* for the specific *group\_name* and aggregating the
    values using *operator*. This function is intended to work on
    IBMProcessManager agents, which do not benefit from groupings.

    |  |  |
    | --- | --- |
    | Parameters: | - **sourcevar** (*str*) – the name of the variable to collect in the sublevel - **operator** (*str*) – the name of the operator to apply to the collected values - **machine\_name** (*str*) – the name of the state machine for which the getter is created - **state\_name** (*str*) – the state name |
    | Returns: | *lambda* – A function which can be applied to an IBMProcessManager agent (i.e. with explicit but ungrouped sublevel agents) which returns the aggregated value for the specified group. |

`emulsion.tools.misc.``create_counter_getter`(*machine\_name*, *state\_name*)[source]¶

`emulsion.tools.misc.``create_duration_getter`(*machine\_name*)[source]¶

`emulsion.tools.misc.``create_group_aggregator`()[source]¶
:   Build a getter for property of the form `newvar_X_Y` where `X`
    and `Y` are states of two different states machines used in a
    grouping, `newvar` is an aggregate variable based on collecting
    all values of *sourcevar* for the specific *group\_name* and
    aggregating the values using *operator*. The functions can handle
    groupings with an arbitrary number of states.

    |  |  |
    | --- | --- |
    | Parameters: | - **sourcevar** (*str*) – the name of the variable to collect in the sublevel - **operator** (*str*) – the name of the operator to apply to the collected values - **process\_name** (*str*) – the name of the process associated with the grouping for which   the population getter is created - **group\_name** (*str* *or* *tuple*) – either a string representing the state name (if only one), or   a tuple of several state names |
    | Returns: | *lambda* – A function which can be applied to an MultiProcessManager agent (i.e. with explicit sublevel agents) which returns the aggregated value for the specified group. |

`emulsion.tools.misc.``create_new_serial`(*end=None*, *model=None*)[source]¶
:   Create the serial number generator associated to the specified
    variable.

`emulsion.tools.misc.``create_population_getter`()[source]¶
:   Build a getter for property of the form `total_X_Y` where `X`
    and `Y` are states of two different states machines used in a
    grouping.

    The functions can handle groupings with an arbitrary number of states.

    |  |  |
    | --- | --- |
    | Parameters: | - **process\_name** (*str*) – the name of the process associated with the grouping for which   the population getter is created - **group\_name** (*str* *or* *tuple*) – either a string representing the state name (if only one), or   a tuple of several state names |
    | Returns: | *callable* – A function which can be applied to an AbstractProcessManager agent which returns the population size for the specified group. |

`emulsion.tools.misc.``create_state_tester`(*state\_name*)[source]¶

`emulsion.tools.misc.``create_weighted_random`(*machine\_name*, *weights*, *model=None*)[source]¶
:   Create a random choice function which returns a random state from
    the given state machine (among non-autoremove states), according
    to the weights. Weights are interpreted either directly as
    probabilities (if the number of weights is stricly one below the
    number of available states, the last state getting the complement
    to 1), or as true weights which are then normalized to be used as
    probabilities.

    machine\_name: str
    :   the name of the state machine where the states must be chosen
        among the *N* non-autoremove states.

    weights: list
    :   a list of *N* or *N-1* model expressions assumed to produce positive numbers

    model: EmulsionModel
    :   the model where this function is defined

    |  |  |
    | --- | --- |
    | Returns: | **lambda** (*a function that returns a random state according to the*) – values of the weights list, interpreted either as probabilities (if size *N-1*) or as weights (if size *N*) which are then normalized to provide probabilities |

`emulsion.tools.misc.``find_operator`(*operator: str*)[source]¶
:   Return an aggregation function named *operator*.

    Search starts with emulsion functions (module
    emulsion.tools.functions), which includes Python built-ins, then
    in numpy.

    A special shortcut is provided for percentiles: `percentileXX`
    is interpreted as the partial function
    `numpy.percentile(q=int(XX))`.

    |  |  |
    | --- | --- |
    | Parameters: | **operator** (*str*) – the name of the aggregation operator |
    | Returns: | *lambda* – a function that takes a list (or array-like) as input and returns the application of the *operator* to the values. |

`emulsion.tools.misc.``load_class`(*module=None*, *class\_name: str = None*, *options: dict = {}*)[source]¶
:   Dynamically load the class with the specified *class\_name* from the
    given *module*.

    |  |  |
    | --- | --- |
    | Parameters: | - **module** – a Python module, where the class is expected to be located. - **class\_name** (*str*) – the name of the class to load - **options** – some options |
    | Returns: | *tuple* – a tuple composed of the class (type) and the options. |

    Todo

    - clarify the role of *options*

`emulsion.tools.misc.``load_module`(*module\_name: str*)[source]¶
:   Dynamically load the module with the specified *module\_name* and
    return it.

    |  |  |
    | --- | --- |
    | Parameters: | **module\_name** (*str*) – the name of a valid Python module (accessible in the `PYTHONPATH` environment variable). |
    | Returns: | *ref* – A reference to the Python module. |

`emulsion.tools.misc.``moving_average`(*values*, *window\_size*, *mode='same'*)[source]¶
:   Compute a moving average of the specified *values* with respect to
    the *window\_size* on which the average is calculated. The return
    moving average has the same size as the original values. To avoid
    boundary effects, use `mode='valid'`, which produce a result of
    size `len(values) - window_size + 1`.

    |  |  |
    | --- | --- |
    | Parameters: | - **values** (*array-like*) – contains the values for moving average computation - **window\_size** (*int*) – width of the moving average window - **mode** (*str*) – a parameter for numpy.convolve |
    | Returns: | *nd\_array* – a numpy `nd_array` containing the values of the moving average. |

`emulsion.tools.misc.``probabilities_to_rates`(*probability\_values: Iterable[float]*) → List[float][source]¶
:   Transform a list of probabilities into a list of rates. The
    last value is expected to represent the probability of staying in
    the current state.

    |  |  |
    | --- | --- |
    | Parameters: | **probability\_values** (*list*) – a list of probabilities, the last one representing the probability to stay in the current state |
    | Returns: | *list* – a list of rates corresponding to those probabilities. |

`emulsion.tools.misc.``rates_to_probabilities`(*total\_rate: float, rate\_values: List[float], delta\_t: float = 1*) → List[float][source]¶
:   Transform the specified list of *rate\_values*, interpreted as
    outgoing rates, into probabilities, according to the specified
    time step (*delta\_t*) and normalized by *total\_rate*.

    For exit rates \(\rho\_i\) (one of the *rate\_values*), the
    probability to stay in current state is given by:

    \[p\_0 = e^{-\delta t.\sum\_i \rho\_i}\]

    Thus, each rate \(\rho\_i\) corresponds to a probability

    \[p\_i = \frac{\rho\_i}{\sum\_i \rho\_i} (1 - p\_0)\]

    |  |  |
    | --- | --- |
    | Parameters: | - **total\_rate** (*float*) – the total exit rate, used for normalization purposes. If   *rate\_values* represent all possible exit rates, *total\_rate*   is their sum. - **rate\_values** (*list*) – the list of rates to transform - **delta\_t** (*float*) – the value of the time step (expressed in time units) |
    | Returns: | *list* – the list of probabilities corresponding to the *rate\_values*. |

`emulsion.tools.misc.``read_from_file`(*filename: str*)[source]¶
:   Read the specified YAML *filename* and return the corresponding
    Python document.

    |  |  |
    | --- | --- |
    | Parameters: | **filename** (*str*) – the name of the YAML file to load |
    | Returns: | *object* – a Python object built by parsing of the YAML file, i.e. either a dict, list, or even str/int/etc… (Most YAML document will produce dictionaries.) |

`emulsion.tools.misc.``retrieve_value`(*value\_or\_function*, *agent*)[source]¶
:   Return a value either directly given by parameter
    *value\_or\_function* if it is a true value, or computed from this
    parameter seen as a function, with the specified *agent* as argument.

    |  |  |
    | --- | --- |
    | Parameters: | - **value\_or\_function** – either a callable (function that applies to an agent to   retrieve an individual value), or the value itself - **agent** – the agent to use as parameter of the callable if necessary |
    | Returns: | *value* – the expected value (agent-based or not). |

`emulsion.tools.misc.``rewrite_keys`(*name*, *position*, *change\_list*)[source]¶

`emulsion.tools.misc.``select_random`(*origin: Iterable*, *quantity: int*, *exclude: sortedcontainers.sortedset.SortedSet = SortedSet([]*, *key=None*, *load=1000)*) → List[source]¶
:   Return a random selection of *quantity* agents from the *origin*
    group, avoiding those explicitly in the *exclude* set. If the
    *origin* population proves too small, all available agents are
    taken, irrespective to the *quantity*.

    |  |  |
    | --- | --- |
    | Parameters: | - **origin** (*iterable*) – the population where agents must be selected - **quantity** (*int*) – the number of agents to select in the population - **exclude** (*set*) – agents which are not available for the selection |
    | Returns: | *list* – a list of randomly selected agents according to the above constraints. |

`emulsion.tools.misc.``serial`(*start=0*, *end=None*, *model=None*)[source]¶
:   A very simple serial number generator.

#### emulsion.tools.parallel module¶

A Python implementation of the EMuLSion framework (Epidemiologic
MUlti-Level SImulatiONs).

Tools for parallel computing.

`emulsion.tools.parallel.``job`(*target\_simulation\_class*, *proc*, *\*\*others*)[source]¶
:   Simple job for a simple processes

`emulsion.tools.parallel.``job_dist`(*total\_task*, *workers*)[source]¶
:   Return a distribution of each worker need to do.

`emulsion.tools.parallel.``parallel_multi`(*target\_simulation\_class=<class 'emulsion.tools.simulation.MultiSimulation'>*, *nb\_simu=None*, *nb\_proc=1*, *\*\*others*)[source]¶
:   Parallel loop for distributing tasks in different processes

`emulsion.tools.parallel.``parallel_sensi`(*target\_simulation\_class=<class 'emulsion.tools.simulation.SensitivitySimulation'>*, *nb\_proc=1*, *\*\*others*)[source]¶
:   Parallel loop for distributing sensitivity tasks in different processes

#### emulsion.tools.plot module¶

A Python implementation of the EMuLSion framework (Epidemiologic
MUlti-Level SImulatiONs).

Plotting tools… to be improved!

`emulsion.tools.plot.``build_machine_plot`(*machine\_name*, *model*, *params*)[source]¶
:   Return a bokeh figure based on the representation of the state
    machine, with the associated colors.

    |  |  |
    | --- | --- |
    | Parameters: | **machine** – the state machine to which the states are related |
    | Returns: | A bokeh gridplot. |

`emulsion.tools.plot.``build_state_plot`(*counts*, *cols*, *machine*, *model*, *y='quantity'*, *group='state'*, *ylab='Number of individuals'*)[source]¶
:   Return a bokeh figure based on the representation of the states in
    the *counts* dataframe, with the associated colors.

    |  |  |
    | --- | --- |
    | Parameters: | - **counts** – a Pandas dataframe containing the values to plot - **cols** (*dict*) – dictionary which maps state/variable names to colors - **machine** – the state machine to which the states are related - **model** – the Emulsion model related to this plot - **y** (*str*) – the name of the field containing y values in the dataframe - **group** (*str*) – either ‘state’ (default) or ‘variables’, name of   the field containing each legend items. |
    | Returns: | A bokeh gridplot (reduced to one figure a the herd level, one figure per herd at metapopulation level). |

`emulsion.tools.plot.``plot_outputs`(*params*)[source]¶
:   Read outputs from previous runs and plot the corresponding
    figures. In the *params* dictionary, output\_dir is expected to
    contain a counts.csv file; figure\_dir is where the plot is
    saved.

#### emulsion.tools.simulation module¶

Tools for providing generic simulation classes.

*class* `emulsion.tools.simulation.``AbstractSimulation`(*start\_id: int = 0*, *model=None*, *model\_path: str = ''*, *stock\_agent: bool = True*, *output\_dir: str = 'outputs/'*, *target\_agent\_class=None*, *save\_results: bool = True*, *input\_dir: str = None*, *load\_from\_file: str = None*, *save\_to\_file: str = None*, *\*\*\_*)[source]¶
:   Bases: `object`

    Abstract class from which any simulation class inherits.

    `__init__`(*start\_id: int = 0*, *model=None*, *model\_path: str = ''*, *stock\_agent: bool = True*, *output\_dir: str = 'outputs/'*, *target\_agent\_class=None*, *save\_results: bool = True*, *input\_dir: str = None*, *load\_from\_file: str = None*, *save\_to\_file: str = None*, *\*\*\_*)[source]¶
    :   Initialize the simulation.

        |  |  |
        | --- | --- |
        | Parameters: | - **start\_id** – ID of the (first) simulation - **model** – instance of the model to run - **model\_path** – path to the filename holding the description of the   model, used if *model* is None - **stock\_agent** – TODO - **output\_dir** – name of the directory for simulation outputs - **target\_agent\_class** – agent class representing the top level in the   simulation - **save\_results** – True if simulation outputs have to be saved,   False otherwise. TODO: should be removed, and replaced   by a set of OutputManagers dedicated to the specific   expected outputs. - **load\_from\_file** – a filename from which the initial state of   the simulation (agents corresponding to levels with   their state) is read (instead of running the   initialize\_level method) - **save\_to\_file** – a filename in which the final state of the   simulation (agents corresponding to levels with their   state) is written (after running the finalize\_level   method) |

    `evolve`(*steps: int = 1*)[source]¶
    :   Operations to perform at each time step. Should be defined in
        subclasses.

        |  |  |
        | --- | --- |
        | Parameters: | **steps** – |

    `run`()[source]¶
    :   Entry point to simulation execution. Should be defined in
        subclasses.

    `update_csv_counts`(*df*, *dparams: dict = {}*)[source]¶
    :   Update the CSV recording of populations in each state.

*class* `emulsion.tools.simulation.``MultiSimulation`(*multi\_id: int = 0*, *nb\_simu: int = 100*, *set\_seed: bool = False*, *silent: bool = False*, *quiet: bool = False*, *dparams: dict = {}*, *\*\*others*)[source]¶
:   Bases: `emulsion.tools.simulation.AbstractSimulation`

    MultiSimulation can handle multiple repetitions of a given model.
    For sensibility study (same model with different values of variables),
    please check out SensitivitySimulation.

    `__init__`(*multi\_id: int = 0*, *nb\_simu: int = 100*, *set\_seed: bool = False*, *silent: bool = False*, *quiet: bool = False*, *dparams: dict = {}*, *\*\*others*)[source]¶
    :   Initialize a simulation with multiple repetitions of the same
        model.

    `counts`¶

    `evolve`(*steps=1*)[source]¶
    :   Operations to perform at each time step. Should be defined in
        subclasses.

        |  |  |
        | --- | --- |
        | Parameters: | **steps** – |

    `run`(*update=True*)[source]¶
    :   Run all repetitions one by one.

    `write_dot`()[source]¶

*class* `emulsion.tools.simulation.``OutputManager`(*model=None*, *output\_dir=''*, *output\_file='counts.csv'*, *log\_file='log.txt'*)[source]¶
:   Bases: `object`

    Manager to handle different outputs (csv, database,… etc)

    `__init__`(*model=None*, *output\_dir=''*, *output\_file='counts.csv'*, *log\_file='log.txt'*)[source]¶
    :   Initialize the output manager, specifying the model and the
        directory where outputs will be stored.

    `update_output_information`()[source]¶
    :   Update csv file path or database connection engine

    `update_output_type`()[source]¶
    :   Update output type if specified in model

    `update_outputs`(*df=None*)[source]¶
    :   Update outputs: writing in csv file or in database

*class* `emulsion.tools.simulation.``SensitivitySimulation`(*scenario\_path=None*, *df=None*, *nb\_multi=None*, *\*\*others*)[source]¶
:   Bases: `emulsion.tools.simulation.AbstractSimulation`

    SensitivitySimulation can handle sensibility study with a given
    pandas DataFrame of parameters or a path linked with file which contains
    scenarios of parameters. Then it will be transformed to a dictionary of
    scenario in the `` `d_scenario` `` attribute.

    For instance, d\_scenario could be the form (QFever model example) :
    :   {0: {‘m’: 0.7, ‘q’: 0.02 …},
        :   1: {‘m’: 0.5, ‘q’: 0.02 …},
            2: …,
            … }

    `__init__`(*scenario\_path=None*, *df=None*, *nb\_multi=None*, *\*\*others*)[source]¶
    :   Initialize the simulation.

        |  |  |
        | --- | --- |
        | Parameters: | - **start\_id** – ID of the (first) simulation - **model** – instance of the model to run - **model\_path** – path to the filename holding the description of the   model, used if *model* is None - **stock\_agent** – TODO - **output\_dir** – name of the directory for simulation outputs - **target\_agent\_class** – agent class representing the top level in the   simulation - **save\_results** – True if simulation outputs have to be saved,   False otherwise. TODO: should be removed, and replaced   by a set of OutputManagers dedicated to the specific   expected outputs. - **load\_from\_file** – a filename from which the initial state of   the simulation (agents corresponding to levels with   their state) is read (instead of running the   initialize\_level method) - **save\_to\_file** – a filename in which the final state of the   simulation (agents corresponding to levels with their   state) is written (after running the finalize\_level   method) |

    `counts`¶

    `run`()[source]¶
    :   Make the simulation advance.

    `write_dot`()[source]¶

*class* `emulsion.tools.simulation.``Simulation`(*steps: int = 100*, *simu\_id: int = 0*, *silent: bool = False*, *quiet: bool = False*, *\*\*others*)[source]¶
:   Bases: `emulsion.tools.simulation.AbstractSimulation`

    Simulation class is aimed at running one repetition of a given
    model (for several repetitions, use MultiSimulation).

    `__init__`(*steps: int = 100*, *simu\_id: int = 0*, *silent: bool = False*, *quiet: bool = False*, *\*\*others*)[source]¶
    :   Create an instance of simulation.

        |  |  |
        | --- | --- |
        | Parameters: | - **steps** – number of time steps to run - **simu\_id** – ID of the simulation - **silent** – if False, show a progress bar during simulation execution - **quiet** – if True, show no progressbar at all |

        See also

        `emulsion.tools.simulation.AbstractSimulation`\_

    `counts`¶
    :   Return a pandas DataFrame contains counts of each process if existing.
        TODO: column steps need to be with one of process
        and NO column steps for inter herd

    `evolve`(*steps: int = 1*)[source]¶
    :   Make the target agent evolve.

        |  |  |
        | --- | --- |
        | Parameters: | **steps** – the number of time steps to run |

    `init_agent`(*\*\*others*)[source]¶
    :   Create an agent from the target class.

    `log_path`()[source]¶
    :   Return the log path used by current simulation

    `run`(*dparams: dict = {}*)[source]¶
    :   Make the simulation progress.

#### emulsion.tools.state module¶

This module defines:

- a class `StateVarDict` which is a special auto-referential
  dictionary to handle state variables
- a class `EmulsionEnum` which is a special kind of enumeration
  intended to handle states from state machines

*class* `emulsion.tools.state.``EmulsionEnum`[source]¶
:   Bases: `enum.Enum`

    This class represents enumerations for states of state machines in
    EMULSION. They are endowed with some special features:

    1. They provide total ordering between items (based on `__lt__`
       and `__eq__` methods).
    2. A comparison with `None` is provided (any state is always
       greater than `None`).
    3. Other features will be developed soon.

*class* `emulsion.tools.state.``StateVarDict`(*\*args*, *\*\*kwargs*)[source]¶
:   Bases: `dict`

    A special dictionary aimed at handling the state variables of
    agents in EMULSION models. In addition to the classical dict
    key-based access, it provides an attribute-like access syntax.

    This class is used in EMULSION to store agent properties. Such
    properties include those defined automatically and used by the
    EMULSION engine, such as the values of the states for each state
    machine, the current time step, or “hidden” values such as the
    time spent in the current state for each state machine. They are
    also used to include user-defined attributes (e.g. age,
    weight…).

    When searching for a model `statevar`, the engine tries first to
    find a classical instance variable (which can be mimicked by a
    Python `@property`-decored function), then looks inside the
    agent’s `statevar` attribute; finally, the search continues in
    the agent’s host (if any).

    Example

    ```
    s = StateVarDict(age=10, sick=True)
    s['age'] += 1
    s.sick = False
    s.new_property = 'Wow !'
    ```

    `__init__`(*\*args*, *\*\*kwargs*)[source]¶
    :   Initialize self. See help(type(self)) for accurate signature.

#### emulsion.tools.timing module¶

A Python implementation of the EMuLSion framework (Epidemiologic
MUlti-Level SImulatiONs).

Tools for performance assessment.

`emulsion.tools.timing.``timethis`(*times=None*)[source]¶
:   A decorator function for printing execution time.

#### emulsion.tools.view module¶

A Python implementation of the EMuLSion framework (Epidemiologic
MUlti-Level SImulatiONs).

Tools for data / map visualization.

`emulsion.tools.view.``build_animation`(*values*, *title*, *unit=None*, *filename=None*, *cmap='coolwarm'*, *framerate=10*, *resolution=100*, *writer='imagemagick'*, *\*\*kwargs*)[source]¶
:   Create an animation based on the `values` list, with the
    specified title. A special color map name can be specified, as
    well as the presence of a colorbar and additional keyword
    arguments passed to `imshow`. If a specified unit is given, each
    frame is marked with its number and the corresponding unit. If a
    filename is provided in the `save` parameter, the animation is
    stored in that file.

`emulsion.tools.view.``show_contour`(*value*, *cmap='hot'*, *save=None*, *colbar=False*, *\*\*kwargs*)[source]¶
:   Display the specified value as a contour map with values. A
    special color map name can be specified, as well as the presence
    of a colorbar and additional keyword arguments passed to
    `contourf`. If a filename is provided in the `save` parameter, the
    image is stored in that file.

`emulsion.tools.view.``show_histo`(*value*, *xlabel*, *facecol='green'*, *save=None*)[source]¶
:   Display the distribution of the specified `value` array using
    `xlabel` for the legend. A specific face color can be used. If a
    filename is provided in the `save` parameter, the image is
    stored in that file.

`emulsion.tools.view.``show_img`(*value*, *cmap='hot'*, *save=None*, *colbar=True*, *\*\*kwargs*)[source]¶
:   Display the specified value as an image. A special color map
    name can be specified, as well as the presence of a colorbar and
    additional keyword arguments passed to `imshow`. If a filename is
    provided in the `save` parameter, the image is stored in that
    file.

### EMULSION

Epidemiological Multi-Level Simulation Framework

##### Navigation

- 1. Installation
- 2. Getting started with EMULSION
- 3. Modelling principles
- 4. Modelling language (basics)
- 5. Modelling language (advanced)
- 6. Feature examples
- 7. Information
- 8. License
- 9. High-level functions for model designers
- 10. emulsion package
  - 10.1. Subpackages
    - emulsion.agent package
    - emulsion.model package
    - emulsion.tools package
  - 10.2. Submodules
  - 10.3. emulsion.init\_emulsion module

##### Related Topics

- Documentation overview
  - 10. emulsion package
    - Previous: emulsion.model package

##### Quick search

©2016, INRA and Univ. Lille.
|
Powered by Sphinx 1.8.5
& Alabaster 0.7.10
|
Page source
