## Supplementary material for "EMULSION: transparent and flexible multiscale stochastic models in human, animal and plant epidemiology": S2 File: index.html

EMULSION Manual — EMULSION (Epidemiological Multi-Level Simulation framework)

### EMULSION Manual¶

Framework EMULSION is intended for modellers in epidemiology, to help
them design, simulate, and revise complex mechanistic stochastic
models, without having to write or rewrite huge amounts of code.

It comes with a *Domain-Specific Language* to represent all components
of epidemiological models (assumptions, model structure,
parameters…) in an explicit, intelligible and revisable way, and
thus facilitate interactions with other scientists (biologists,
veterinarians, economists…) throughout the modelling
process. EMULSION models are automatically processed by a modular
simulation engine, which, if needed, can also incorporate small code
add-ons for representing very specific features of a model
(Fig. 1).

Models can use classical modelling paradigms (compartments,
individual-based models, metapopulations) and multiple scales (from
individuals to metapopulations), thanks to recent research in
Artificial Intelligence (see Information).

Fig. 1 Principles of framework EMULSION

#### Table of contents¶

- 1. Installation
  - 1.1. Requirements
  - 1.2. Install with `pip` (recommended)
    - Linux and MacOS:
    - Windows
  - 1.3. Install third-party software
    - Linux
    - MacOS
    - Windows
  - 1.4. Test your installation
  - 1.5. Alternative: install with `git`
- 2. Getting started with EMULSION
  - 2.1. Running EMULSION
  - 2.2. Producing model diagrams
  - 2.3. Viewing parameters
  - 2.4. Changing parameters
  - 2.5. Changing the model
  - 2.6. Going further…
- 3. Modelling principles
  - 3.1. Individuals, populations, metapopulations
  - 3.2. From flow diagrams to state machines
    - Flow diagrams
    - State machines
- 4. Modelling language (basics)
  - 4.1. YAML Syntax in a nutshell
  - 4.2. Model structure
    - `model_name`
    - `model_info`
    - `time_info`
    - `state_machines`
    - `levels`
    - `grouping`
    - `processes`
    - `parameters`
    - `prototypes`
    - `initial_conditions`
    - `outputs`
    - `actions`
    - `statevars`
- 5. Modelling language (advanced)
  - 5.1. Compartments, IBM or hybrid models?
  - 5.2. Master state machines
    - Set states attributes
    - Customize transitions
    - Produce new individuals
    - State machines without transitions
  - 5.3. Design prototypes for typical individuals or populations
  - 5.4. Regulate time
  - 5.5. Complexify grouping
  - 5.6. Aggregate variables
  - 5.7. Automatic variables
  - 5.8. Built-in functions
  - 5.9. Built-in actions
  - 5.10. Changing scale: metapopulations
  - 5.11. Connecting to Python code add-ons
- 6. Feature examples
  - 6.1. SIR model
  - 6.2. SEIRS model
  - 6.3. SIRS model with periodic external risk
  - 6.4. Custom state durations
  - 6.5. SIR model with basic demography (births/deaths)
  - 6.6. SIR model with age groups
  - 6.7. SIR model with age groups and random initialization
  - 6.8. SIR model with cumulative incidence
  - 6.9. SIR model with individual actions and variable aggregation
  - 6.10. SIR model with age groups and explicit age
  - 6.11. SIR model with explicit gestation
  - 6.12. SIR model with structured population
  - 6.13. SIR model with metapopulation
  - 6.14. SIR model with metapopulation and data-driven movements
- 7. Information
  - 7.1. Contributors and contact
  - 7.2. How to cite
  - 7.3. Selected publications
  - 7.4. Acknowledgements
- 8. License
- 9. High-level functions for model designers
  - 9.1. Functions Available for Models
  - 9.2. Rates / probabilities
  - 9.3. Computations
  - 9.4. Selecting agents
  - 9.5. Durations
  - 9.6. Agent State and Variable Changes
  - 9.7. Introspection
- 10. emulsion package
  - 10.1. Subpackages
    - emulsion.agent package
    - emulsion.model package
    - emulsion.tools package
  - 10.2. Submodules
  - 10.3. emulsion.init\_emulsion module

##### Indices and tables¶

- Index
- Module Index
- Search Page

### EMULSION

Epidemiological Multi-Level Simulation Framework

##### Navigation

- 1. Installation
- 2. Getting started with EMULSION
- 3. Modelling principles
- 4. Modelling language (basics)
- 5. Modelling language (advanced)
- 6. Feature examples
- 7. Information
- 8. License
- 9. High-level functions for model designers
- 10. emulsion package

##### Related Topics

- Documentation overview
  - Next: 1. Installation

##### Quick search

©2016, INRA and Univ. Lille.
|
Powered by Sphinx 1.8.5
& Alabaster 0.7.10
|
Page source
