## Supplementary material for "EMULSION: transparent and flexible multiscale stochastic models in human, animal and plant epidemiology": S2 File: modules.html

emulsion — EMULSION (Epidemiological Multi-Level Simulation framework)

### emulsion¶

- 10. emulsion package
  - 10.1. Subpackages
    - emulsion.agent package
      - Subpackages
      - Submodules
      - emulsion.agent.action module
      - emulsion.agent.atoms module
      - emulsion.agent.comparts module
      - emulsion.agent.exceptions module
      - emulsion.agent.meta module
      - emulsion.agent.process module
      - emulsion.agent.views module
    - emulsion.model package
      - Submodules
      - emulsion.model.emulsion\_model module
      - emulsion.model.exceptions module
      - emulsion.model.functions module
      - emulsion.model.state\_machines module
    - emulsion.tools package
      - Submodules
      - emulsion.tools.calendar module
      - emulsion.tools.functions module
      - emulsion.tools.graph module
      - emulsion.tools.misc module
      - emulsion.tools.parallel module
      - emulsion.tools.plot module
      - emulsion.tools.simulation module
      - emulsion.tools.state module
      - emulsion.tools.timing module
      - emulsion.tools.view module
  - 10.2. Submodules
  - 10.3. emulsion.init\_emulsion module

### EMULSION

Epidemiological Multi-Level Simulation Framework

##### Navigation

- 1. Installation
- 2. Getting started with EMULSION
- 3. Modelling principles
- 4. Modelling language (basics)
- 5. Modelling language (advanced)
- 6. Feature examples
- 7. Information
- 8. License
- 9. High-level functions for model designers
- 10. emulsion package

##### Related Topics

- Documentation overview

##### Quick search

©2016, INRA and Univ. Lille.
|
Powered by Sphinx 1.8.5
& Alabaster 0.7.10
|
Page source
