## Supplementary material for "EMULSION: transparent and flexible multiscale stochastic models in human, animal and plant epidemiology": S2 File: LICENSE

                                 Apache License
                           Version 2.0, January 2004
                        http://www.apache.org/licenses/

   TERMS AND CONDITIONS FOR USE, REPRODUCTION, AND DISTRIBUTION

   1. Definitions.

      "License" shall mean the terms and conditions for use, reproduction,
      and distribution as defined by Sections 1 through 9 of this document.

      "Licensor" shall mean the copyright owner or entity authorized by
      the copyright owner that is granting the License.

      "Legal Entity" shall mean the union of the acting entity and all
      other entities that control, are controlled by, or are under common
      control with that entity. For the purposes of this definition,
      "control" means (i) the power, direct or indirect, to cause the
      direction or management of such entity, whether by contract or
      otherwise, or (ii) ownership of fifty percent (50%) or more of the
      outstanding shares, or (iii) beneficial ownership of such entity.

      "You" (or "Your") shall mean an individual or Legal Entity
      exercising permissions granted by this License.

      "Source" form shall mean the preferred form for making modifications,
      including but not limited to software source code, documentation
      source, and configuration files.

      "Object" form shall mean any form resulting from mechanical
      transformation or translation of a Source form, including but
      not limited to compiled object code, generated documentation,
      and conversions to other media types.

      "Work" shall mean the work of authorship, whether in Source or
      Object form, made available under the License, as indicated by a
      copyright notice that is included in or attached to the work
      (an example is provided in the Appendix below).

      "Derivative Works" shall mean any work, whether in Source or Object
      form, that is based on (or derived from) the Work and for which the
      editorial revisions, annotations, elaborations, or other modifications
      represent, as a whole, an original work of authorship. For the purposes
      of this License, Derivative Works shall not include works that remain
      separable from, or merely link (or bind by name) to the interfaces of,
      the Work and Derivative Works thereof.

      "Contribution" shall mean any work of authorship, including
      the original version of the Work and any modifications or additions
      to that Work or Derivative Works thereof, that is intentionally
      submitted to Licensor for inclusion in the Work by the copyright owner
      or by an individual or Legal Entity authorized to submit on behalf of
      the copyright owner. For the purposes of this definition, "submitted"
      means any form of electronic, verbal, or written communication sent
      to the Licensor or its representatives, including but not limited to
      communication on electronic mailing lists, source code control systems,
      and issue tracking systems that are managed by, or on behalf of, the
      Licensor for the purpose of discussing and improving the Work, but
      excluding communication that is conspicuously marked or otherwise
      designated in writing by the copyright owner as "Not a Contribution."

      "Contributor" shall mean Licensor and any individual or Legal Entity
      on behalf of whom a Contribution has been received by Licensor and
      subsequently incorporated within the Work.

   2. Grant of Copyright License. Subject to the terms and conditions of
      this License, each Contributor hereby grants to You a perpetual,
      worldwide, non-exclusive, no-charge, royalty-free, irrevocable
      copyright license to reproduce, prepare Derivative Works of,
      publicly display, publicly perform, sublicense, and distribute the
      Work and such Derivative Works in Source or Object form.

   3. Grant of Patent License. Subject to the terms and conditions of
      this License, each Contributor hereby grants to You a perpetual,
      worldwide, non-exclusive, no-charge, royalty-free, irrevocable
      (except as stated in this section) patent license to make, have made,
      use, offer to sell, sell, import, and otherwise transfer the Work,
      where such license applies only to those patent claims licensable
      by such Contributor that are necessarily infringed by their
      Contribution(s) alone or by combination of their Contribution(s)
      with the Work to which such Contribution(s) was submitted. If You
      institute patent litigation against any entity (including a
      cross-claim or counterclaim in a lawsuit) alleging that the Work
      or a Contribution incorporated within the Work constitutes direct
      or contributory patent infringement, then any patent licenses
      granted to You under this License for that Work shall terminate
      as of the date such litigation is filed.

   4. Redistribution. You may reproduce and distribute copies of the
      Work or Derivative Works thereof in any medium, with or without
      modifications, and in Source or Object form, provided that You
      meet the following conditions:

      (a) You must give any other recipients of the Work or
          Derivative Works a copy of this License; and

      (b) You must cause any modified files to carry prominent notices
          stating that You changed the files; and

      (c) You must retain, in the Source form of any Derivative Works
          that You distribute, all copyright, patent, trademark, and
          attribution notices from the Source form of the Work,
          excluding those notices that do not pertain to any part of
          the Derivative Works; and

      (d) If the Work includes a "NOTICE" text file as part of its
          distribution, then any Derivative Works that You distribute must
          include a readable copy of the attribution notices contained
          within such NOTICE file, excluding those notices that do not
          pertain to any part of the Derivative Works, in at least one
          of the following places: within a NOTICE text file distributed
          as part of the Derivative Works; within the Source form or
          documentation, if provided along with the Derivative Works; or,
          within a display generated by the Derivative Works, if and
          wherever such third-party notices normally appear. The contents
          of the NOTICE file are for informational purposes only and
          do not modify the License. You may add Your own attribution
          notices within Derivative Works that You distribute, alongside
          or as an addendum to the NOTICE text from the Work, provided
          that such additional attribution notices cannot be construed
          as modifying the License.

      You may add Your own copyright statement to Your modifications and
      may provide additional or different license terms and conditions
      for use, reproduction, or distribution of Your modifications, or
      for any such Derivative Works as a whole, provided Your use,
      reproduction, and distribution of the Work otherwise complies with
      the conditions stated in this License.

   5. Submission of Contributions. Unless You explicitly state otherwise,
      any Contribution intentionally submitted for inclusion in the Work
      by You to the Licensor shall be under the terms and conditions of
      this License, without any additional terms or conditions.
      Notwithstanding the above, nothing herein shall supersede or modify
      the terms of any separate license agreement you may have executed
      with Licensor regarding such Contributions.

   6. Trademarks. This License does not grant permission to use the trade
      names, trademarks, service marks, or product names of the Licensor,
      except as required for reasonable and customary use in describing the
      origin of the Work and reproducing the content of the NOTICE file.

   7. Disclaimer of Warranty. Unless required by applicable law or
      agreed to in writing, Licensor provides the Work (and each
      Contributor provides its Contributions) on an "AS IS" BASIS,
      WITHOUT WARRANTIES OR CONDITIONS OF ANY KIND, either express or
      implied, including, without limitation, any warranties or conditions
      of TITLE, NON-INFRINGEMENT, MERCHANTABILITY, or FITNESS FOR A
      PARTICULAR PURPOSE. You are solely responsible for determining the
      appropriateness of using or redistributing the Work and assume any
      risks associated with Your exercise of permissions under this License.

   8. Limitation of Liability. In no event and under no legal theory,
      whether in tort (including negligence), contract, or otherwise,
      unless required by applicable law (such as deliberate and grossly
      negligent acts) or agreed to in writing, shall any Contributor be
      liable to You for damages, including any direct, indirect, special,
      incidental, or consequential damages of any character arising as a
      result of this License or out of the use or inability to use the
      Work (including but not limited to damages for loss of goodwill,
      work stoppage, computer failure or malfunction, or any and all
      other commercial damages or losses), even if such Contributor
      has been advised of the possibility of such damages.

   9. Accepting Warranty or Additional Liability. While redistributing
      the Work or Derivative Works thereof, You may choose to offer,
      and charge a fee for, acceptance of support, warranty, indemnity,
      or other liability obligations and/or rights consistent with this
      License. However, in accepting such obligations, You may act only
      on Your own behalf and on Your sole responsibility, not on behalf
      of any other Contributor, and only if You agree to indemnify,
      defend, and hold each Contributor harmless for any liability
      incurred by, or claims asserted against, such Contributor by reason
      of your accepting any such warranty or additional liability.

   END OF TERMS AND CONDITIONS
