## Supplementary material for "EMULSION: transparent and flexible multiscale stochastic models in human, animal and plant epidemiology": S2 File: hybrid_gest.yaml

```
---

# EMULSION (Epidemiological Multi-Level Simulation framework)
# ===========================================================
# 
# Contributors and contact:
# -------------------------
# 
#     - Sébastien Picault
#     - Yu-Lin Huang
#     - Vianney Sicard
#     - Sandie Arnoux
#     - Gaël Beaunée
#     - Pauline Ezanno
# 
#     BIOEPAR, INRA, Oniris, Atlanpole La Chantrerie,
#     Nantes CS 44307 CEDEX, France
# 
# 
# How to cite:
# ------------
# 
#     S. Picault, Y.-L. Huang, V. Sicard, P. Ezanno (2017). "Enhancing
#     Sustainability of Complex Epidemiological Models through a Generic
#     Multilevel Agent-based Approach", in: C. Sierra (ed.), 26th
#     International Joint Conference on Artificial Intelligence (IJCAI),
#     AAAI, p. 374-380. DOI: 10.24963/ijcai.2017/53
# 
# 
# License:
# --------
# 
#    Copyright 2016 INRA and Univ. Lille
# 
#    Inter Deposit Digital Number: IDDN.FR.001.280043.000.R.P.2018.000.10000
# 
#    Agence pour la Protection des Programmes,
#    54 rue de Paradis, 75010 Paris, France
# 
#    Licensed under the Apache License, Version 2.0 (the "License");
#    you may not use this file except in compliance with the License.
#    You may obtain a copy of the License at
# 
#        http://www.apache.org/licenses/LICENSE-2.0
# 
#    Unless required by applicable law or agreed to in writing, software
#    distributed under the License is distributed on an "AS IS" BASIS,
#    WITHOUT WARRANTIES OR CONDITIONS OF ANY KIND, either express or implied.
#    See the License for the specific language governing permissions and
#    limitations under the License.

#  __  __           _      _
# |  \/  |         | |    | |
# | \  / | ___   __| | ___| |
# | |\/| |/ _ \ / _` |/ _ \ |
# | |  | | (_) | (_| |  __/ |
# |_|  |_|\___/ \__,_|\___|_|
#
#  _____        __                           _   _
# |_   _|      / _|                         | | (_)
#   | |  _ __ | |_ ___  _ __ _ __ ___   __ _| |_ _  ___  _ __
#   | | | '_ \|  _/ _ \| '__| '_ ` _ \ / _` | __| |/ _ \| '_ \
#  _| |_| | | | || (_) | |  | | | | | | (_| | |_| | (_) | | | |
# |_____|_| |_|_| \___/|_|  |_| |_| |_|\__,_|\__|_|\___/|_| |_|


# NAME OF THE MODEL
model_name: hybrid_gest


# DESCRIPTION OF THE MODEL
model_info:
  abstract: 'This model is a simple discrete-time, stochastic,
  hybrid model with explicit gestation and age. Individuals are also
  grouped automatically by parity, i.e. the number of successful
  gestations.
  '
  author: 'Sebastien Picault'


#  _______ _
# |__   __(_)
#    | |   _ _ __ ___   ___
#    | |  | | '_ ` _ \ / _ \
#    | |  | | | | | | |  __/
#    |_|  |_|_| |_| |_|\___|

# TIME INFORMATION
# This section defines the time unit (used e.g. for expressing
# durations, rates, etc.), the duration of the time step, the origin
# date and total duration of the simulation.
time_info:
  time_unit: 'days'
  delta_t: 1
  origin: 'January 1'
  total_duration: '200'


#  _                    _
# | |                  | |
# | |     _____   _____| |___
# | |    / _ \ \ / / _ \ / __|
# | |___|  __/\ V /  __/ \__ \
# |______\___| \_/ \___|_|___/

# ORGANIZATION LEVELS USED IN THE SIMULATION
# A hybrid model considers two levels: the population and
# its components (individuals)
levels:
  herd:
    desc: 'level of the population'
    aggregation_type: 'hybrid'
    contains:
      - animals
  animals:
    desc: 'level of the individuals'


#  _____
# |  __ \
# | |__) | __ ___   ___ ___  ___ ___  ___  ___
# |  ___/ '__/ _ \ / __/ _ \/ __/ __|/ _ \/ __|
# | |   | | | (_) | (_|  __/\__ \__ \  __/\__ \
# |_|   |_|  \___/ \___\___||___/___/\___||___/

# LIST OF PROCESSES (IN ORDER) TO BE EXECUTED DURING EACH TIME STEP
# In a hybrid model, processes take place at the population level
# (here, herd) and are managed by groupings
processes:
  herd:
    - aging
    - sex_repartition
    - parity_grouping


#   _____                       _
#  / ____|                     (_)
# | |  __ _ __ ___  _   _ _ __  _ _ __   __ _
# | | |_ | '__/ _ \| | | | '_ \| | '_ \ / _` |
# | |__| | | | (_) | |_| | |_) | | | | | (_| |
#  \_____|_|  \___/ \__,_| .__/|_|_| |_|\__, |
#                        | |             __/ |
#                        |_|            |___/

# DESCRIPTION OF GROUPINGS, ASSOCIATED WITH PROCESSES WHICH ARE NOT
# DIRECTLY BASED ON STATE MACHINES
# A grouping is based on one or more variables. For instance, the
# grouping for managing the 'infection' process is based on the
# variable 'health_state' (coming from the state machine with the same
# name): hence, one group will be defined for all possible values of
# the health_state variable.

grouping:
  herd:
    aging:
      machine_name: age_group
      key_variables: [age_group, sex]
    sex_repartition:
      machine_name: sex
      key_variables: [sex]
    parity_grouping:
      machine_name: parity
      key_variables: [parity]

#   _____ _        _         __  __            _     _
#  / ____| |      | |       |  \/  |          | |   (_)
# | (___ | |_ __ _| |_ ___  | \  / | __ _  ___| |__  _ _ __   ___  ___
#  \___ \| __/ _` | __/ _ \ | |\/| |/ _` |/ __| '_ \| | '_ \ / _ \/ __|
#  ____) | || (_| | ||  __/ | |  | | (_| | (__| | | | | | | |  __/\__ \
# |_____/ \__\__,_|\__\___| |_|  |_|\__,_|\___|_| |_|_|_| |_|\___||___/

# DESCRIPTION OF THE STATE MACHINES INVOLVED IN THE MODEL
state_machines:
  sex:
    desc: 'A state machine without transitions just for representing
    male/female animals
    '
    states:
      - Female:
          name: 'Female animals'
          desc: ''
          fillcolor: 'orange'
          default: yes
      - Male:
          name: 'Male animals'
          desc: ''
          fillcolor: 'deepskyblue'
  parity:
    desc: 'A state machine without transitions, just for representing
    the succession of relevant parities (number of successive
    farrowings). The order of states defines a predecessor/successor
    order used by keyword next_state in prototype next_parity
    '
    states:
      - P0:
          name: 'Nulliparous'
          desc: 'Nulliparous animal (no previous farrowing)'
          fillcolor: 'orange'
      - P1:
          name: 'Parity 1'
          desc: 'Animal with parity 1 (first farrowing)'
          fillcolor: 'pink'
      - P2:
          name: 'Parity 2'
          desc: 'Animal with parity 2'
          fillcolor: 'red'
      - P3:
          name: 'Parity 3+'
          desc: 'Animal with parity 3 and more'
          fillcolor: 'purple'
  age_group:
    desc: 'The state machine which defines the evolution of age
    groups.
    '
    states:
      - J:
          name: 'Juvenile'
          desc: 'juvenile animals'
          fillcolor: 'orange'
          on_stay:
            - set_var: age
              value: 'age + delta_t'
      - A:
          name: 'Non-gestating Adult'
          desc: 'adult animals'
          fillcolor: 'brown'
          on_stay:
            - set_var: age
              value: 'age + delta_t'
      - G:
          name: 'Gestating Adult'
          desc: 'adult animals (producing new juveniles)'
          fillcolor: 'deepskyblue'
          duration: dur_gestation
          on_stay:
            - set_var: age
              value: 'age + delta_t'
      - D:
          name: 'Dead'
          desc: 'compartment to put dead animals'
          fillcolor: white
          autoremove: yes
    transitions:
      - from: J
        to: A
        cond: 'GreaterThan(age, adult_age)'
        proba: 1
        desc: 'all Juveniles with age > adult_age move to
        Non-gestating Adults
        '
      - from: A
        to: D
        rate: '1 / avg_adult_death_age'
        desc: 'all Non-gestating Adults are subject to mortality due
        to old age
        '
      - from: A
        to: G
        cond: 'is_Female'
        rate: 'reproduction'
        desc: 'non-gestating adult females start gestation at rate
        "reproduction"
        '
      - from: G
        to: A
        proba: 1
        on_cross:
          - produce_offspring: 'newborn'
            amount: 1
          - become: 'next_parity'
        desc: 'at end of gestation, gestating adult females return to
        A state and produce offspring. Their parity is updated using
        the prototype next_parity which increments the value of
        discrete variable parity
        '
      - from: G
        to: D
        escape: yes
        rate: '1 / avg_adult_death_age'
        desc: 'all Gestating Adults are subject to mortality due
        to old age - this can occur even during gestation, hence
        escape is set to yes
        '

#  _____                               _
# |  __ \                             | |
# | |__) |_ _ _ __ __ _ _ __ ___   ___| |_ ___ _ __ ___
# |  ___/ _` | '__/ _` | '_ ` _ \ / _ \ __/ _ \ '__/ __|
# | |  | (_| | | | (_| | | | | | |  __/ ||  __/ |  \__ \
# |_|   \__,_|_|  \__,_|_| |_| |_|\___|\__\___|_|  |___/

# PARAMETERS/FUNCTIONS/EXPRESSIONS USED IN THE MODEL
parameters:
  initial_herd_size:
    desc: 'initial number of animals in the herd'
    value: 100
  initial_infected:
    desc: 'initial number of infectious animals in the herd'
    value: 1
  death:
    desc: 'the death rate (/day)'
    value: 0.01
  dur_gestation:
    desc: 'duration of gestation (days)'
    value: 40
  reproduction:
    desc: 'reproduction rate (/days)'
    value: '1/20'
  avg_adult_death_age:
    desc: 'mean longevity of adults (days)'
    value: 120
  adult_age:
    desc: 'age when individuals become adults (days)'
    value: 20
  init_age_distrib:
    desc: 'distribution of initial ages'
    value: 'adult_age + random_integers(0, 100)'

#   _____ _        _
#  / ____| |      | |
# | (___ | |_ __ _| |_ _____   ____ _ _ __ ___
#  \___ \| __/ _` | __/ _ \ \ / / _` | '__/ __|
#  ____) | || (_| | ||  __/\ V / (_| | |  \__ \
# |_____/ \__\__,_|\__\___| \_/ \__,_|_|  |___/


statevars:
  age:
    desc: 'age of the animal'


#  _____           _        _
# |  __ \         | |      | |
# | |__) | __ ___ | |_ ___ | |_ _   _ _ __   ___  ___
# |  ___/ '__/ _ \| __/ _ \| __| | | | '_ \ / _ \/ __|
# | |   | | | (_) | || (_) | |_| |_| | |_) |  __/\__ \
# |_|   |_|  \___/ \__\___/ \__|\__, | .__/ \___||___/
#                                __/ | |
#                               |___/|_|

# PROTOTYPES describe typical individuals, characterized by specific
# values of their variables
prototypes:
  animals:
    - initial:
        desc: 'initial animals'
        age_group: A
        age: init_age_distrib
        sex: random
        parity: random
    - newborn:
        desc: 'newly created animals'
        age_group: J
        sex: random
        age: 0
        parity: P0
    - next_parity:
        desc: 'increase the parity of a female at farrowing, based on
        the order in which parities are defined in the corresponding
        state machine
        '
        parity: next_state


#  _____       _ _   _       _
# |_   _|     (_) | (_)     | |
#   | |  _ __  _| |_ _  __ _| |
#   | | | '_ \| | __| |/ _` | |
#  _| |_| | | | | |_| | (_| | |
# |_____|_| |_|_|\__|_|\__,_|_|

#   _____                _ _ _   _
#  / ____|              | (_) | (_)
# | |     ___  _ __   __| |_| |_ _  ___  _ __  ___
# | |    / _ \| '_ \ / _` | | __| |/ _ \| '_ \/ __|
# | |___| (_) | | | | (_| | | |_| | (_) | | | \__ \
#  \_____\___/|_| |_|\__,_|_|\__|_|\___/|_| |_|___/

# INITIAL CONDITIONS FOR THE SIMULATION
# In IBM or hybrid models, initial conditions describe the prototypes
# involved with the amount of individuals in each of those prototypes
initial_conditions:
  herd:
    - prototype: initial
      amount: 'initial_herd_size'


#   ____        _               _
#  / __ \      | |             | |
# | |  | |_   _| |_ _ __  _   _| |_ ___
# | |  | | | | | __| '_ \| | | | __/ __|
# | |__| | |_| | |_| |_) | |_| | |_\__ \
#  \____/ \__,_|\__| .__/ \__,_|\__|___/
#                  | |
#                  |_|

# TYPE AND PERIODICITY OF OUTPUTS
# The amount of individuals in each state is automatically recorded
# for all state machines each time step during the simulation.
# Additional variables (e.g. expressions defined in the 'parameters'
# section) can be specified below (as 'extra_vars').
outputs:
  type: csv
  herd:
    period: 1
    extra_vars:
      - total_herd
      - total_G_Male
      - total_G_Female
...
```
