## Supplementary material for "EMULSION: transparent and flexible multiscale stochastic models in human, animal and plant epidemiology": S2 File: hybrid_SIR_metapop_data.py

```
# EMULSION (Epidemiological Multi-Level Simulation framework)
# ===========================================================
# 
# Contributors and contact:
# -------------------------
# 
#     - Sébastien Picault
#     - Yu-Lin Huang
#     - Vianney Sicard
#     - Sandie Arnoux
#     - Gaël Beaunée
#     - Pauline Ezanno
# 
#     BIOEPAR, INRA, Oniris, Atlanpole La Chantrerie,
#     Nantes CS 44307 CEDEX, France
# 
# 
# How to cite:
# ------------
# 
#     S. Picault, Y.-L. Huang, V. Sicard, P. Ezanno (2017). "Enhancing
#     Sustainability of Complex Epidemiological Models through a Generic
#     Multilevel Agent-based Approach", in: C. Sierra (ed.), 26th
#     International Joint Conference on Artificial Intelligence (IJCAI),
#     AAAI, p. 374-380. DOI: 10.24963/ijcai.2017/53
# 
# 
# License:
# --------
# 
#    Copyright 2016 INRA and Univ. Lille
# 
#    Inter Deposit Digital Number: IDDN.FR.001.280043.000.R.P.2018.000.10000
# 
#    Agence pour la Protection des Programmes,
#    54 rue de Paradis, 75010 Paris, France
# 
#    Licensed under the Apache License, Version 2.0 (the "License");
#    you may not use this file except in compliance with the License.
#    You may obtain a copy of the License at
# 
#        http://www.apache.org/licenses/LICENSE-2.0
# 
#    Unless required by applicable law or agreed to in writing, software
#    distributed under the License is distributed on an "AS IS" BASIS,
#    WITHOUT WARRANTIES OR CONDITIONS OF ANY KIND, either express or implied.
#    See the License for the specific language governing permissions and
#    limitations under the License.

"""

"""
from   pathlib               import Path
import csv
import dateutil.parser       as     dup
import numpy                 as     np
from emulsion.agent.managers import MetapopProcessManager

DATA_FILE = 'moves.csv'

#===============================================================
# CLASS Metapopulation (LEVEL 'metapop')
#===============================================================
class Metapopulation(MetapopProcessManager):
    """
    level of the metapop.

    """
    def initialize_level(self, **others):
        """Initialize an instance of Metapopulation.
        Additional initialization parameters can be introduced here if needed.
        """
        # read a CSV data file for moves:
        # date of movement, source herd, destination herd, age group, quantity

        # and restructure it according to origin_date and delta_t:
        # {step: {source_id: [(dest_id, age_group, qty), ...],
        #         ...},
        #  ...}
        origin = self.model.origin_date
        step_duration = self.model.step_duration
        moves = {}
        with open(Path(self.model.input_dir, DATA_FILE)) as csvfile:
            # read the CSV file
            csvreader = csv.DictReader(csvfile, delimiter=',')
            for row in csvreader:
                day = dup.parse(row['date'])
                if day < origin:
                    # ignore dates before origin_date
                    continue
                # convert dates into simulation steps
                step = (day - origin) // step_duration
                # group information by step and source herd
                if step not in moves:
                    moves[step] = {}
                src, dest, qty = int(row['source']), int(row['dest']), int(row['qty'])
                if src not in moves[step]:
                    moves[step][src] = []
                moves[step][src].append([dest, row['age'], qty])
        self.moves = moves

    #----------------------------------------------------------------
    # Processes
    #----------------------------------------------------------------
    def exchange_animals(self):
        """

        => INDICATE HERE HOW TO PERFORM PROCESS exchange_animals.
        """
        if self.statevars.step in self.moves:
            herds = self.get_populations()
            for source in self.moves[self.statevars.step]:
                for dest, age, qty in self.moves[self.statevars.step][source]:
                    # neither source/dest in simulated herds
                    if (source not in herds) and (dest not in herds):
                        # print('ignoring movement from {} to {} (outside the metapopulation)'.format(source, dest))
                        continue
                    # source not in simulated herds: create animal from prototype
                    if source not in herds:
                        # print('movement to {} coming from outside the metapopulation'.format(dest))

                        # retrieve prototype definition from the model
                        prototype = self.model.get_prototype(name='imported_movement', level='animals')
                        # change age group to comply with movement
                        prototype['age_group'] = self.get_model_value(age)
                        animals = [herds[dest].new_atom(custom_prototype=prototype)
                                   for _ in range(qty)]
                        # print('importing', animals)
                    else:
                        # find convenient animals
                        candidates = herds[source].select_atoms('age_group', age, process='aging')
                        # try to move the appropriate quantity
                        nb = min(len(candidates), qty)
                        animals = np.random.choice(candidates, nb, replace=False)
                        herds[source].remove_atoms(animals)
                        # print('removing', animals, 'from', source)
                        # update variable 'sold' in source herd
                        herds[source].statevars.sold += len(animals)
                    if dest not in herds:
                        pass
                        # print('movement from {} going outside the metapopulation'.format(source))
                    else:
                        herds[dest].add_atoms(animals)
                        # update variable 'purchased' in dest herd
                        herds[dest].statevars.purchased += len(animals)
                        # print('adding', animals, 'to', dest)
```
