## Supplementary material for "EMULSION: transparent and flexible multiscale stochastic models in human, animal and plant epidemiology": S2 File: IBM_SIR_JA_demo_random.yaml

```
---

# EMULSION (Epidemiological Multi-Level Simulation framework)
# ===========================================================
# 
# Contributors and contact:
# -------------------------
# 
#     - Sébastien Picault
#     - Yu-Lin Huang
#     - Vianney Sicard
#     - Sandie Arnoux
#     - Gaël Beaunée
#     - Pauline Ezanno
# 
#     BIOEPAR, INRA, Oniris, Atlanpole La Chantrerie,
#     Nantes CS 44307 CEDEX, France
# 
# 
# How to cite:
# ------------
# 
#     S. Picault, Y.-L. Huang, V. Sicard, P. Ezanno (2017). "Enhancing
#     Sustainability of Complex Epidemiological Models through a Generic
#     Multilevel Agent-based Approach", in: C. Sierra (ed.), 26th
#     International Joint Conference on Artificial Intelligence (IJCAI),
#     AAAI, p. 374-380. DOI: 10.24963/ijcai.2017/53
# 
# 
# License:
# --------
# 
#    Copyright 2016 INRA and Univ. Lille
# 
#    Inter Deposit Digital Number: IDDN.FR.001.280043.000.R.P.2018.000.10000
# 
#    Agence pour la Protection des Programmes,
#    54 rue de Paradis, 75010 Paris, France
# 
#    Licensed under the Apache License, Version 2.0 (the "License");
#    you may not use this file except in compliance with the License.
#    You may obtain a copy of the License at
# 
#        http://www.apache.org/licenses/LICENSE-2.0
# 
#    Unless required by applicable law or agreed to in writing, software
#    distributed under the License is distributed on an "AS IS" BASIS,
#    WITHOUT WARRANTIES OR CONDITIONS OF ANY KIND, either express or implied.
#    See the License for the specific language governing permissions and
#    limitations under the License.

#  __  __           _      _
# |  \/  |         | |    | |
# | \  / | ___   __| | ___| |
# | |\/| |/ _ \ / _` |/ _ \ |
# | |  | | (_) | (_| |  __/ |
# |_|  |_|\___/ \__,_|\___|_|
#
#  _____        __                           _   _
# |_   _|      / _|                         | | (_)
#   | |  _ __ | |_ ___  _ __ _ __ ___   __ _| |_ _  ___  _ __
#   | | | '_ \|  _/ _ \| '__| '_ ` _ \ / _` | __| |/ _ \| '_ \
#  _| |_| | | | || (_) | |  | | | | | | (_| | |_| | (_) | | | |
# |_____|_| |_|_| \___/|_|  |_| |_| |_|\__,_|\__|_|\___/|_| |_|


# NAME OF THE MODEL
model_name: IBM_SIR_JA_demo_random


# DESCRIPTION OF THE MODEL
model_info:
  abstract: 'This model is a simple discrete-time, stochastic,
  individual-based SIR model with explicit age groups and demography.
  Initialization uses random multinomial sampling of prototypes
  (between healthy and infectious) and of states (for age group).
  '
  author: 'Sebastien Picault'


#  _______ _
# |__   __(_)
#    | |   _ _ __ ___   ___
#    | |  | | '_ ` _ \ / _ \
#    | |  | | | | | | |  __/
#    |_|  |_|_| |_| |_|\___|

# TIME INFORMATION
# This section defines the time unit (used e.g. for expressing
# durations, rates, etc.), the duration of the time step, the origin
# date and total duration of the simulation.
time_info:
  time_unit: 'days'
  delta_t: 1
  origin: 'January 1'
  total_duration: '100'


#  _                    _
# | |                  | |
# | |     _____   _____| |___
# | |    / _ \ \ / / _ \ / __|
# | |___|  __/\ V /  __/ \__ \
# |______\___| \_/ \___|_|___/

# ORGANIZATION LEVELS USED IN THE SIMULATION
# An individual-based model considers two levels: the population and
# its components (individuals)
levels:
  population:
    desc: 'level of the population'
    aggregation_type: 'IBM'   # the keyword for designing
                              # invididual-based models
    contains:
      - individuals
  individuals:
    desc: 'level of the individuals'


#  _____
# |  __ \
# | |__) | __ ___   ___ ___  ___ ___  ___  ___
# |  ___/ '__/ _ \ / __/ _ \/ __/ __|/ _ \/ __|
# | |   | | | (_) | (_|  __/\__ \__ \  __/\__ \
# |_|   |_|  \___/ \___\___||___/___/\___||___/

# LIST OF PROCESSES (IN ORDER) TO BE EXECUTED DURING EACH TIME STEP In
# an Individual-Based Model, processes take place at the individuals
# level and are thus directly managed by state machines
processes:
  individuals:
    - health_state
    - age_group


#   _____                       _
#  / ____|                     (_)
# | |  __ _ __ ___  _   _ _ __  _ _ __   __ _
# | | |_ | '__/ _ \| | | | '_ \| | '_ \ / _` |
# | |__| | | | (_) | |_| | |_) | | | | | (_| |
#  \_____|_|  \___/ \__,_| .__/|_|_| |_|\__, |
#                        | |             __/ |
#                        |_|            |___/

# DESCRIPTION OF GROUPINGS, ASSOCIATED WITH PROCESSES WHICH ARE NOT
# DIRECTLY BASED ON STATE MACHINES
# Individual-Based Models do not use groupings


#   _____ _        _         __  __            _     _
#  / ____| |      | |       |  \/  |          | |   (_)
# | (___ | |_ __ _| |_ ___  | \  / | __ _  ___| |__  _ _ __   ___  ___
#  \___ \| __/ _` | __/ _ \ | |\/| |/ _` |/ __| '_ \| | '_ \ / _ \/ __|
#  ____) | || (_| | ||  __/ | |  | | (_| | (__| | | | | | | |  __/\__ \
# |_____/ \__\__,_|\__\___| |_|  |_|\__,_|\___|_| |_|_|_| |_|\___||___/

# DESCRIPTION OF THE STATE MACHINES INVOLVED IN THE MODEL
# Each state machine is composed at least of several states.
# The name of a state machine defines an individual variable which
# contains the value of the current state in the individual.
# Transitions define how each change can change to another one.
# Productions define how individuals in one state can produce
# new individuals in another state.
state_machines:
  # Below, the definition of the state machine used by the process
  # 'infection'
  health_state:
    desc: 'The state machine which defines the evolution of health
    states
    '
    # Below, the list of states with their attributes.
    states:
      - S:
          name: 'Susceptible'
          desc: 'suceptible of becoming infected'
          fillcolor: 'wheat'
          default: yes
      - I:
          name: 'Infectious'
          desc: 'infected and able to transmit the disease'
          fillcolor: 'maroon'
      - R:
          name: 'Resistant'
          desc: 'healthy again and resistant to infection'
          fillcolor: 'deepskyblue'
    transitions:
      - {from: S, to: I, rate: 'force_of_infection'}
      - {from: I, to: R, rate: 'recovery'}
  age_group:
    desc: 'The state machine which defines the evolution of age
    groups.
    '
    states:
      - J:
          name: 'Juvenile'
          desc: 'juvenile individuals'
          fillcolor: 'orange'
      - A:
          name: 'Adult'
          desc: 'adult individuals (producing new juveniles)'
          fillcolor: 'brown'
      - D:
          name: 'Dead'
          desc: 'compartment to put dead individuals'
          fillcolor: white
          autoremove: yes
    transitions:
      - {from: J, to: A, rate: 'maturation'}
      # death (assuming no disease-induced mortality)
      - {from: J, to: D, rate: 'death'}
      - {from: A, to: D, rate: 'death'}
    productions:
      # births (assuming no vertical transmission)
      - {from: A, to: J, rate: 'birth', prototype: 'newborn'}


#  _____                               _
# |  __ \                             | |
# | |__) |_ _ _ __ __ _ _ __ ___   ___| |_ ___ _ __ ___
# |  ___/ _` | '__/ _` | '_ ` _ \ / _ \ __/ _ \ '__/ __|
# | |  | (_| | | | (_| | | | | | |  __/ ||  __/ |  \__ \
# |_|   \__,_|_|  \__,_|_| |_| |_|\___|\__\___|_|  |___/

# PARAMETERS/FUNCTIONS/EXPRESSIONS USED IN THE MODEL
parameters:
  initial_population_size:
    desc: 'initial number of individuals in the population'
    value: 100
  initial_proba_infection:
    desc: 'probability that an individual of the population is infectious at
    t=0
    '
    value: 0.1
  transmission_I:
    desc: 'transmission rate from infectious individuals (/day)'
    value: 0.5
  recovery:
    desc: 'recovery rate (/day)'
    value: 0.1
  force_of_infection:
    desc: 'infection function'
    value: 'transmission_I * total_I / total_population'
    source: 'classical function assuming frequency dependence'
  percentage_prevalence:
    desc: 'proportion of infected+infectious individuals'
    value: '100 * total_I / total_population'
  birth:
    desc: 'the birth rate (/day)'
    value: 0.01
  death:
    desc: 'the death rate (/day)'
    value: 'birth'
  maturation:
    desc: 'rate at which juveniles become adults (/day)'
    value: '1/20'
  init_prop_juveniles:
    desc: 'average proportion of juveniles in the initial population'
    value: 0.75


#  _____           _        _
# |  __ \         | |      | |
# | |__) | __ ___ | |_ ___ | |_ _   _ _ __   ___  ___
# |  ___/ '__/ _ \| __/ _ \| __| | | | '_ \ / _ \/ __|
# | |   | | | (_) | || (_) | |_| |_| | |_) |  __/\__ \
# |_|   |_|  \___/ \__\___/ \__|\__, | .__/ \___||___/
#                                __/ | |
#                               |___/|_|

# PROTOTYPES describe typical individuals, characterized by specific
# values of their variables
prototypes:
  individuals:
    - healthy:
        desc: 'healthy individuals'
        health_state: S
        age_group: 'random(init_prop_juveniles, 1-init_prop_juveniles)'
    - infected:
        desc: 'infected individuals'
        health_state: I
        age_group: 'random(init_prop_juveniles, 1-init_prop_juveniles)'
    - newborn:
        desc: 'newly created individuals'
        health_state: default
        age_group: J


#  _____       _ _   _       _
# |_   _|     (_) | (_)     | |
#   | |  _ __  _| |_ _  __ _| |
#   | | | '_ \| | __| |/ _` | |
#  _| |_| | | | | |_| | (_| | |
# |_____|_| |_|_|\__|_|\__,_|_|

#   _____                _ _ _   _
#  / ____|              | (_) | (_)
# | |     ___  _ __   __| |_| |_ _  ___  _ __  ___
# | |    / _ \| '_ \ / _` | | __| |/ _ \| '_ \/ __|
# | |___| (_) | | | | (_| | | |_| | (_) | | | \__ \
#  \_____\___/|_| |_|\__,_|_|\__|_|\___/|_| |_|___/

# INITIAL CONDITIONS FOR THE SIMULATION
# In IBM or hybrid models, initial conditions describe the prototypes
# involved with the amount of individuals in each of those prototypes
initial_conditions:
  population:
    - prototype: [infected, healthy]
      # two possible prototypes for individual initially in the population
      amount: initial_population_size
      # drawn among a population of size initial_population_size
      proba: [initial_proba_infection, 1-initial_proba_infection]
      # with respective probabilities initial_proba_infection and
      # 1-initial_proba_infection


#   ____        _               _
#  / __ \      | |             | |
# | |  | |_   _| |_ _ __  _   _| |_ ___
# | |  | | | | | __| '_ \| | | | __/ __|
# | |__| | |_| | |_| |_) | |_| | |_\__ \
#  \____/ \__,_|\__| .__/ \__,_|\__|___/
#                  | |
#                  |_|

# TYPE AND PERIODICITY OF OUTPUTS
# The amount of individuals in each state is automatically recorded
# for all state machines each time step during the simulation.
# Additional variables (e.g. expressions defined in the 'parameters'
# section) can be specified below (as 'extra_vars').
outputs:
  type: csv
  population:
    period: 1
    extra_vars:
      - percentage_prevalence
      - total_population
...
```
