## Supplementary material for "EMULSION: transparent and flexible multiscale stochastic models in human, animal and plant epidemiology": S2 File: quickstart.yaml

```
---

# EMULSION (Epidemiological Multi-Level Simulation framework)
# ===========================================================
# 
# Contributors and contact:
# -------------------------
# 
#     - Sébastien Picault
#     - Yu-Lin Huang
#     - Vianney Sicard
#     - Sandie Arnoux
#     - Gaël Beaunée
#     - Pauline Ezanno
# 
#     BIOEPAR, INRA, Oniris, Atlanpole La Chantrerie,
#     Nantes CS 44307 CEDEX, France
# 
# 
# How to cite:
# ------------
# 
#     S. Picault, Y.-L. Huang, V. Sicard, P. Ezanno (2017). "Enhancing
#     Sustainability of Complex Epidemiological Models through a Generic
#     Multilevel Agent-based Approach", in: C. Sierra (ed.), 26th
#     International Joint Conference on Artificial Intelligence (IJCAI),
#     AAAI, p. 374-380. DOI: 10.24963/ijcai.2017/53
# 
# 
# License:
# --------
# 
#    Copyright 2016 INRA and Univ. Lille
# 
#    Inter Deposit Digital Number: IDDN.FR.001.280043.000.R.P.2018.000.10000
# 
#    Agence pour la Protection des Programmes,
#    54 rue de Paradis, 75010 Paris, France
# 
#    Licensed under the Apache License, Version 2.0 (the "License");
#    you may not use this file except in compliance with the License.
#    You may obtain a copy of the License at
# 
#        http://www.apache.org/licenses/LICENSE-2.0
# 
#    Unless required by applicable law or agreed to in writing, software
#    distributed under the License is distributed on an "AS IS" BASIS,
#    WITHOUT WARRANTIES OR CONDITIONS OF ANY KIND, either express or implied.
#    See the License for the specific language governing permissions and
#    limitations under the License.

#   _____                           _
#  / ____|                         | |
# | |  __  ___ _ __   ___ _ __ __ _| |
# | | |_ |/ _ \ '_ \ / _ \ '__/ _` | |
# | |__| |  __/ | | |  __/ | | (_| | |
#  \_____|\___|_| |_|\___|_|  \__,_|_|

#  _____        __                           _   _
# |_   _|      / _|                         | | (_)
#   | |  _ __ | |_ ___  _ __ _ __ ___   __ _| |_ _  ___  _ __
#   | | | '_ \|  _/ _ \| '__| '_ ` _ \ / _` | __| |/ _ \| '_ \
#  _| |_| | | | || (_) | |  | | | | | | (_| | |_| | (_) | | | |
# |_____|_| |_|_| \___/|_|  |_| |_| |_|\__,_|\__|_|\___/|_| |_|


# NAME OF THE DISEASE
model_name: Quickstart

model_info:
  abstract: 'This model is aimed at providing an overview of several EMULSION features.'

#  _______ _
# |__   __(_)
#    | |   _ _ __ ___   ___
#    | |  | | '_ ` _ \ / _ \
#    | |  | | | | | | |  __/
#    |_|  |_|_| |_| |_|\___|


# TIME INFORMATION
time_info:
  time_unit: days
  delta_t: 7
  origin: 'January 1'
  total_duration: '2*365'
  calendars:
    quarantines:
      period: {days: 365}
      events:
        quarantine_period: {begin: 'April 1', end: 'September 1'}

#  _                    _
# | |                  | |
# | |     _____   _____| |___
# | |    / _ \ \ / / _ \ / __|
# | |___|  __/\ V /  __/ \__ \
# |______\___| \_/ \___|_|___/


# LEVELS OF AGENTS
levels:
  individuals:
    desc: 'level of individuals'
  population:
    desc: 'level of the population'
    contains:
      - individuals
    aggregation_type: hybrid
    aggregate_vars:
      - name: nb_total_abortions
        collect: nb_abortions
        operator: 'sum'


#  _____
# |  __ \
# | |__) | __ ___   ___ ___  ___ ___  ___  ___
# |  ___/ '__/ _ \ / __/ _ \/ __/ __|/ _ \/ __|
# | |   | | | (_) | (_|  __/\__ \__ \  __/\__ \
# |_|   |_|  \___/ \___\___||___/___/\___||___/


# LIST OF PROCESSES (IN ORDER)
processes:
  population:
    - aging
    - lifecycle
    - infection


#   _____
#  / ____|
# | |  __ _ __ ___  _   _ _ __  ___
# | | |_ | '__/ _ \| | | | '_ \/ __|
# | |__| | | | (_) | |_| | |_) \__ \
#  \_____|_|  \___/ \__,_| .__/|___/
#                        | |
#                        |_|

# DESCRIPTION OF COMPARTMENTS ASSOCIATED WITH PROCESSES WHICH ARE NOT
# BASED ON METHODS:
grouping:
  population:
    infection:
      machine_name: health_state
      key_variables: [health_state]
    lifecycle:
      machine_name: life_cycle
      key_variables: [life_cycle, age_group]
    aging:
      machine_name: age_group
      key_variables: [age_group]

#   _____ _        _         __  __            _     _
#  / ____| |      | |       |  \/  |          | |   (_)
# | (___ | |_ __ _| |_ ___  | \  / | __ _  ___| |__  _ _ __   ___  ___
#  \___ \| __/ _` | __/ _ \ | |\/| |/ _` |/ __| '_ \| | '_ \ / _ \/ __|
#  ____) | || (_| | ||  __/ | |  | | (_| | (__| | | | | | | |  __/\__ \
# |_____/ \__\__,_|\__\___| |_|  |_|\__,_|\___|_| |_|_|_| |_|\___||___/

# DESCRIPTION OF THE STATE MACHINES INVOLVED IN THE MODEL
state_machines:
  age_group:
    desc: 'Age groups in the population'
    states:
      - J:
          name: 'Juvenile'
          desc: 'juvenile animals'
          fillcolor: 'orange'
          default: yes
          on_stay:
            - set_var: age
              value: 'age + delta_t'
      - A:
          name: 'Adult'
          desc: 'adult animals'
          fillcolor: 'brown'
          on_stay:
            - set_var: age
              value: 'age + delta_t'
    transitions:
      - {from: J, to: A, cond: 'GreaterThan(age, adult_age)', proba: 1}
  life_cycle:
    desc: 'Life cycle of animals regarding gestation'
    states:
      - G:
          name: 'Gestating'
          desc: 'state for gestating animals'
          fillcolor: 'green'
          duration: 'gestation_duration'
      - NG:
          name: 'NonGestating'
          desc: 'state for empty animals'
          fillcolor: 'cyan'
          duration: 'delay_before_new_gestation'
          default: yes
    transitions:
      - {from: NG, to: G, cond: is_A, rate: reproduction}
      - desc: 'normal farrowing at end of gestation'
        from: G
        to: NG
        amount-all-but: 0
        on_cross:
          - produce_offspring: [infected_newborn, protected_newborn]
            amount: 1
            proba:
              - 'OR(is_E, is_I, is_Q)*proba_vertical_transmission'
      - desc: 'abortion do tue infection'
        from: G
        to: NG
        # abortions can occur only if animal is infectious
        escape: is_I
        proba: daily_abortion
        on_cross:
          - set_var: nb_abortions
            value: 'nb_abortions + 1'


  health_state:
    desc: 'Evolution of health states'
    states:
      - M:
          name: 'Maternal protection'
          desc: 'animal protected by maternal antibodies'
          fillcolor: 'cyan'
          duration: 'adult_age'
      - S:
          name: 'Susceptible'
          desc: 'suceptible of becoming infectious'
          default: yes
          fillcolor: 'wheat'
      - E:
          name: 'Exposed'
          desc: 'already infected but not yet able to transmit the disease'
          fillcolor: 'lightcoral'
          duration: 'incubation_duration'
      - I:
          name: 'Infectious'
          desc: 'sick and able to transmit the disease'
          fillcolor: 'maroon'
      - R:
          name: 'Resistant'
          desc: 'healthy again and resistant to infection'
          fillcolor: 'deepskyblue'
      - Q:
          name: 'Quarantine'
          desc: 'sick animals that are put aside for a while'
          fillcolor: 'green'
          duration: 'quarantine_duration'
      - D:
          name: 'Dead'
          desc: 'dead animals'
          fillcolor: 'white'
          autoremove: yes
    transitions:
      - {from: M, to: S, rate: epsilon}
      - {from: S, to: E, rate: f}
      - {from: E, to: I, rate: alpha}
      - {from: I, to: R, rate: gamma}
      - {from: I, to: Q, rate: detection, when: 'quarantine_period'}
      - {from: R, to: S, rate: delta}
      - {from: Q, to: R, proba: 1}
      - {from: M, to: D, rate: mortality_healthy, escape: yes}
      - {from: S, to: D, rate: mortality_healthy}
      - {from: R, to: D, rate: mortality_healthy}
      - {from: E, to: D, rate: mortality_sick, escape: yes}
      - {from: I, to: D, rate: mortality_sick}
      - {from: Q, to: D, rate: mortality_sick, escape: yes}

#  _____                               _
# |  __ \                             | |
# | |__) |_ _ _ __ __ _ _ __ ___   ___| |_ ___ _ __ ___
# |  ___/ _` | '__/ _` | '_ ` _ \ / _ \ __/ _ \ '__/ __|
# | |  | (_| | | | (_| | | | | | |  __/ ||  __/ |  \__ \
# |_|   \__,_|_|  \__,_|_| |_| |_|\___|\__\___|_|  |___/

parameters:
  initial_population_size:
    desc: 'initial number of animals in the population'
    value: 100
  initial_infected:
    desc: 'initial number of infected animals in the population'
    value: 10
  adult_age:
    desc: 'minimal age required to produce offspring (days)'
    value: '365*2'
  init_age_distrib:
    desc: 'distribution of initial ages'
    value: 'randint(0, adult_age)'
  gestation_duration:
    desc: 'duration of the gestation (in days)'
    value: '30*8'
  delay_before_new_gestation:
    desc: 'minimum time to wait before new gestation can start'
    value: '30*3'
  reproduction:
    desc: 'reproduction rate of the animals'
    value: 0.05
  mortality_healthy:
    desc: 'mortality rate for healthy animals'
    value: 0.002
  mortality_sick:
    desc: 'mortality rate for sick animals (E / I / Q)'
    value: 0.004
  proba_vertical_transmission:
    desc: 'probability that the offspring of an infected animal (E/ I /Q) becomes Infectious'
    value: 0.3
  adult_age:
    desc: 'age when individuals become adults (days)'
    value: 30
  epsilon:
    desc: 'maternal antibodies loss rate'
    value: 0.04
  incubation_duration:
    desc: 'minimum duration of the exposed state'
    value: 10
  alpha:
    desc: 'rate for E->I transition'
    value: 0.3
  beta:
    desc: 'contamination rate per sick individual'
    value: 0.1
  gamma:
    desc: 'recovery rate'
    value: 0.01
  f:
    desc: 'force of infection'
    value: 'beta * total_I / total_population'
  delta:
    desc: 'resistance loss rate'
    value: 0.01
  quarantine_duration:
    desc: 'duration of the quarantine (days)'
    value: 30
  quarantine_size:
    desc: 'number of animals that can be put simultaneously in the quarantine area'
    value: 20
  detection:
    desc: 'rate at which animals are detected sick (falls to 0 if
    quarantine zone is full)
    '
    value: '0.01*room_in_quarantine'
  room_in_quarantine:
    desc: '1 if number of animals in quarantine < quarantine_size, 0
    otherwise
    '
    value: 'StrictLessThan(total_Q, quarantine_size)'
  percentage_prevalence:
    desc: '% of animals infected (E+I+Q)'
    value: '100 * (total_E + total_I + total_Q) / total_population'
  view_quarantine:
    desc: 'show quarantine period (*100 for visualization)'
    value: '100 * quarantine_period'
  global_abortion:
    desc: 'probability that an infected animal aborts during
    gestation
    '
    value: 0.2
  daily_abortion:
    desc: 'daily probability that an infected animal aborts during
    gestation
    '
    value: '1 - (1 - global_abortion)**(1/gestation_duration)'
#   _____ _        _
#  / ____| |      | |
# | (___ | |_ __ _| |_ _____   ____ _ _ __ ___
#  \___ \| __/ _` | __/ _ \ \ / / _` | '__/ __|
#  ____) | || (_| | ||  __/\ V / (_| | |  \__ \
# |_____/ \__\__,_|\__\___| \_/ \__,_|_|  |___/


statevars:
  age:
    desc: 'age of the animal'
  nb_abortions:
    desc: 'number of abortions experienced by the animal'


#  _____           _        _
# |  __ \         | |      | |
# | |__) | __ ___ | |_ ___ | |_ _   _ _ __   ___  ___
# |  ___/ '__/ _ \| __/ _ \| __| | | | '_ \ / _ \/ __|
# | |   | | | (_) | || (_) | |_| |_| | |_) |  __/\__ \
# |_|   |_|  \___/ \__\___/ \__|\__, | .__/ \___||___/
#                                __/ | |
#                               |___/|_|

prototypes:
  individuals:
    - protected_newborn:
        health_state: M
        life_cycle: NG
        age_group: J
        age: 0
        nb_abortions: 0
    - infected_newborn:
        health_state: I
        life_cycle: NG
        age_group: J
        age: 0
        nb_abortions: 0
    - default_healthy:
        health_state: S
        life_cycle: NG
        age: init_age_distrib
        age_group: J
        nb_abortions: 0
    - default_infected:
        health_state: I
        life_cycle: NG
        age: init_age_distrib
        age_group: J
        nb_abortions: 0

#  _____       _ _   _       _
# |_   _|     (_) | (_)     | |
#   | |  _ __  _| |_ _  __ _| |
#   | | | '_ \| | __| |/ _` | |
#  _| |_| | | | | |_| | (_| | |
# |_____|_| |_|_|\__|_|\__,_|_|

#                      _ _ _   _
#                     | (_) | (_)
#   ___ ___  _ __   __| |_| |_ _  ___  _ __  ___
#  / __/ _ \| '_ \ / _` | | __| |/ _ \| '_ \/ __|
# | (_| (_) | | | | (_| | | |_| | (_) | | | \__ \
#  \___\___/|_| |_|\__,_|_|\__|_|\___/|_| |_|___/

initial_conditions:
  population:
    - prototype: default_healthy
      amount: 'initial_population_size - initial_infected'
    - prototype: default_infected
      amount: 'initial_infected'


#   ____        _               _
#  / __ \      | |             | |
# | |  | |_   _| |_ _ __  _   _| |_ ___
# | |  | | | | | __| '_ \| | | | __/ __|
# | |__| | |_| | |_| |_) | |_| | |_\__ \
#  \____/ \__,_|\__| .__/ \__,_|\__|___/
#                  | |
#                  |_|

outputs:
  type: csv
  ## possible alternative: database
  # database_information:
  #   database: Quickstart
  #   dialect: mysql
  #   driver: mysqldb
  #   server_name: localhost # 10.0.26.250
  #   username: EMuLSion
  #   password: 'EMuLSion'
  #   # port: '' # 3306
  population:
    period: 1
    extra_vars:
      - total_population
      - percentage_prevalence
      - view_quarantine
      - nb_total_abortions
...
```
