## Supplementary material for "EMULSION: transparent and flexible multiscale stochastic models in human, animal and plant epidemiology": S2 File: NOTICE

=========================================================================
==  NOTICE file for use with the Apache License, Version 2.0,          ==
==         for the EMULSION 1.0 distribution.                          ==
=========================================================================

EMULSION (Epidemiological Multi-Level Simulation framework), version 1.0.

Copyright 2016 INRA and Univ. Lille

This product includes software developed at INRA, the French National
Institute for Agricultural Research (http://www.inra.fr)


Contributors and contact:
-------------------------

    - Sébastien Picault, INRA & Univ. Lille
    - Yu-Lin Huang, INRA
    - Vianney Sicard, INRA
    - Sandie Arnoux
    - Gaël Beaunée
    - Pauline Ezanno, INRA

    BIOEPAR, INRA, Oniris, 44307 Nantes, France


How to cite:
------------

    S. Picault, Y.-L. Huang, V. Sicard, P. Ezanno (2017). "Enhancing
    Sustainability of Complex Epidemiological Models through a Generic
    Multilevel Agent-based Approach", in: C. Sierra (ed.), 26th
    International Joint Conference on Artificial Intelligence (IJCAI),
    AAAI, p. 374-380. DOI: 10.24963/ijcai.2017/53
