## Supplementary material for "EMULSION: transparent and flexible multiscale stochastic models in human, animal and plant epidemiology": S2 File: README.md

EMULSION: Epidemiological Multi-Level Simulation framework
======================

![Version](https://img.shields.io/badge/version-1.0.10-f16152.svg)
![License](https://img.shields.io/badge/license-Apache--2.0-8cd0c3.svg)

<!-- markdown-toc start - Don't edit this section. Run M-x markdown-toc-refresh-toc -->
**Table of Contents**

- [EMULSION - Quick Start](#emulsion---quick-start)
    - [Part A - General information](#part-a---general-information)
    - [Part B - Getting Started](#part-b---getting-started)
    - [Part C - Content of the repository](#part-c---content-of-the-repository)

<!-- markdown-toc end -->


Part A - General information
----------------------------

Framework EMULSION is intended for modellers in epidemiology, to help
them design, simulate, and revise complex mechanistic stochastic
models, without having to write or rewrite huge amounts of code.

It comes with a Domain-Specific Language to represent all components
of epidemiological models (assumptions, model structure, parameters…)
in an explicit, intelligible and revisable way, and thus facilitate
interactions with other scientists (biologists, veterinarians,
economists…) throughout the modelling process. EMULSION models are
automatically processed by a modular simulation engine, which, if
needed, can also incorporate small code add-ons for representing very
specific features of a model.

Models can use classical modelling paradigms (compartments,
individual-based models, metapopulations) and multiple scales (from
individuals to metapopulations), thanks to recent research in
Artificial Intelligence.

- **Version 1.0.10**
- **License:** Apache-2.0
- **Contributors and contact:**
  - Sébastien Picault (``)
  - Yu-Lin Huang
  - Vianney Sicard
  - Sandie Arnoux
  - Gaël Beaunée
  - Pauline Ezanno (``)
- **How to cite:**
  S. Picault, Y.-L. Huang, V. Sicard, P. Ezanno (2017). "Enhancing
  Sustainability of Complex Epidemiological Models through a Generic
  Multilevel Agent-based Approach", in: C. Sierra (ed.), _26th
  International Joint Conference on Artificial Intelligence (IJCAI)_,
  AAAI, p. 374-380. DOI: `10.24963/ijcai.2017/53`

  BibTeX:

      @InProceedings{ijcai2017-emulsion,
        author =       {Picault, Sébastien and Huang, Yu-Lin and Sicard,
                        Vianney and Ezanno, Pauline},
        title =        {Enhancing Sustainability of Complex Epidemiological
                        Models through a Generic Multilevel Agent-based
                        Approach},
        booktitle =    {Proceedings of the 26th International Joint
                        Conference on Artificial Intelligence (IJCAI'2017)},
        year =         {2017},
        editor =       {Sierra, Carles},
        publisher =    {AAAI},
        pages =        {374--380},
        DOI =          {10.24963/ijcai.2017/53},
        ISBN =         {978-0-9992411-0-3},
      }

- **Acknowledgements** This research was funded by the French Research
 Agency (ANR) through projects MIHMES (ANR-10-BINF-07) and CADENCE
 (ANR-16-CE32-0007-01), the European fund for the Regional Development
 (FEDER) of Pays-de-la-Loire, and the Animal Health Division of
 INRA. This work was also supported by the SANT'Innov project of the
 For and On Regional Development (PSDR) French research programme,
 which is funded by INRA, the National Research Institute of Science
 and Technology for Environment and Agriculture (IRSTEA), and the
 French regions of Bretagne, Pays de la Loire, Normandie and Nouvelle
 Aquitaine.


Part B - Getting Started
---------------------

Please refer to [EMULSION on-line documentation](https://sourcesup.renater.fr/www/emulsion-public/).


Part C - Content of the repository
----------------------------------

  | File name           | Role                                                          |
  |---------------------|---------------------------------------------------------------|
  | `README.md`         | This file |
  | `LICENSE`         | The license |
  | `NOTICE`         | Complements to license |
  | `doc/`              | Location for technical report and web documentation |
  | `models/`           | Location for models (YAML and Python files) |
  | `models/X/outputs/` | Default location for outputs of model X |
  | `models/X/img/`     | Default location for image files generated by model X (plots, diagrams, maps...) |
  | `scripts/`       | Location for (shell/R) utility scripts and syntactic rules    |
  | `src/`       | Location of EMULSION source files  |


-----
_Last update: 2019-01-30, S. Picault (``)_
