## Supplementary figures and images for "EMULSION: transparent and flexible multiscale stochastic models in human, animal and plant epidemiology"

### compart_SEIRS_health_state_machine.png

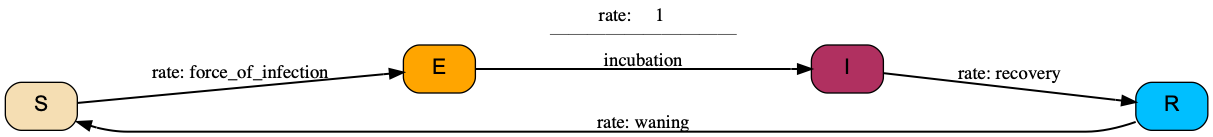

### compart_SIR_age_demo_health_state_machine.png

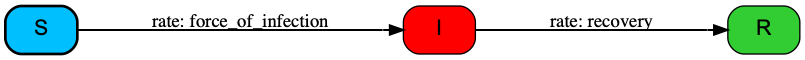

### compart_SIR_demo_health_state_machine.png

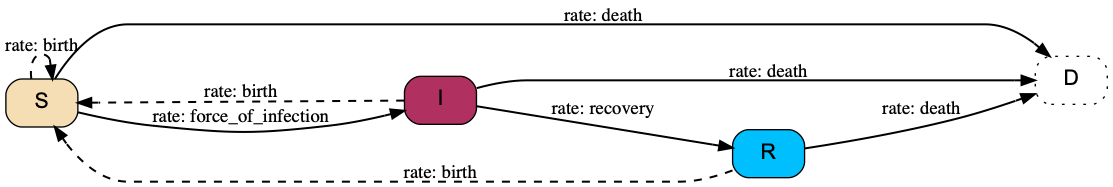

### compart_SIR_health_state_machine.pdf

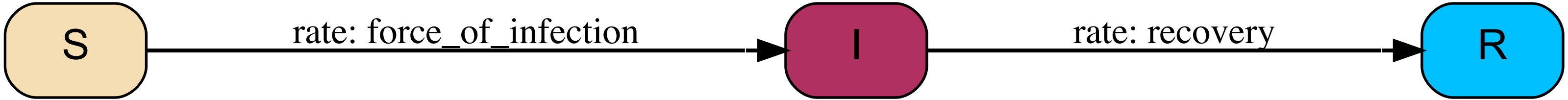

### compart_SIR_health_state_machine.png

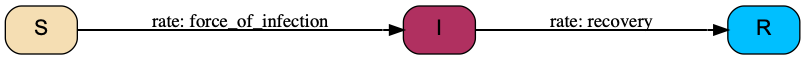

### compart_SIR_JA_demo_health_state_machine.png

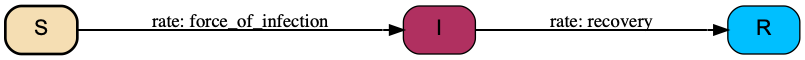

### compart_SIRS_periodic_risk_health_state_machine.png

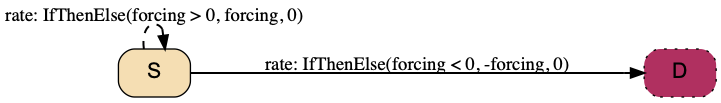

### hybrid_duration_age_group_machine.png

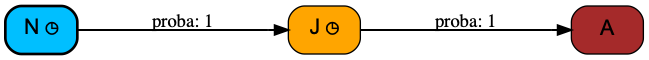

### hybrid_gest_age_group_machine.png

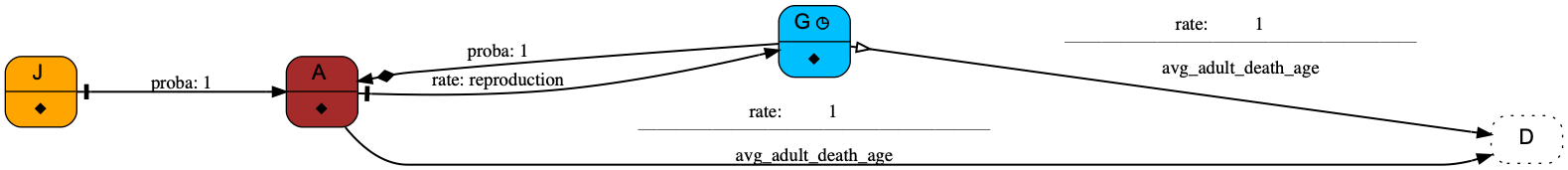

### hybrid_gest_sex_machine.png

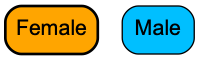

### hybrid_SEIR_aggreg_health_state_machine.png

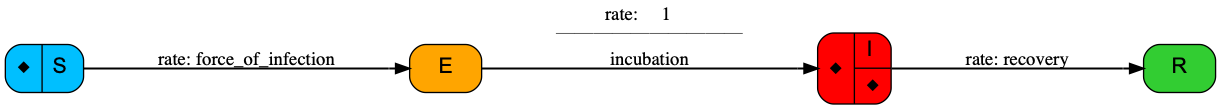

### hybrid_SIR_aggreg_health_state_machine.png

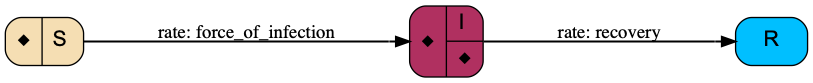

### hybrid_SIR_demo_health_state_machine.png

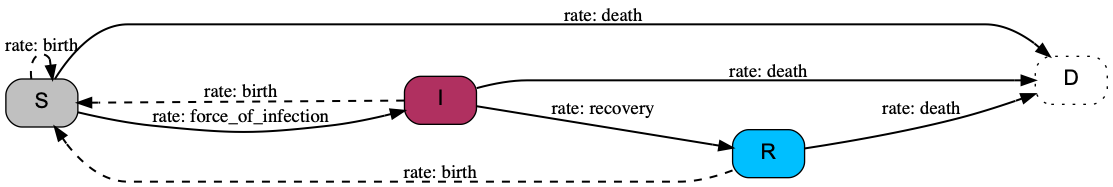

### hybrid_SIR_metapop_age_group_machine.png

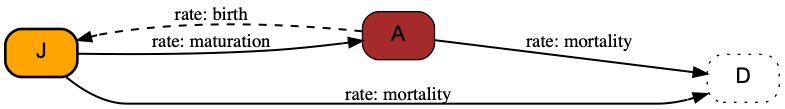

### hybrid_SIR_struct_age_group_machine.png

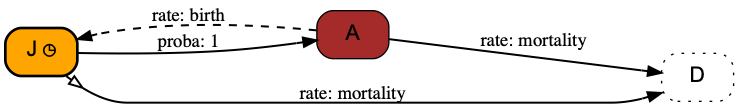

### hybrid_SIRS_aggreg_health_state_machine.png

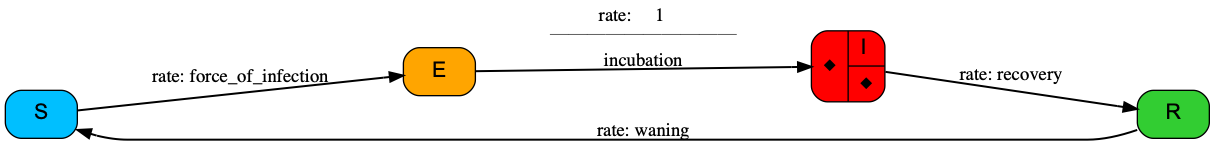

### IBM_duration_age_group_machine.png

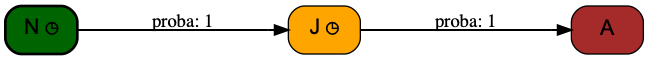

### IBM_gest_age_group_machine.png

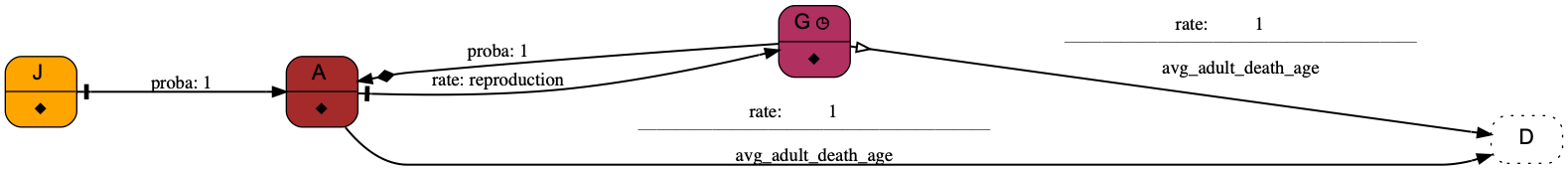

### IBM_gest_sex_machine.png

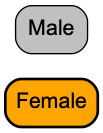

### IBM_SEIRS_health_state_machine.png

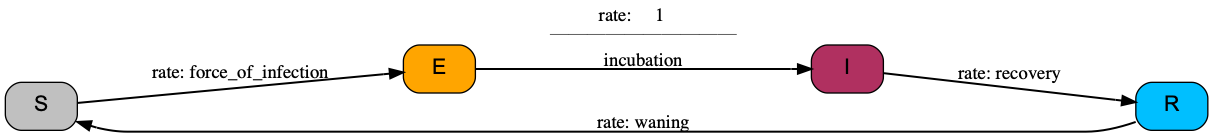

### IBM_SIR_age_demo_age_group_machine.png

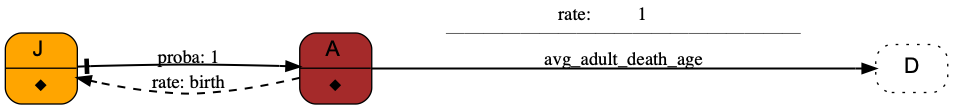

### IBM_SIR_age_demo_health_state_machine.png

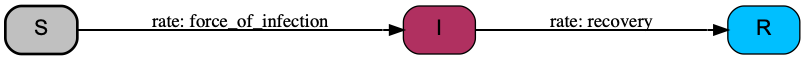

### IBM_SIR_aggreg_health_state_machine.png

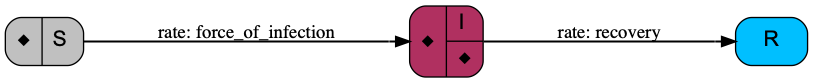

### IBM_SIR_health_state_machine.png

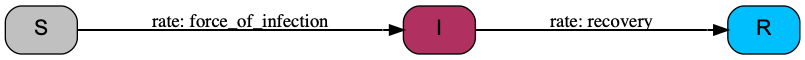

### IBM_SIR_JA_demo_age_group_machine.pdf

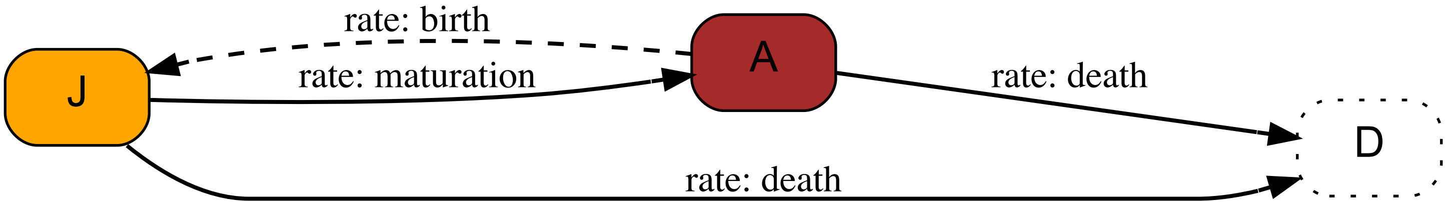

### IBM_SIR_JA_demo_age_group_machine.png

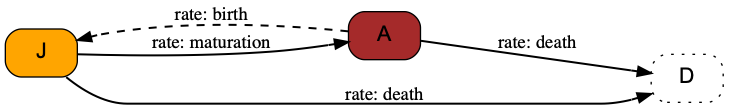

### IBM_SIR_JA_demo_health_state_machine.pdf

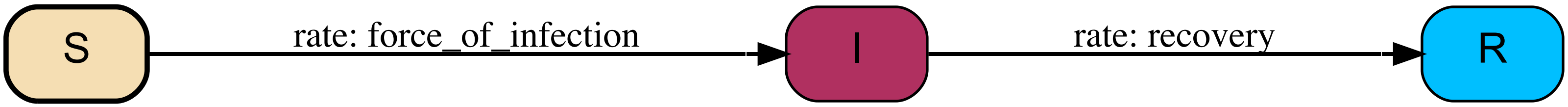

### IBM_SIR_metapop_age_group_machine.png

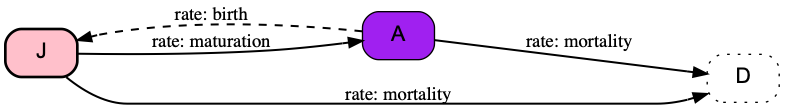

### IBM_SIRS_periodic_risk_health_state_machine.png

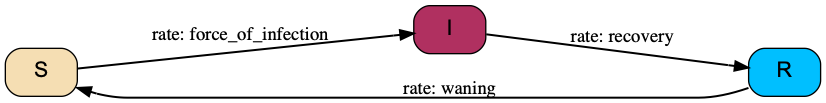

### Quickstart_age_group_machine.png

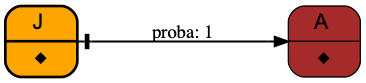

### Quickstart_health_state.png

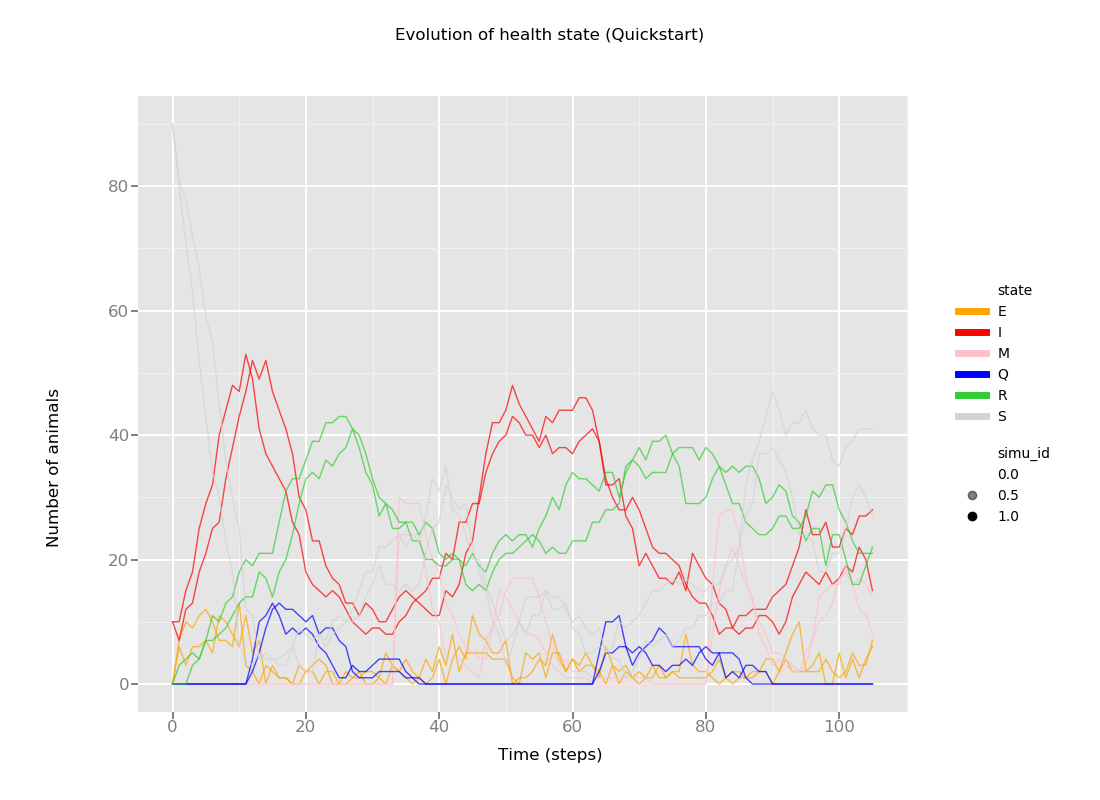

### Quickstart_health_state_machine.png

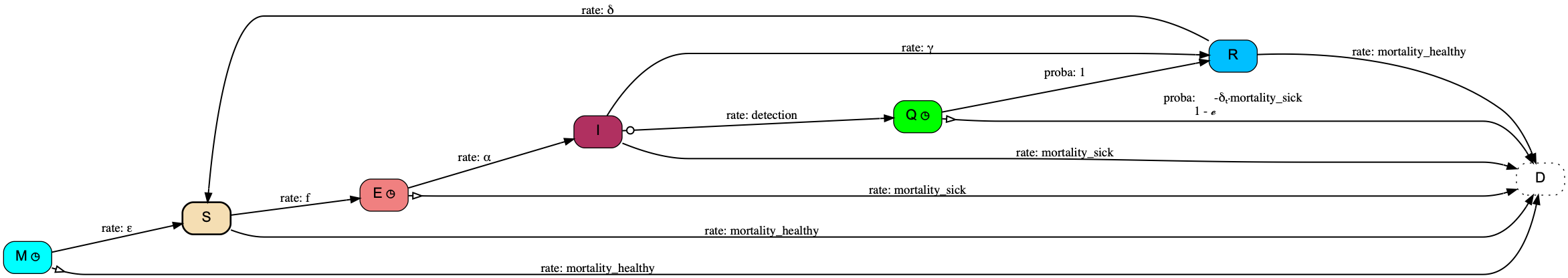

### Quickstart_life_cycle.png

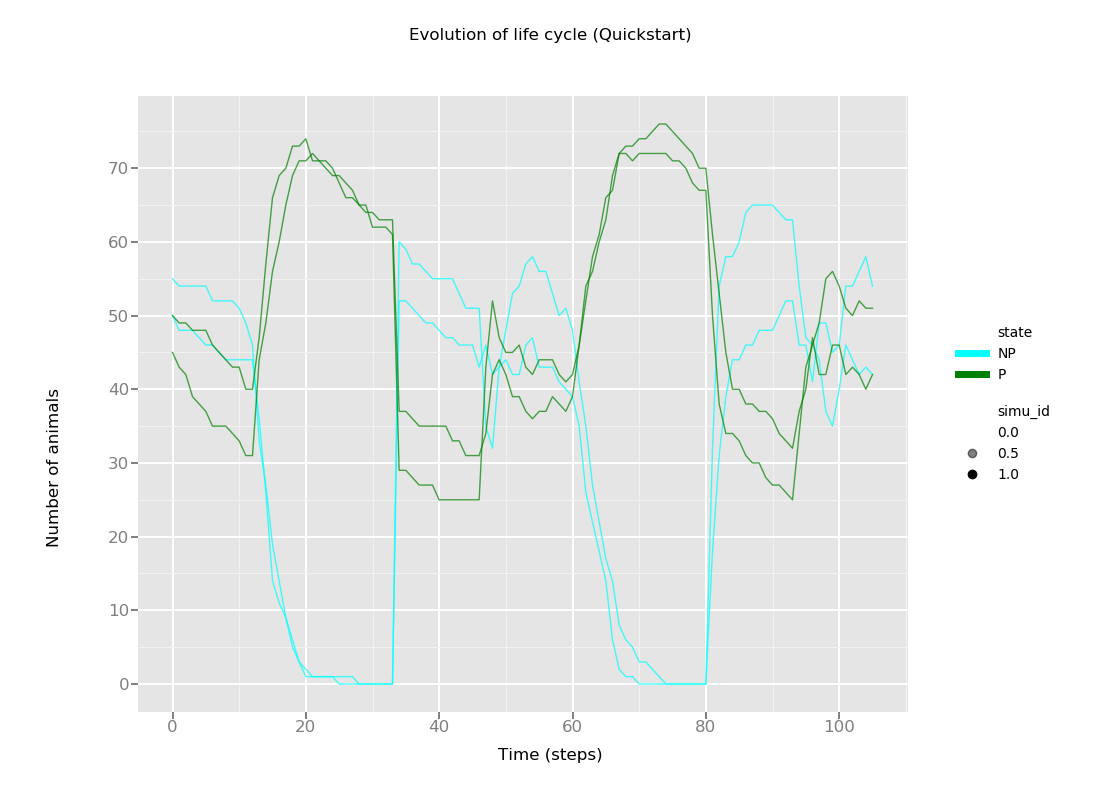

### Quickstart_life_cycle_machine.png

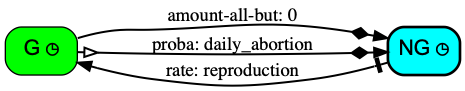

### stoch_compart_SIR_health_state_machine.png

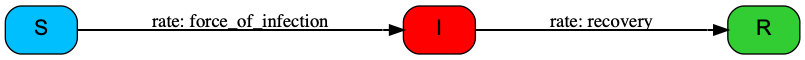

### VariablesAggregation_hybrid_age_group_machine.png

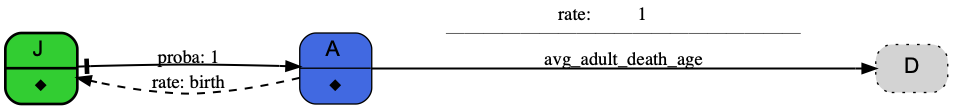

### VariablesAggregation_hybrid_health_state_machine.png

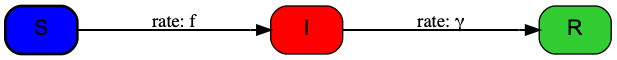
